# Pathological PNPase variants with altered RNA binding and degradation activity affect the phenotype of bacterial and human cell models

**DOI:** 10.1101/2024.10.03.616462

**Authors:** R. Pizzoccheri, F. A. Falchi, A. Alloni, M. Caldarulo, T. Camboni, F. Zambelli, G. Pavesi, C. Visentin, C. Camilloni, S. Sertic, F. Briani

**Affiliations:** Dipartimento di Bioscienze, Università degli Studi di Milano, Milano, 20133, Italy; Institute for Biomedical Technologies (ITB), National Research Council, Segrate, 20054, Italy

**Keywords:** PNPT1, polyribonucleotide phosphorylase, mitochondrial disease models, RNA binding proteins, RNA degradation

## Abstract

Human PNPase (hPNPase) is an essential RNA exonuclease located in mitochondria, where it contributes to RNA import from the cytoplasm, degradation of mitochondrial RNA, and R-loop homeostasis. Biallelic mutations in the hPNPase *PNPT1* gene cause different genetic diseases, ranging from hereditary hearing loss to Leigh syndrome. In this work, we used an *Escherichia coli* model we recently developed to test the effects of four pathological *PNPT1* mutations associated with diseases of different severity. Moreover, we generated a new human cell model by introducing *PNPT1* mutations into 293T cells via CRISPR-Cas editing. Notably, the bacterial cells expressing the different mutant alleles exhibited similar phenotypes consistent with hPNPase loss of function. In contrast, the human cell model responded differently to the two mutations tested, with responses correlating with the severity of the respective pathologies. We interpreted the data derived from both models in the light of the *in vitro* RNA binding and degradation activity of the purified wild-type and mutated hPNPase variants. We found that all pathogenic mutations tested caused defects in protein assembly and affected the degradation and RNA binding efficiency to varying degrees. However, the severity of the conditions caused by different mutations did not correlate with the catalytic activity of the mutant proteins. Also, we demonstrated that human PNPase, like its bacterial orthologue, is capable of trimming ssDNA and some mutations inhibit such activity.

## INTRODUCTION

Polyribonucleotide phosphorylase (PNPase) is a phosphorolytic exonuclease that degrades the RNA from the 3’- to the 5’-end (1). PNPase is widely conserved from bacteria to humans with some notable exceptions, as it is absent in Archaea and some unicellular eukaryotes (2–4).

Structural studies on PNPases of different organisms have shown that the protein is a homotrimer in which the protomers are assembled in a doughnut shape with the KH and S1 RNA binding domains on the top of a central channel where the catalytic site is located (3, 5–11). Relevant for RNA binding are also residues that forms a constriction point in the channel above the catalytic site (8, 10, 12, 13).

Human PNPase (hPNPase), encoded by the nuclear *PNPT1* gene (14), is a mitochondrial protein (15, 16) involved in different processes ranging from mitochondrial RNA (mtRNA) degradation and RNA import to mitochondrial DNA (mtDNA) homeostasis and resolution of R-loops, namely stable DNA-RNA hybrids that may impair DNA replication (reviewed in (4)). The protein is essential in mice and in human cells (15, 17–19) and its overexpression triggers reactive oxygen species (ROS) accumulation in Hela cells (20). Several missense *PNPT1* mutations cause genetic diseases with different symptoms and severity ranging from hereditary hearing loss to multisystem oxidative phosphorylation (OXPHOS) deficiency disorders, Leigh syndrome and spinocerebellar ataxia type 25 (21, 22). Understanding the molecular defects of pathological hPNPase variants in human cells is not an easy task given the plethora of processes involving hPNPase (4) and the heterogenous genetic background of patients, which prevents a clearcut comparison of the effects of different mutations.

In a previous work, with the aim to develop a simple model system for the functional characterization of hPNPase variants, we constructed an *Escherichia coli* strain expressing the hPNPase instead of the endogenous EcPNPase (23). Interestingly, hPNPase expression in *E. coli* caused phenotypes like increased ROS and R-loops levels that are related to hPNPase function in human cells. We hypothesized that the enzyme may retain the RNA binding, and not the RNA degrading activity in the bacterial cell context, in which phosphate concentration is optimal for EcPNPase but inhibitory for hPNPase (23, 24).

In this work, we have studied four pathological variants of hPNPase with missense mutations in different hPNPase domains (**Fig. 1A**) (25–28). In particular, the c.419C>T p. (P140L) mutation was identified in heterozygosity with c.407G > A p.(R136H) in a child with Leigh symptoms died at the age of 2 (27). In contrast, the c.1160A>G p.(Q387R) and c.1424A>G p.(E475G) mutations were found in homozygous patients, with the former causing severe neurological symptoms compatible with OXPHOS disease and the second determining adult-onset hearing loss (25, 26). Finally, the c.2234C>T p.(M745T) was found in two compound heterozygotes with a one-base insertion in the other allele causing a frame-shift mutation (c.1748dupA; p.(E584Gfs*17)). The patients (two siblings) had congenital hearing loss with a neurodegenerative course with ataxia, dystonia, and cognitive decline in the adulthood (28). In all cases, the heterozygous parents carrying a wild-type *PNPT1* allele were asymptomatic.

**Fig. 1.**
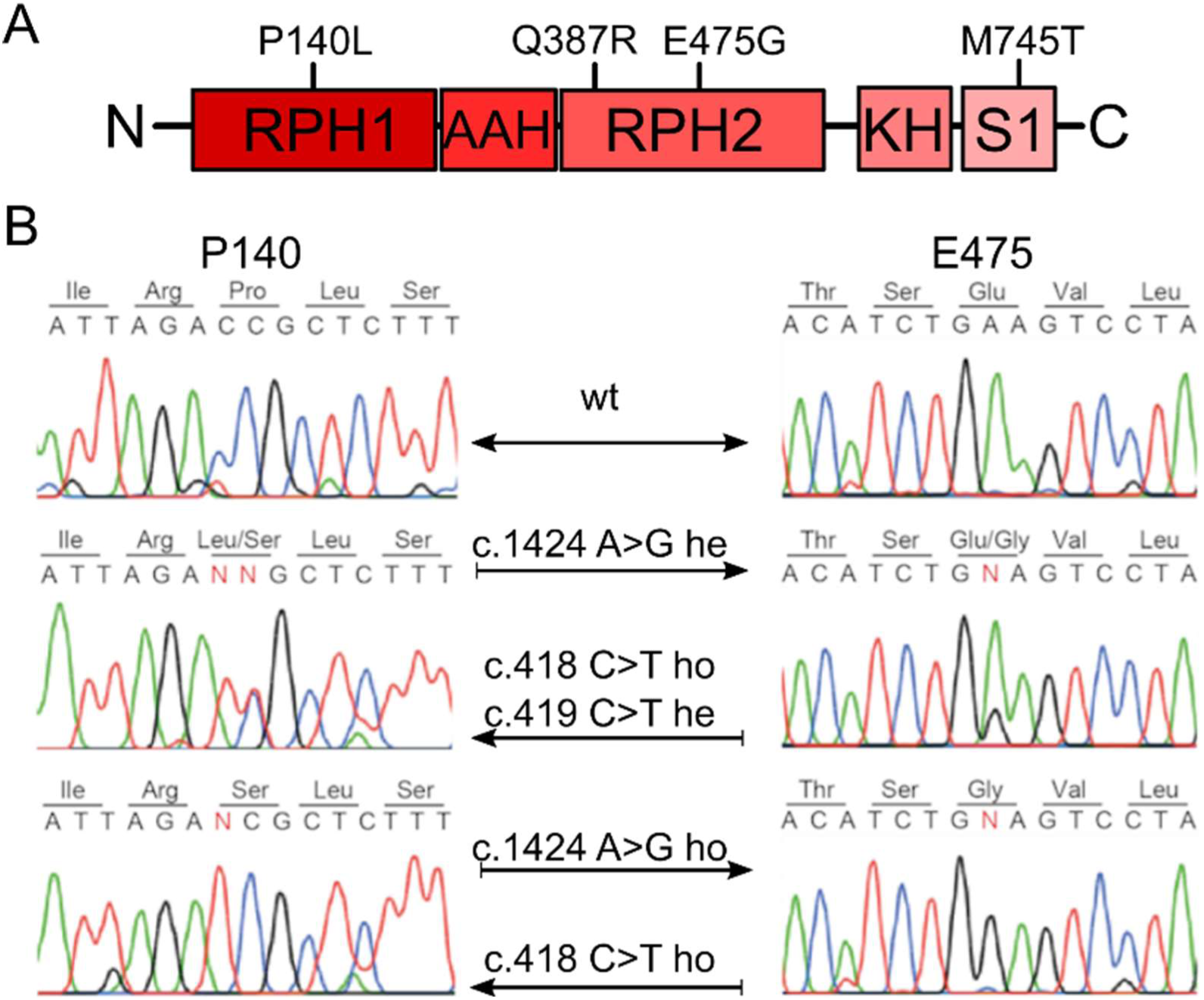
hPNPase domains and *PNPT1* mutations in 293T edited cells. A. RPH1 and RPH2, RNase PH domains 1 and 2; AAH, all α-helical domain; KH and S1, RNA binding domains. B. *PNPT1*^+^ mutations in edited 293T lines. Left panels. Sequence of the region of interest in exon 5 in *PNPT1*^+^ homozygous (wt, on top), c.418C>T / c.419C>T compound heterozygous (he, middle electropherogram) and homozygous c.418C>T lines (ho, on bottom). Right panels. Sequence of the region of interest in exon 17 in *PNPT1*^+^ homozygous (wt, on top), heterozygous (he) and homozygous (ho) lines for the c.1424A>G mutation (middle and bottom electropherograms, respectively). Nucleotide substitutions are represented with the red letter “N”. Aminoacidic sequence is shown with amino acid abbreviations above the nucleotide sequences.

We have analysed the effect of the four mutations *in vitro*, by testing the RNA binding and degradation activity of mutant proteins, and *in vivo*, in the *E. coli* model. Moreover, we introduced missense mutations in the P140 and E475 *PNPT1* codons of 293T human cells by CRISPR-Cas editing. Overall, our data suggest that the loss of phosphorolytic activity is not a major determinant of the severity of hPNPase-linked conditions and validate *E. coli* and 293T cells as useful models, which may be instrumental in the research of molecules restoring hPNPase activity.

## MATERIALS AND METHODS

### Bacterial strains, Human cell lines and plasmids

*Escherichia coli* strains, human cell lines, plasmids and oligonucleotides used in this work are listed in **Supplementary Table S1**. Genomic coordinates of *E. coli* genes refer to MG1655 reference genome (GenBank Accession Number U00096.3). Strains expressing *PNPT1*-mutant variants, namely C-6027, C-6028, C-6029, and C-6030, are C-1a (29) derivatives obtained by replacing the *pnp* ORF (region 3309033–3311168) with *PNPT1_Ec_*:kanR cassettes by λRED recombination and removing the kanR cassette by FLP-mediated recombination (30). *PNPT1_Ec_* encodes hPNPase with *E. coli* codon usage (23). The cassettes were prepared as described (23) by combining amplicons obtained by PCR with primers specific for either the ends of *PNPT1*_Ec_ open reading frame (ORF) on pET28b-hPNP^P140L^, pET28b-hPNP^Q387R^, pET28b-hPNP^E475G^ and pET28b-hPNP^M745T^ plasmid DNA, or the kanR cassette with flanking FLP sites on pKD13 (30).

pET28b-hPNP^P140L^, pET28b-hPNP^Q387R^, pET28b-hPNP^E475G^, pET28b-hPNP^M745T^ were constructed by site directed mutagenesis using Q5® Site-Directed Mutagenesis Kit (New England Biolabs) with ad hoc primers. pET29-hPNP was constructed by PCR amplification of *PNPT1_Ec_* on pET28-hPNP. The amplicon was cloned in pET29b (+) digested with *Nde*I-*Xho*I, by NEBuilder Hi-Fi DNA Assembly (New England Biolabs), in frame with a C-terminal 6x His-tag. Plasmids expressing sgRNA for Base Editor-dCas9 mediated editing were generated using Q5® Site-Directed Mutagenesis Kit (New England Biolabs), using the following primers: pFYF1320_P140La/b; pFYF1320_E475Ga/b; pFYF1320_P140L/Sa/b; pFYF1320-L141La/b.

Bacterial cultures were grown aerobically in LD broth or M9 supplemented-medium (23), with the addition of 100 μg/mL ampicillin, 30 μg/mL chloramphenicol, 30 μg/mL kanamycin, and *25* or 100 μg/mL streptomycin, when needed. Human **c**ells were cultured in complete DMEM High Glucose GlutaMAX (GibcoTM) supplemented with 10% heat inactivated-fetal bovine serum, 0.5% Sodium Pyruvate (Lonza™ BioWhittaker), 1% Penicillin/Streptomycin (Gibco™), and 0,1% Plasmocin® prophylactic (InvivoGen) (hereafter DF10 medium). When needed, media were supplemented with 50 µg/mL uridine. Cell cultures were grown at 37 °C in a humified atmosphere with 5% CO_2_.

### Protein expression and purification

Cultures of *E. coli* strain SHuffle® T7 (New England Biolabs) carrying the pET vector expressing hPNPase variants fused to N- or C-terminal 6x His tag were grown in 1L of LD with 30 μg/ml kanamycin to OD_600_ = 1.0 at 37 °C. Induction was performed with 0.5 mM IPTG and further incubation 20 h at 18 °C. Bacterial pellets were lysed by two passages through a cell disruptor (One Shot model; Constant Systems Ltd.) in buffer A (50 mM Tris-HCl pH 8.0, 300 mM NaCl, 5 mM MgCl_2_, 1 mM EDTA, 10% glycerol), supplemented with DNase I (0.2 mg/g cells) RNase A (0.2 mg/g cells), 1 mM phenylmethanesulfonylfluoride (PMSF) and 1x Halt™ Protease Inhibitor Cocktail (Thermo Scientific). Crude cell extracts were passed through a Ni-NTA column (Nuvia™, Bio-Rad). The columns were washed with 10 column volumes (CV) of 4% buffer B (50 mM Tris HCl, pH 8.0, 300 mM NaCl, 500 mM imidazole, 10% glycerol) in buffer A. Proteins were eluted by a 10%, 20%, 50%, 70%, and 100% buffer B stepwise gradient, at 1 CV per step.

Samples were applied to a gel filtration column (Superdex 200 Increase 10*/*300 GL, GE Healthcare) and eluted with 50 mM Tris-HCl, pH 8.0, 150 mM NaCl and 10% glycerol. Purified protein samples were stored at −80 °C until use. Oligomerization state of the purified proteins was checked by Blue-Native gel electrophoresis (31) on 4-16% gradient polyacrylamide gel (NativePAGE™ Bis-Tris gel system; Invitrogen).

### Electrophoretic mobility shift and Degradation assays

Probes for electrophoretic mobility shift assay (EMSA) and *in vitro* degradation were prepared as follows. ^32^P-radiolabelled RNA and DNA probes were obtained by radiolabeling at the 5’-end 10 pmol of a 20-mer RNA oligonucleotide (RNA20) and a 20-mer ssDNA oligonucleotide (DNA20; (32)) by incubation with [γ-^32^P] ATP and T4 polynucleotide kinase. For fluorescent COX RNA probe production, a DNA fragment containing the T7 promoter controlling transcription of the 100 bp at the 3’ end of the human mtRNA COX1 was transcribed using the High Yield T7 Atto488 RNA Labeling Kit (UTP-based) (Jena Biosciences). All probes were purified by phenol-chloroform extraction, ethanol precipitated in the presence of 1 mg/ml glycogen and resuspended in 30 μl of RNase-free water. EMSA was performed as described (32) by incubating the probe (0.6 nM RNA20 or DNA20 or 0.4 nM Atto488-COX) for 20 min at 21 °C in Binding buffer (50 mM Tris HCl pH 7.5, 50 mM NaCl, 0.5 mM DTT, 0.025% NP40, 10% glycerol) with purified hPNPase in a final volume of 10 μl. The samples were fractionated by native 5% PAGE at 4 °C. For *in vitro* degradation experiments, the probe (2 nM radiolabelled RNA20 or DNA20 and 0.3 nM Atto488-COX) was incubated with 50 nM hPNPase in 10 mM Tris HCl, pH 7.5, 10 mM KCl, 2 mM MnCl_2_, 0.75 mM DTT, 2% PEG-6000, 0.5 mM Pi and either 2 mM MnCl_2_ (for DNA) or 2 mM MgCl_2_ (for RNA) at 37°C in a final volume of 25 μl. 5 µl samples were withdrawn at different time points and transferred in 10 µl of RNA loading dye (2 mg/ml xylene cyanol and bromophenol blue, 10 mM EDTA in formamide) to stop the reaction. The samples were fractionated by denaturing 10% PAGE. Images of radioactive gels were acquired by phosphoroimaging with Typhoon™ FLA 7000 (GE Healthcare), whereas fluorescent gels were acquired with ChemiDoc MP Imaging System (Bio-Rad) using the DyLight488 channel. Signals were quantified with Image Lab (Bio-Rad) software.

### Western blotting

Protein extracts were obtained from mid-exponential *E. coli* cultures grown aerobically at 37 °C as described (23). hPNPase stability in *E. coli* was measured by adding 25 µg/mL streptomycin to cultures at OD_600_ = 0.6. 2 ml samples were collected at 20 min intervals and centrifuged to pellet bacteria. The pellets were resuspended in SDS Buffer (50 mM Tris HCl pH 6.8, 2% SDS, 10% glycerol, 100 mM β-mercaptoethanol, 0.1 % bromophenol blue) at final OD_600_ = 25, boiled 3 min to lyse the bacteria and directly analysed by SDS-PAGE. Total protein extracts were prepared from adherent human cells pre-washed with PBS by incubating the cells for 30 minutes in ice with RIPA buffer (50 mM Tris HCl pH 8.0, 150 mM NaCl, 1% NP-40, 0.5% Sodium Deoxycholate, 0.1% SDS, 1mM EDTA, 10 mM NaF) supplemented with 1 mM PMSF and 1x Halt™ Protease Inhibitor Cocktail (ThermoScientific™) and vortexing every 10 minutes. Samples were clarified by centrifugation at 16000 r.c.f. for 15 minutes at 4 °C. Total proteins were quantified using either the Pierce™ Detergent Compatible Bradford Assay Kit (ThermoFisher Scientific) or DC Protein Assay (Bio-Rad), following the manufacturer’s instructions. Protein samples for denaturing gel loading were prepared in SDS Buffer, boiled for 10 minutes at 95 °C, and analysed by 10 or 12% SDS-PAGE. Gels were blotted onto nitrocellulose membranes with the Trans-Blot Turbo Transfer System (Bio-Rad). Membranes were stained with Ponceau S solution (Sigma) and immunodetection was performed with the following antibodies: monoclonal anti-hPNPase (Anti-hPNP: PNPase (G-11) sc-365049; Santa Cruz Biotechnology); polyclonal anti-hPNPase (PA5-22397; Invitrogen); monoclonal anti-β2-actin (sc-8432; Santa Cruz Biotechnology); monoclonal anti-GAPDH (sc-47724; Santa Cruz Biotechnology); polyclonal anti-S3 ribosomal protein (33).

### ROS and R-loop determination

ROS and R-loop levels were determined in *E. coli* as previously described (23). In brief, for ROS determination, 200 µL of exponential cultures grown at 37 °C up to OD_600_ = 0.8 were harvested and resuspended in 200 µL of 1x PBS containing 1.0 mM of DCFH-DA (Sigma-Aldrich).

Samples were incubated 30 min at 37 °C, washed 3x with 1x PBS and transferred to a 96-well plate. OD_600_ and fluorescence at 488/530 nm were measured with the Ensight Plate Reader (PerkinElmer). Fluorescence values were normalized to the OD_600_ of the tested sample. R-loops were quantified on 10 μg of total nucleic acids extracted from overnight cultures grown at 37 °C in LB, spotted on Hybond-N^+^ membrane (PerkinElmer) using a Dot-blot apparatus (Hybri-Dot Manifold; BRL) and UV-crosslinked. The membrane was blocked with 5% milk solution in TBS supplemented with 0.1% Tween-20 (TBS-T) and incubated overnight at 4 °C with the anti-DNA: RNA hybrid antibody (MABE1095, clone S9.6, EDM Millipore) diluted 1:5000 in blocking buffer. After incubation with a secondary anti-mouse IgG antibody (Thermo Fisher Scientific) for 1h at room temperature, the signals were detected with PDS standard ECL GeneSpin kit using the ChemiDoc Touch Imaging System (Bio-Rad). After S9.6 signal detection, the membrane was washed, and nucleic acids were stained with 0.05% methylene blue in 0.5 M sodium acetate buffer (pH 5.2).

### Real-Time quantitative PCR (RT-qPCR)

RNA was extracted from *E. coli* cultures as described (23). 1 µg of total RNA was treated with RQ1 RNase-free DNase (Promega) and retrotranscribed using the iScript™ cDNA Synthesis Kit (Bio-Rad). Total human RNA was extracted with the RNeasy Mini Kit (QIAGEN) from adherent cells grown at 80-90% confluency in 35 mm plates and washed with 1x PBS before extraction. 1 µg of RNA was treated with RQ1 RNA-free DNase and retrotranscribed using either LunaScript® RT Supermix (New England Biolabs) or iScript™ cDNA Synthesis Kit (Bio-Rad). qPCR was performed by mixing the bacterial cDNA dilutions with primers specific for *recA* mRNA (recA-a and recA-b) or 16S rRNA (16Sa and 16Sb), and the human cDNA dilutions with primers specific for *PNTP1* (PNTP1a and PNTP1b); β-actin (ACTa and ACTb) or *MT-CO1* (CO1a and CO1b) in iTaq Universal SYBR® Green Super Mix (Bio-Rad). Two technical duplicates were performed for each biological replicate and reactions were run in a CFX Connect Real-Time PCR Detection System (Bio-Rad). 16S rRNA and β-actin mRNAs were used as reference genes to normalize the results relative to the expression of bacterial and human genes, respectively, to calculate the relative fold change in gene expression using the CFXmaestro Software (Bio-Rad).

### 293T transfection and genomic DNA preparation

293T cells were seeded into 48- or 6-well plates and transfected respectively with 2 µL or 7.5 µL of Lipofectamine 3000 (ThermoFisher Scientific) using 750 ng or 3 µg of either ABE-GFP or BE4-GFP base editor-expressing plasmid, 250 ng or 1 µg of sgRNA-expressing plasmid, and 20 ng or 0.1 µg of mCherry-expressing plasmid. Cells were cultured for 72 hours and single cell-sorted for both GFP and mCherry expression using a FACSAria II (BD Biosciences) into 96-well plates. Clonal expansion was monitored, and wells with single clones were identified under an optical microscope. Once clones reached almost full confluency into the 96-well plate, they were transferred by trypsinisation to progressively larger wells and grown until complete confluency in two 35 mm plates. Cells were harvested for both genomic DNA (gDNA) extraction and cryo-conservation. gDNA was extracted using the Monarch Genomic DNA Purification Kit (NewEngland Biolabs®) following the manufacturer’s instructions. Exon 5 and exon 17 of *PNPT1* were amplified by PCR using GoTaq® Green Master Mix (Promega) and the primers 5a - 5b for exon 5 and 17a - 17b for exon 17. PCR amplicons were purified using NZYGelpure kit (NZY) and processed for Sanger sequencing (Eurofins). DNA sequences were compared to the Ensembl reference sequence ENST00000447944.7.

### Cell proliferation and viability assays

Cell proliferation was determined with the CellTiter 96 AQueous One Solution Cell Proliferation Assay (Promega) according to the manufacturer’s instructions. Briefly, 24-hours following seeding in 96 well plates, 20 µL of MTS reagent was added to 100 µL of DF10 in which cells were seeded, and the plate was incubated for 2 hours at 37 °C into an humified incubator. Bioreduction of the MTS compound into soluble and coloured formazan in metabolically active cells was recorded as a change in absorbance at 490 nm using the Ensight Plate Reader (PerkinElmer). Cell viability was assessed performing trypan-blue exclusion test. Briefly, 20 µL of cell suspension was mixed with 20 µL of 0.4% V/W trypan blue and the number of viable cells detected using the TC20™ Automated Cell Counter (Bio-Rad).

### Mitochondrial DNA copy number determination

Changes in mtDNA over nuclear DNA content were measured as described (34) by Real-time qPCR. qPCR was performed by mixing 6 ng of total DNA with primers for mt-tRNA(Leu (UUR)) (UURa and UURb) or for β2-microglobulin nuclear gene (GLa and GLb) in iTaq Universal SYBR Green Supermix (Bio-Rad). Three technical replicates were performed for each biological replicate and reactions were run in a CFX Connect Real-Time PCR Detection System (Bio-Rad). Calculation of the mtDNA content relative to the genomic DNA was performed using the following equations: ΔCt = (nDNA Ct − mtDNA Ct); Relative mitochondrial DNA content = 2×2ΔCt.

### Mitochondrial stress test

2×10^4^ cells were seeded into poly-D-lysine-coated XF24-well plates with DF10 for 8-16 hours before the assay. DF10 medium was then replaced with the Seahorse XF DMEM medium (Agilent), and the cells were incubated one hour at 37 °C without supplied CO_2_. Agilent Seahorse XF Cell Mito Stress Test Kit (Agilent) was used to assess mitochondrial function with a Seahorse XFe24 Analyzer according to the manufacturer’s protocol and using the following concentration of each compound: 1.5 µM oligomycin, 2 µM carbonyl cyanide 4-(trifluoromethoxy)phenylhydrazone (FCCP), 0.5 µM rotenone/antimycin A. The results were normalized over the number of cells detected using the SMARTer ICELL8 cx Single-Cell System, following cell staining with Hoechst 33342 (ThermoScientific).

### RNA-Seq analysis of global transcription of human cell lines

8.5×10^5^ cells were seeded into a 6-well plate and, the day after, RNA was isolated from cultures incubated or not with 1 µM staurosporine for 6 hours to induce apoptosis using the RNeasy Mini Kit (QIAGEN). RNA samples were checked for quality on Agilent 4200 TapeStation System (Agilent Technologies) and quantified with a Qubit fluorometer (ThermoFisher Scientific). RNA-seq libraries were prepared using 200 ng of the RNA and the Illumina TruSeq Stranded Total RNA Prep, Ligation with Ribo-Zero Plus (Illumina), according to manufacturer’s instructions. Then, libraries were normalized, pooled and sequenced on an Illumina NextSeq 1000 platform with 2×100bp paired-end runs yielding ∼33-73M reads per sample. Sequence reads were mapped with STAR v2.7.1 against the RefSeq gene annotation (2023-05-29), excluding model transcripts (XM_ and XR_) of the human hs1 (T2T) genome retrieved from the UCSC Genome Browser Database (35). Read counts and subsequent normalized transcripts per million (TPM) were computed with RSEM v1.3.1 (rsem-calculate-expression), on the same RefSeq gene annotation. Differential expression analysis was performed with edgeR (36). Raw expected counts from RSEM were normalized for library composition across all samples by trimmed mean of M values (TMM) after removing non-expressed genes, with default parameters. Differentially expressed genes (DEGs) in pairwise comparisons were identified by the exact test function of edgeR (exactTest) using a Benjamini–Hochberg corrected pvalue threshold of 0.01. Enrichment analyses were performed with Enrichr (37).

### Statistical analysis

The replicate number for each sample and the statistical test applied are indicated in the figure legends, when appropriate. The differences in means were considered statistically significant at p < 0.05.

## RESULTS

### Disease-linked mutations affect hPNPase migration in native gel

We purified from *E. coli* His-tagged hPNPase variants devoid of the mitochondrial localization signal and tested their migration in denaturing and native gel. All mutant variants had electrophoretic mobility comparable to the wild-type (wt) in SDS-PAGE with the exception of M745T, which migrated slightly faster (**FIG. 2A**). This “gel-shifting” phenomenon of proteins differing for a single amino acid was previously described (38, 39).

The fusion of an His-tag at the N-terminal end of purified hPNPase was previously related with the formation of dimeric forms of hPNPase (6). In contrast, we observed in Blue Native (BN) gels, that the wt hPNPase with the His-tag at either the N- or the C-terminal formed a complex compatible with a homotrimer (predicted size, approximately 270 kDa *vs*. 234 kDa of the *E. coli* PNPase homotrimer) (**Fig. 2B**). On the contrary, all mutant proteins except M745T had altered migration compared to the wt protein (**Fig. 2C**). In detail, P140L hPNPase homotrimer was barely detectable and other forms were not visible suggesting that this mutant protein may precipitate and/or form aggregates not entering the gel. The Q387R and E475G proteins formed a smeared band at around 250-240 kDa and a fainter one between 146 and 242 kDa molecular weight (MW) markers, implying that the homotrimers formed by these variants tend to dissociate during the electrophoretic run and dimers are formed, consistent with results reported by others (6, 26).

**Fig. 2.**
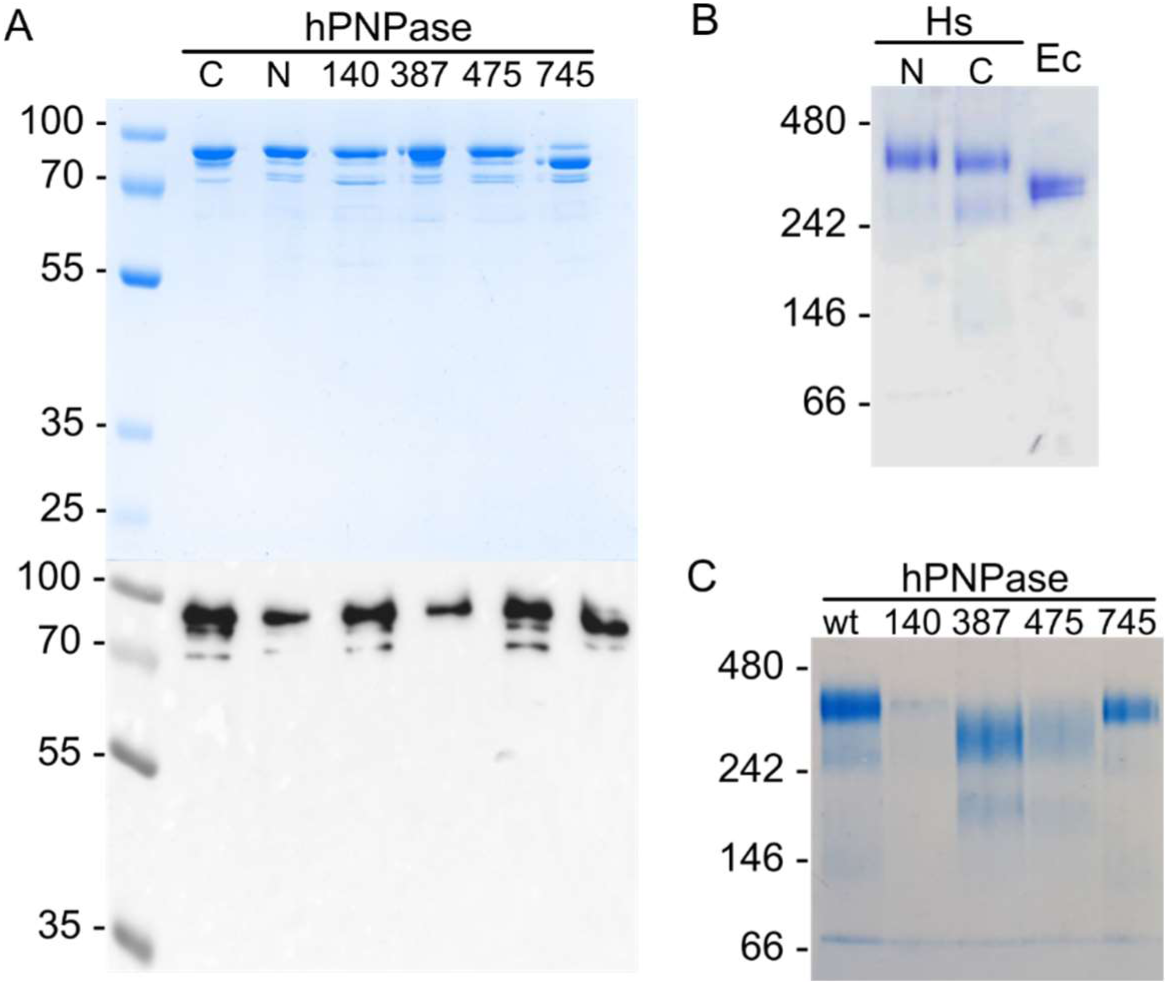
Migration in denaturing and native gels of hPNPase variants. A. Commassie staining (upper panel) and western blot (lower panel) of purified hPNPase variants. 4 or 0.8 µg of purified hPNPase variants run in 10 % SDS-PAGE were analysed either by gel-staining with Coomassie or by western blotting with monoclonal anti-hPNPase antibodies, respectively. The migration of molecular weight (MW; kDa) markers (Thermo Scientific PageRuler Plus Prestained Ladder) is reported on the left. hPNPase pathological variants are indicated with the position of the mutated amino acid. N and C indicate the wt hPNPase with the His tag at the N- or C-terminal, respectively. B. Migration of EcPNPase (Ec) and hPNPase with His-tag at either the N-ter (N) or the C-ter (C) in 4-16% Blue Native (BN) gel. 2 µg of purified proteins were loaded. C. Migration of wt and mutated hPNPase variants in 4-16% BN gel. 4 µg of purified proteins were loaded. hPNPase pathological variants are indicated as in A. In B and C, the migration of MW (kDa) markers (NativeMark Unstained Protein Standard, Invitrogen) is reported on the left.

### *PNPT1* pathological mutations impinge on the RNA binding activity of hPNPase

We investigated the RNA binding and degradation activities of purified hPNPase mutant proteins. For these experiments, we employed a 100 nt long heteropolymeric RNA probe corresponding to the 100 nt at the 3’-end of *MT-CO1* (mitochondrially encoded cytochrome C oxidase I) mRNA (COX RNA) (**Supplementary Fig. S1**). In electrophoretic mobility shift assays (EMSA), we observed that the unbound COX signal disappeared when the wt hPNPase or the P140L variant were at a 25- to 62-fold molar excess relative to the probe, and at a 62- to 125-fold molar excess for the other mutants. A complex (designated as complex 1) with significantly reduced mobility relative to free COX RNA was visible upon incubation of COX RNA with the wt hPNPase and at high concentrations of the M745T mutant protein. This complex was absent or barely detectable when the probe was incubated with the other variants (**Fig. 3A and Supplementary Figs. S2A-B**). Complexes with higher mobility than complex 1 (designated as complex 2 and 3) were detected with all mutated hPNPases and occasionally also with the wt (**Fig. 3A and Supplementary Fig. S2A**), consistent with the formation of complexes involving dimeric/monomeric forms of hPNPase. Thus, the wt and mutated hPNPases bind COX with comparable efficiency but form different complexes.

**Fig. 3.**
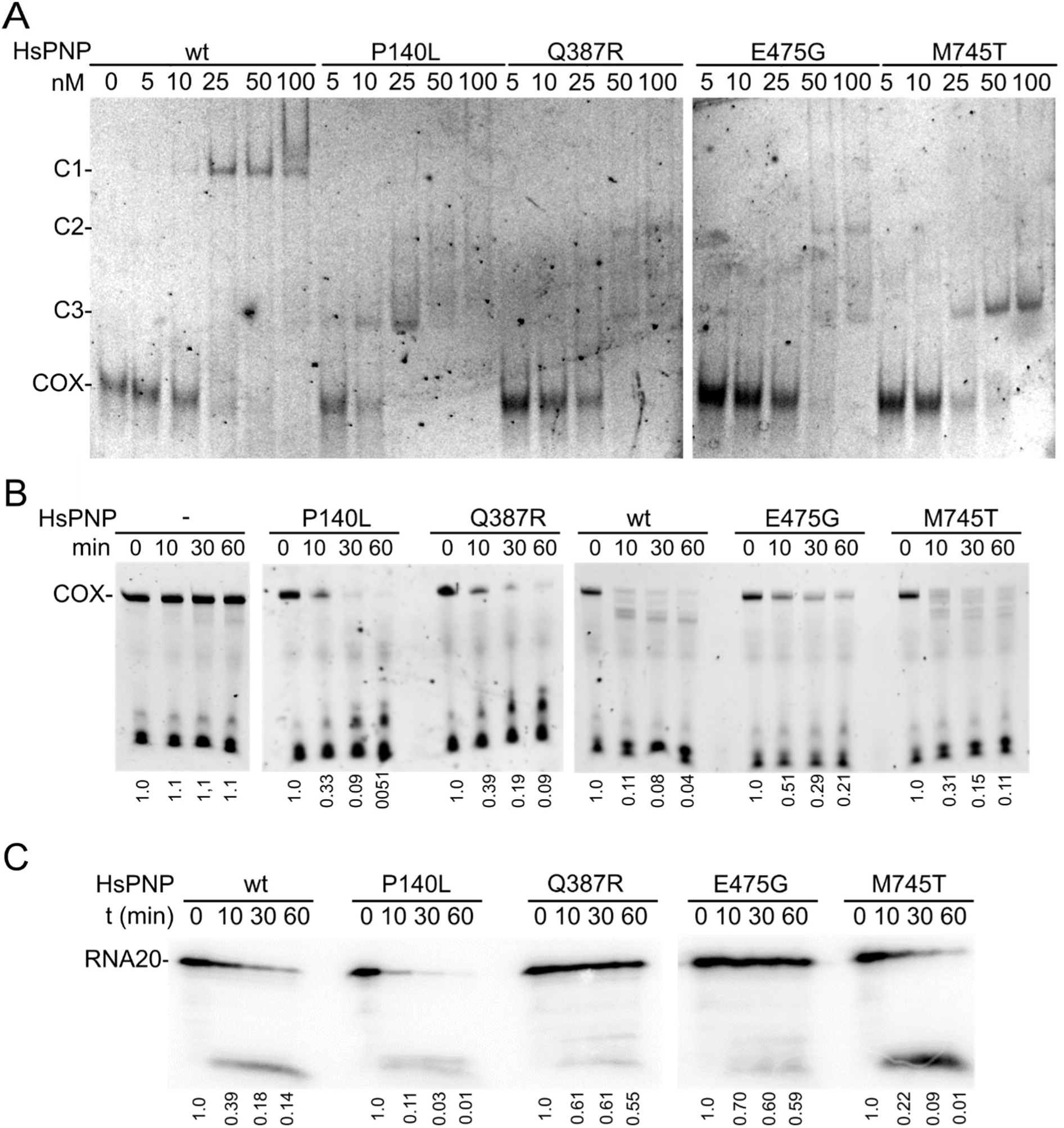
RNA binding and degradation activity. A. EMSA with fluorescent COX RNA (0.4 nM) incubated with the indicated amount of hPNPase (HsPNP) and run in 5% polyacrylamide native gels. C1, C2 and C3, complex 1, 2 and 3, respectively; COX, unbound COX probe. B and C. Degradation assay. 0.3 nM COX RNA (B) and 1.9 nM RNA20 (C) incubated at 37 °C with 50 nM hPNPase (hPNP) for the time specified on top of the lanes. Samples were run in 10% polyacrylamide denaturing gel. COX, full-length COX probe. In all panels, experiments representative of ≥ 2 replicates are shown. hPNPase variants are indicated with the mutated residue.

In contrast with these data, it was reported that the binding affinity of the Q387R and E475G hPNPases containing the additional S484A mutation, which inhibits catalytic activity, to a 15nt long poly(U) RNA was more than 200 times lower compared to the affinity of the single S484A mutant (6). This discrepancy could be due to the different probes used, as the COX RNA is longer than poly(U) and has the potential to form a stem-loop structure (**Supplementary Fig. S1**). To test this hypothesis, we examined the binding of hPNPase to RNA20, a heteropolymeric 20-mer RNA lacking secondary structures (**Supplementary Fig. S1**), which is efficiently bound by EcPNPase (40). As shown in **Supplementary Fig. 2C**, the wt hPNPase formed a complex with RNA20 with slow electrophoretic mobility, similar to complex 1. However, although hPNPase was in molar excess with respect to RNA20 (i.e. the protein at 100 nM was at ca. 160-fold molar excess), complex 1 formation involved only a minor fraction of the RNA20 indicating inefficient binding. Complex 1 was absent in presence of Q387R and E475G variants and strongly reduced with P140L, whereas smeared signals comigrating with complex 1 or more slowly were visible with M745T hPNPase. No other retarded complexes were visible with any tested hPNPase. Curiously, signals due to RNA species migrating faster than RNA20 were visible upon incubation with all the variants, and in particular all the mutated ones, indicating partial degradation of RNA20. Degradation was stimulated by the addition of 0.5 mM Pi and inhibited both in the presence of 50 mM Pi, a condition impairing hPNPase activity (24), and, at least for the P140L, in the absence of the detergent NP40 (**Supplementary Fig. S2D**), which could stabilize the mutated proteins. This suggests that RNA20 degradation in EMSA experiments depended on hPNPase and low Pi present as sample contaminant.

### *PNPT1* pathological mutations have different effects on the RNA degradation activity of hPNPase

RNA degradation activity of wt and mutated hPNPase variants was measured by incubating COX RNA or RNA20 with the enzyme at 37 °C in presence of 0.5 mM phosphate. Degradation of COX RNA by wild-type and M745T hPNPases occurred at similar rates and produced similar patterns of degradation intermediates. In contrast, full-length COX RNA disappeared more slowly in the presence of the other variants, particularly E475G, and the degradation pattern was different (**Fig. 3B**). As for the RNA20 degradation, only the P140L and M745T variants, which can form a complex with RNA20 in EMSA experiments, efficiently degraded the probe, whereas the Q387R and E475G variants were strongly impaired in RNA20 degradation (**Fig. 3C**). These two variants were able to degrade the RNA20 in the EMSA assay conditions **(Supplementary Figs. S2C-D)**.

We did not investigate further the reason of this discrepancy that could be due to the stabilization of the mutant proteins, and thus increased activity, at lower temperature (the samples were incubated at 21 and 37 °C in EMSA and degradation experiments, respectively) and in the presence of NP40, which is absent in the buffer used in degradation assays.

### hPNPase can trim ssDNA

Since EcPNPase can bind and degrade single-stranded DNA molecules (ssDNA) (41), we tested whether this was also true for hPNPase. We found that both the wt and the M745T hPNPases could (inefficiently) bind a ssDNA 20mer (i.e. DNA20) and trim it in presence of manganese and low phosphate, whereas the other variants did not bind, or shorten, the DNA20 (**Figs. 4A-B; Supplementary Fig. S2E**).

**Fig. 4.**
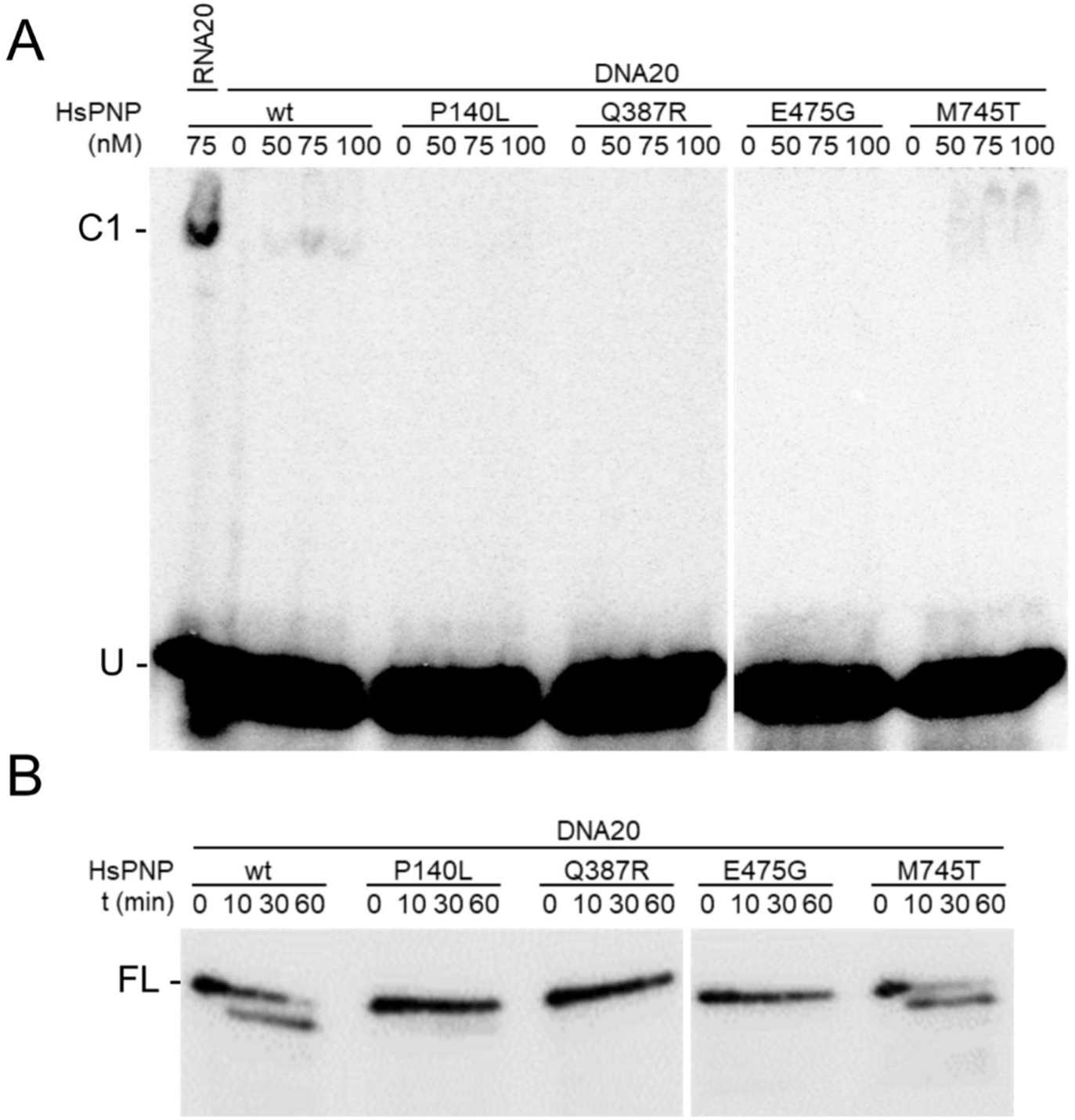
DNA binding and degradation activity. DNA20 is a DNA oligonucleotide with the same sequence as RNA20, and thymine in place of uracil. A. EMSA with radiolabelled DNA20 or RNA20 (0.6 nM) incubated with the indicated amount of hPNPase (HsPNP) and run in 5% polyacrylamide native gels. C1, complex 1; U, unbound DNA20 or RNA20 probes. B. Degradation assay. 1.9 nM DNA20 incubated at 37 °C with 50 nM hPNPase (HsPNP) for the time specified on top of the lanes. Samples were run in 10% polyacrylamide denaturing gel. FL, full-length DNA20. Experiments representative of two replicates are shown in both panels. Mutated hPNPase variants are indicated with the mutated residues.

### Mutations in *PNPT1* affect hPNPase-dependent phenotypes in *E. coli*

We studied the effect of *PNPT1* mutations in derivatives of the *E. coli* C-6001 strain in which the endogenous *pnp* locus was replaced by the *PNPT1* open reading frame optimized for bacterial codon usage (*PNPT1_Ec_*) (23). We first measured whether the mutations affected the hPNPase levels. We found that the amount of P140L hPNPase was significantly reduced compared to wt protein levels, whereas the levels of Q387R, E475G, and M745T mutant enzymes varied among different clones, with M745T being expressed, on average, at lower levels than the wild-type protein (**Fig. 5A**). To assess whether differences in protein levels might be due to differential stability of mutant hPNPases, we measured the protein amount at different time points after translation inhibition with streptomycin. As shown in **Fig. 5B**, P140L and M745T hPNPases were less stable than the wt protein, whereas the other two variants had comparable stability.

**Fig. 5.**
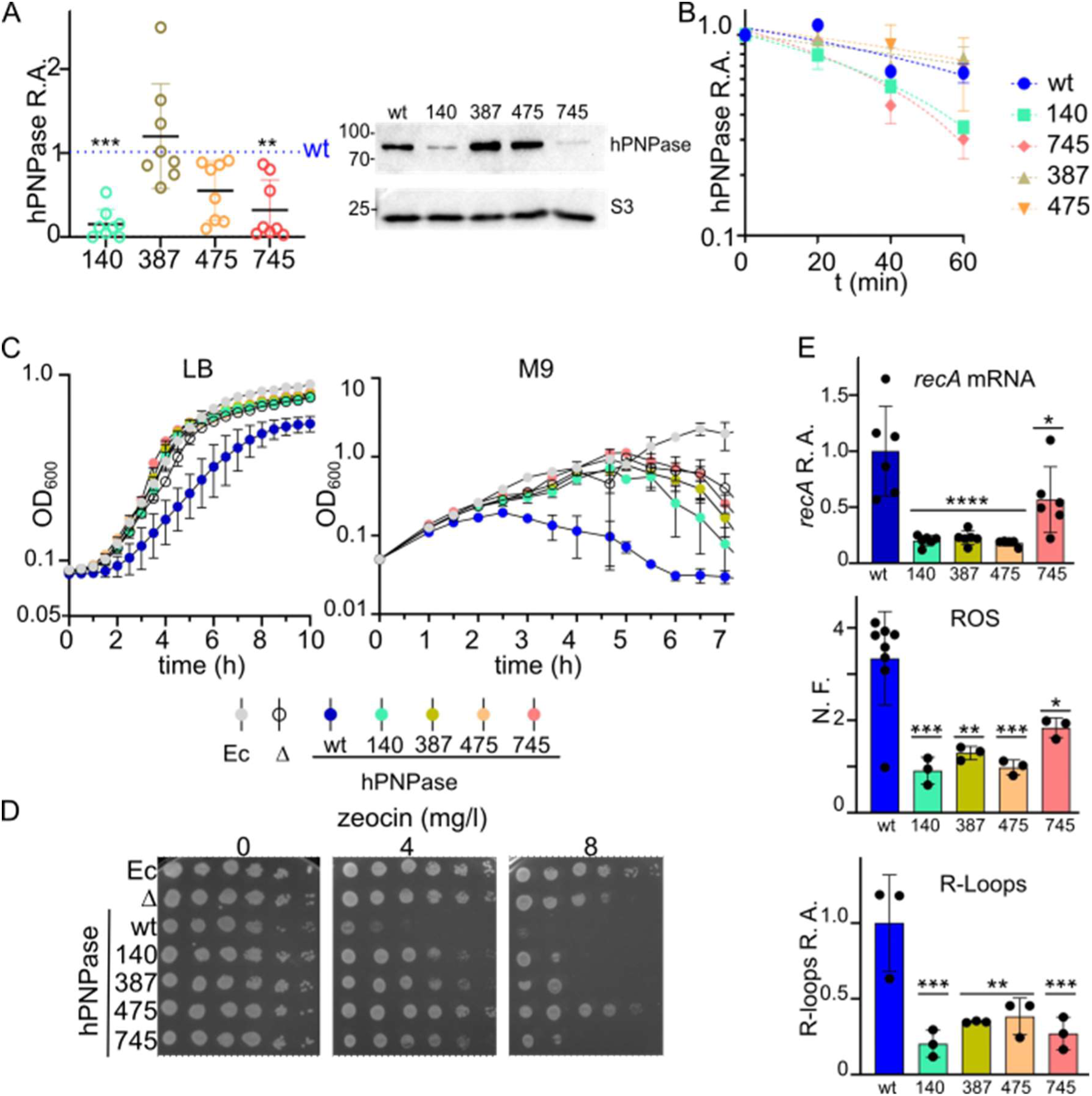
Phenotype of *E. coli* expressing mutated hPNPases. A. and B. Western blot analysis of protein levels and stability. Crude extracts were subjected to 10 % SDS-PAGE and immunoblot using hPNPase- and S3-specific (loading control) antibodies. hPNPase signals were quantified and normalized against S3 and wt hPNPase signals. A. R.A., relative amount. Symbols represent replicates, bars median values and stars significance of the difference with respect to wt hPNPase amount (dotted line) estimated with ANOVA and Dunnett’s multiple comparison. ***, P< 0.001; **, P < 0.01, no symbol, not significant. A representative experiment is shown on the right of the graph. The position of MW markers (kDa) is reported on the left. B. hPNPase stability. hPNPase signals were quantified with ImageLab (Bio-Rad) and normalized against the signal at t = 0.Symbols represent average (N = 2) with range. Bars are in some cases hidden by symbols. The exponential one phase decay curves interpolating data are shown as dotted lines (calculated with GraphPad Prism). C. Cultures were grown in LB or M9 supp measuring the optical density (OD_600_) at intervals. Symbols indicate average (N = 3) with standard deviation. D. Cultures of the indicated strains were serially diluted (×10 from left to right) and replicated on LB-agar containing the indicated amount of zeocin. Plates were incubated 24h at 37 °C. E. ROS, R-loops and *recA* mRNA levels. Histograms represent average with SD and dots represent single determinations. Significance of the difference with respect to the wt is indicated by stars and was evaluated with Welch (ROS) or ordinary ANOVA and Dunnett’s multiple comparison using GraphPad Prism. *, P < 0.05; **, P < 0.01; ***, P < 0.001; ****, P < 0.0001. N. F., Normalized fluorescence with respect to that of the Δ*pnp*; R.A., relative amount with respect to wt levels. A-E. Different PNPase loci are indicated as follows: Ec, endogenous *pnp* gene; wt, wt *PNPT1*_Ec_; Δ, Δ*pnp* mutant. *PNPT1_Ec_* mutant alleles are denoted by the number of the mutated codon.

We then tested whether *PNPT1* mutations impacted *E. coli* phenotypes observed in the presence of wt hPNPase, namely: slower growth in LB broth, increased aggregation in M9 minimal medium, enhanced sensitivity to zeocin and high ROS, R-loop and *recA* mRNA levels (23) (**Figs. 5C-E**).

All *PNPT1_Ec_* mutants exhibited a higher growth rate in LB and showed increased aggregation at higher optical density in M9 minimal medium compared to wt *PNPT1_Ec_*. Zeocin sensitivity was reduced in all mutants, particularly in the E475G mutant. Additionally, all mutants had lower levels of ROS, R-loops, and *recA* mRNA compared to wt *PNPT1_Ec_*. In particular, ROS and *recA* mRNA levels were strongly reduced in the P140L, Q387R, and E475G mutants, with a more moderate reduction observed in the M745T mutant. (**Figs. 5C-E**). Overall, the phenotype of the four mutants more closely resembled that of the C-5691 Δ*pnp* strain than that of the C-6001 strain expressing wt *PNPT1_Ec_,* indicating a loss of function of the mutated proteins.

### The homozygous P140L mutation is not tolerated in stable human cell lines

The effect of mutations in *PNPT1* has been studied primarily in patient-derived cells, which have a different genetic background. We developed a model in human cell lines to study the effect of pathological *PNPT1* mutations in a homogenous genetic background. Specifically, we selected two mutations associated with diseases of different severity, namely P140L and E475G, and we undertook genome editing using CRISPR-Cas system to introduce bi-allelic mutations causing these amino acid changes in hPNPase. To insert the c.419C>T p.(P140L) and c.1424A>G p.(E475G) mutations, we utilized two base editor-dCas9 systems: Be4-max, which converts CG base pairs to TA, and ABE7.10 converting AT to GC (42, 43). Since *PNPT1* mutations cause neurological defects, our initial choice was to use SH-SY5Y cells of neuronal origin for generating the human cell model. However, we switched to 293T cells because transfection of SH-SY5Y cells with the editing plasmids, using both lipofectamine and nucleofection, proved to be very inefficient. 293T cells were transfected with a mixture of three plasmids: one expressing either BE4-GFP or ABE-GFP base editors, a second expressing the targeting sgRNA, and a third expressing the mCherry transfection marker (42, 43). After 72 hours, GFP and mCherry double positive cells were single-cell sorted, and the amplified clones we obtained were analysed by PCR amplification and sequencing of the exon targeted by the base editor. The results and the genotype of the edited clones are summarized in **Table 1** and **Fig. 1B**. For the P140L mutation, around 47% of the sequenced clones carried a mutation in codon 140. However, all edited clones but one harboured the c.418C>T transition in the editing window, either in homozygosity or heterozygosity, resulting in the production of P140S hPNPase, which is classified as variant of uncertain significance in ClinVar database. Only one clone carried both the c.418C>T mutation in homozygosity and a heterozygous c.419C>T mutation, resulting in the co-expression of P140L and P140S hPNPase variants. This clone was used as a host for an additional round of editing with Be4-max and an sgRNA specific for introducing the c.419C>T mutation in the allele encoding the P140S variant. However, the vast majority of sorted cells did not proliferate, and despite five attempts with two different sgRNAs, we could not obtain a homozygous cell line producing only the P140L variant.

**Table 1.** Results of editing experiments.

|  | P140L |  |  |  |  |  | E475G |  |
| --- | --- | --- | --- | --- | --- | --- | --- | --- |
|  | Editing round |  |  |  |  |  | 1 | 2 |
|  | 1 | 2 |  |  |  |  |  |  |
| Sorted cells | 192 | 192 | 192 | 192 | 384 | 384 | 384 | 192 |
| Clones obtained | 113 | 6 | 24 | 7 | 14 | 12 | 170 | 64 |
| Sequenced<sup>a</sup> | 36 | 6 | 24 | 7 | 14 | 12 | 51 | 16 |
| Codon Edited<sup>b</sup> | 17 | 0 | 0 | 0 | 0 | 0 | 15 | 8 |
| Desired homozygous clones | 0 | 0 | 0 | 0 | 0 | 0 | 0 | 2 |

<sup>a</sup>Relevant *PNPT1* exons were sequenced as detailed in the text
<sup>b</sup>All the identified mutations in the 140 and 475 codons are considered

Regarding the E475G mutation, after the first editing round, codon 475 was edited in 15 out of 51 sequenced clones, with an editing efficiency of approximately 29%. Seven clones carried the c.1425A>G transition, resulting in a silent aminoacidic substitution, whereas the remaining five clones harboured the desired c.1424A>G mutation in heterozygosity. One of these mutants was used for a second round of ABE7.10 editing, which resulted in the identification of eight edited clones, of which two carried the desired homozygous c.1424A>G mutation, whereas the other six carried the c.1425A>G silent mutation in heterozygosity with c.1424A>G (**Table 1** and **Fig. 1B**).

The level of *PNPT1* mRNA assessed by RT-qPCR was not altered in the mutant cell lines, whereas the level of hPNPase protein was reduced to 60% of the normal level in the P140L/P140S mutant cell line. On average, the protein level remained unchanged in the homozygous P140S and heterozygous E475G lines, while it was reduced by approximately 20% in the homozygous E475G line. Moreover, the level of E475G hPNPase showed fluctuations in replicate experiments (**Figs. 6A-B**).

**Fig. 6.**
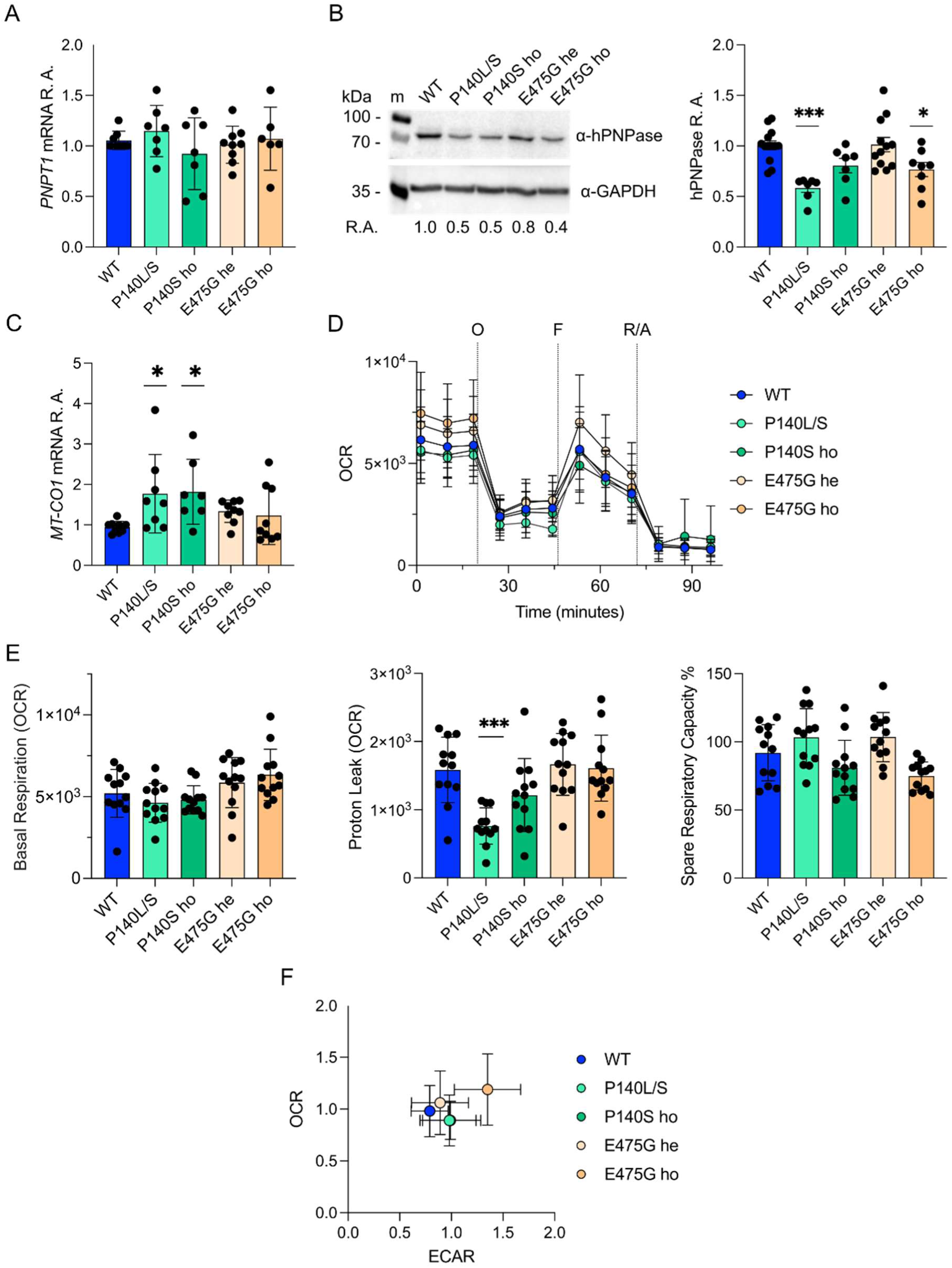
Phenotype of stable cell lines with *PNPT1* mutations. A and B. *PNPT1* expression at the mRNA (A) and protein (B) levels estimated by RT-qPCR and western blotting, respectively. In the two graphs, symbols represent single determinations and bars the average with standard deviation. B. Typical results of a western blotting experiment are shown on the left. GAPDH was used as loading control. C. *MT-CO1* expression levels measured by real-time RT-PCR. Dots, single determinations; bars, average with standard deviation. D-F. Agilent Seahorse XF analysis of mitochondrial stress. The graphs summarize the results of three independent experiments, with four technical replicates each. Average and SD are plotted in all cases. D. Oxygen consumption rate (OCR, measured in pmol/min/1000 cells) over time following the addition of mitochondrial inhibitors: O, oligomycin; F, FCCP; R/A, rotenone and antimycin A. E. Basal respiration (pmol/min/1000 cells), proton leak (pmol/min/1000 cells), and relative spare respiratory capacity (%). F. OCR plotted over the extracellular acidification rate (ECAR). Data from each experiment were normalized to the respective average values of OCR and ECAR. A-F. 293T are indicated as wt and 293T mutant derivatives are denoted by the amino acid substitutions in hPNPase. he, heterozygous; ho, homozygous. A to E. Significance of the difference with the wt determined with ANOVA and Dunnett’s multiple comparison (calculated with GraphPad Prism) is indicated by stars. ***, P < 0.001; *, P < 0.05. Non-significant comparisons are not indicated.

### Mitochondrial function is only slightly affected in *PNPT1* mutant cell lines

To investigate the effect of *PNPT1* mutations on the growth and mitochondrial function, we measured the growth rate, mitochondrial DNA (mtDNA) content and mitochondrial-related oxygen consumption rate. Both the growth rate and the copy number of mtDNA, measured as the ratio between mitochondrial and genomic DNA content, remained unvaried across all tested lines, suggesting no major consequences of the mutations on mitochondrial number (**Supplementary Figs. 3A-B**).

To analyse the effect of the mutations on the mitochondrial function, we measured the expression of the mitochondrial cytochrome C oxidase I gene *MT-CO1* by RT-qPCR and the respiratory activity with the Seahorse XF Cell Mito Stress Test. We found that *MT-CO1* levels were slightly higher in the P140L/S compound heterozygous and P140S homozygous cells compared to the *PNPT1*^+^, and not altered in the other lines. Concerning the respiratory activity, no significant difference was found among the cell lines in terms of spare respiratory capacity or basal respiration, whereas proton leak was slightly reduced only in the P140L/S compound heterozygous (**Figs. 6D-E**). Plotting the Oxygen Consumption Rate (OCR) against the Extracellular Acidification Rate (ECAR), which reflects the accumulation of lactate due to pyruvate reduction from glycolysis, revealed that only the E475G homozygous line was slightly separated, while the other lines clustered closely with the wt (**Fig. 6F**).

### The E475G mutation strongly impacts gene expression

To better understand the effect of the hPNPase E475G mutation, the transcription profile of the mutant line was compared with that of the wt and the heterozygous lines. We found that the mutation did not change the relative amount of mitochondrial *vs*. total RNA but affected the overall transcription profile (**Supplementary Tables S2-S3; Figs. 7A-B** and **Supplementary Figs. S3C-D**). 342 and 172 genes were differentially expressed in the wt *vs*. E475G homozygous or heterozygous lines, respectively, and 142 in the heterozygous *vs*. homozygous E475G lines, considering in all cases only comparisons with |log2(FoldChange)| ≥ 1 and FDR < 0.01.

**Fig. 7.**
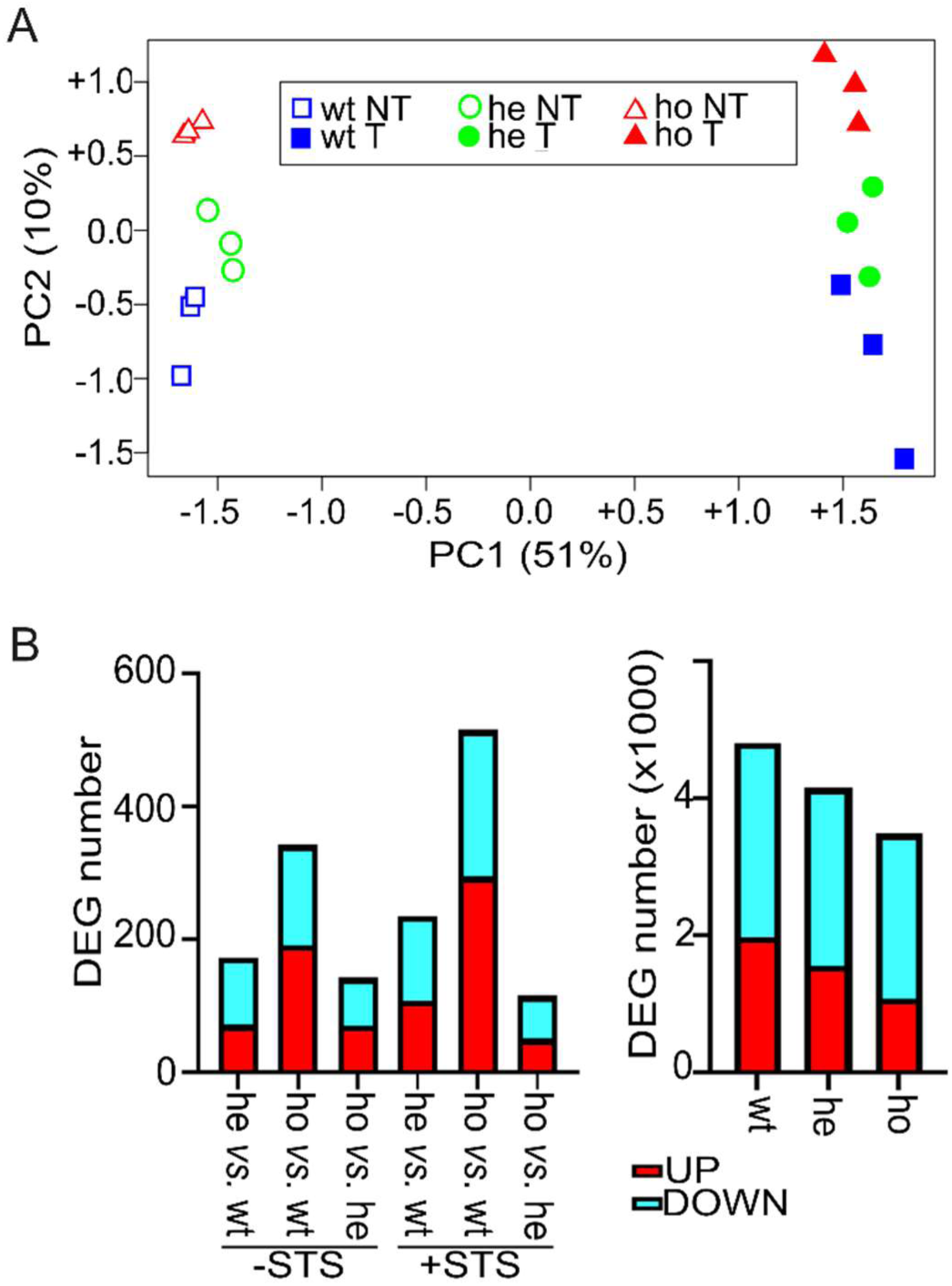
Differentially expressed genes (DEGs) in human cells expressing E475G hPNPase. A. Principal Component analysis of transcriptomic data. wt, 293T *PNPT1*^+^; he and ho, heterozygous and homozygous E475G mutation, respectively; NT, not treated; T, treated with staurosporine. B, left panel. Number of up- and down-regulated genes with |log2(FoldChange)| ≥ 1 and FDR < 0.01 in the indicated pairwise comparisons. STS, staurosporine. Right panel, numbers of up- and down-regulated genes in wt and mutant cell lines, indicated as in A, upon STS incubation.

Interestingly, although 293T cells are not of neuronal origin, genes encoding factors involved in neuronal development and nervous disorders like SPARC, SYNDIG1, SLC19A3 and FOSB (44–54) are among the genes with the highest up- or down-regulation level in the comparison between the E475G homozygous and the *PNPT1*^+^ lines (**Supplementary Table S2**). Accordingly, Gene Ontology (GO) analysis for overrepresented gene sets in the homozygous *vs.* wt comparison identified processes mostly related to the development of nervous system (**Table 2**).

**Table 2.** Top10 significant p-values and q-values for GO Biological Process 2023.

| Gene Ontology 2023 Term<sup>a</sup> | p-value | q-value |
| --- | --- | --- |
| **p. E475G homozygous vs. wt cells** |  |  |
| Positive Regulation of Transcription by RNA Polymerase II (GO:0045944) | 3.26E-07 | 0.001 |
| Nervous System Development (GO:0007399) | 1.41E-06 | 0.003 |
| Positive Regulation of Neurogenesis (GO:0050769) | 3.69E-06 | 0.005 |
| Neuron Projection Development (GO:0031175) | 6.11E-06 | 0.005 |
| Regulation of Cell Cycle (GO:0051726) | 7.70E-06 | 0.005 |
| Dendrite Morphogenesis (GO:0048813) | 8.01E-06 | 0.005 |
| Regulation of Dendritic Spine Morphogenesis (GO:0061001) | 9.13E-06 | 0.005 |
| Positive Regulation of Signal Transduction (GO:0009967) | 1.07E-05 | 0.005 |
| Peptidyl-Serine Phosphorylation (GO:0018105) | 1.42E-05 | 0.006 |
| Neuron Migration (GO:0001764) | 1.47E-05 | 0.006 |

<sup>a</sup>Analysis performed with Enrichr (37) on DEGs with FDR > 0.01 in the indicated pairwise comparison. p- and q-values were calculated by the software using the Benjamini-Hochberg method for correction for multiple hypotheses testing.

Since hPNPase has been implied in cytoplasmic mRNA decay upon apoptosis induction (55), we analysed the effect of the E475G hPNPase mutation in cells incubated six hours with staurosporine (STS). 514 and 234 genes were differentially expressed in the wt *vs*. E475G homozygous or heterozygous lines, respectively, and 115 in the heterozygous *vs*. homozygous E475G lines, also in this case with |log2(FoldChange)| ≥ 1 and FDR < 0.01 as thresholds (**Supplementary Table S2** and **Figs. 7A-B**). In all pairwise comparisons between samples treated with staurosporine and untreated ones, genes involved in apoptosis regulation were significantly enriched, as expected (**Supplementary Table S4**). The number of differentially expressed genes (DEGs) in such comparisons decreased in the following order: wt > heterozygote > homozygote (**Fig. 7B**). These findings are consistent with the hPNPase E475G mutation affecting mRNA levels during apoptosis.

## DISCUSSION

Establishing a correlation between specific *PNPT1* mutations and their phenotypic consequences is challenging due to the rarity of patients diagnosed with *PNPT1*-dependent diseases, the presence of compound heterozygosity in most cases, and the still incomplete understanding of hPNPase function in the various cellular processes it has been involved in, sometimes in an anecdotal manner (4). The “humanized” *E. coli* model, expressing human PNPase (23), along with the new model in 293T cells, may provide valuable insights into the effects of various pathological mutations on hPNPase function. Although such models are too simple to recapitulate the phenotype of patients, we anticipate that in perspective, they will be valuable tools to understand the molecular mechanisms at the root of the disease insurgence, to predict the phenotypic effects of *PNPT1* mutations and to develop strategies to correct, or mitigate, protein defects caused by the mutations.

In the *E. coli* model, the four *PNPT1_Ec_* missense mutations we tested resulted in the loss or attenuation of all phenotypic traits observed in the presence of the wt enzyme. Consistent with this observation, all mutations alter the structure of the protein and/or its RNA binding activity, albeit to varying extents. However, the effects of the mutations are not identical, as strains producing M745T and E475G hPNPases showed a milder reduction in ROS and *recA* levels in the former and a greater reduction in zeocin sensitivity in the latter, compared to the other mutant strains. This finding suggests that hPNPase may be involved in diverse cellular pathways, with mutations affecting them to different degrees.

The results of *in vitro* analyses we conducted on purified proteins help rationalizing the results we obtained in *E. coli* for the M745T variant. Indeed, the M745T hPNPase migrated as a homotrimer in BN gel, and had RNA and DNA binding efficiency and degradation similar to the wt. However, complexes formed with the RNA and DNA by this mutated protein have altered migration, suggesting that the M745T mutation somehow impacts the protein structure and/or the architecture of complexes with the RNA. Consistently, the fluctuations we observed in M745T protein levels and its reduced stability in *E. coli* may be due to protein misfolding, which promotes aggregation. This, in turn, could lead to protein degradation or targeting to inclusion bodies. The structure of the S1 domain, where M745 is located, has not been solved, as such domain is absent in the available structure of hPNPase (13). However, based on the AlphaFold2 prediction, M745 is positioned in the hydrophobic core of the domain. While the mutation to threonine does not appear to cause significant steric clashes, it may force a reorganization of neighboring residues, including Y735, with potential long-range effect (**Supplementary Fig. S4A**).

The other three mutations we tested affected the quaternary structure of purified hPNPase, as shown by altered migration in BN gels of the three mutated variants. Specifically, consistent with published data (6, 26), we confirmed that purified Q387R and E475G tend to dissociate and form dimers in BN gels. This behavior is compatible with the involvement of Q387 and E475 in the formation of hydrogen bonds - destroyed by the mutations - with residues of the flanking protomer stabilizing the quaternary structure (6, 26). These two variants do not bind the RNA20 (or DNA20) and are those showing the strongest reduction in both RNA20 and COX degradation activity. The two mutations did not have identical effects on *PNPT1_Ec_* expression in *E. coli*: the Q387R hPNPase level remained consistently high, whereas the E475G hPNPase level fluctuated across different cultures. However, both mutations impaired all hPNPase-dependent phenotypes.

In patients with homozygous *PNPT1* mutations, the condition caused by the Q387R hPNPase is more severe than that caused by the E475G mutant protein (21). Since only a few case studies have been reported, we cannot rule out that the patients’ genetic background influences the severity of the disease. However, as in principle the Q387R mutation can be generated using the ABE7.10 base editor, we plan to study its effect in the 293T model we presented in this paper.

In agreement with the relatively mild effect of the E475G hPNPase in patients (21), we were able to generate cell lines heterozygous and homozygous for the mutation causing such substitution. The mutant lines did not show defects in cell proliferation, mtDNA, or mtRNA content, and the homozygous mutant exhibited only a slight shift in the ECAR *vs*. OCR parameter toward glycolytic metabolism, suggesting subtle changes in mitochondrial activity. However, the overall transcription profile was clearly affected by the mutation at the heterozygous and, to a stronger degree, homozygous state. The effect of the E475G mutation on gene expression was detectable even in apoptotic cells, as the number of differentially expressed genes between cells treated with or without staurosporine was lower in the mutant than in the *PNPT1*^+^ or the heterozygote. This finding is consistent with a PNPase role in controlling the stability of polyadenylated RNA during apoptosis (55), an activity that could be impaired, or limited, by the mutation.

Interestingly, genes involved in neuronal development were among those most sensitive to the mutation in spite of the non-neuronal origin of 293T cells. Accordingly, Gene Ontology analysis of differentially expressed genes upon PNPase gene knock-out in mouse embryonic fibroblasts identified pathways related to neural functions including axonogenesis, axon guidance and axon development among those overrepresented (18). It remains to be established whether the effect of hPNPase defects on the expression of such genes depends on a direct action in controlling the level of key mRNAs by hPNPase or it is an indirect effect on gene expression due to subtle differences in mitochondrial function, which is known to regulate neurogenesis (56). Introducing other homozygous *PNPT1* mutations in the cell model may help discriminating between these two hypotheses. Interestingly, it has been recently suggested that hPNPase may play a direct role in neuronal differentiation by degrading nuclear encoded mRNAs when cells are exposed to differentiation cues (57).

We were unable to obtain a cell line expressing only the P140L hPNPase variant, which is caused by a mutation that, to our knowledge, has never been reported in homozygous patients. P140 is highly conserved and is located in the active site, although it does not directly participate in the catalysis or RNA binding (4, 27). This residue is positioned at the C-terminus of an alpha-helical region where its side chain occupies a tiny space with only backbone atoms in its vicinity. The space available does not allow to fit a substitution into leucine without a large perturbation of its surrounding (**Supplementary Figs. S4B-C**). The purified P140L variant barely entered BN gels, with the residual protein running as a trimer. Since the protein runs normally in SDS-PAGE, we think that it may form high molecular weight aggregates preventing gel entry in BN electrophoresis conditions. Accordingly, the levels of the P140L hPNPase were reduced in both *E. coli* and 293T cells, which is compatible with protein aggregation/degradation ensuing misfolding. In contrast with our data, no effect on trimer assembly was previously observed in BN PAGE of proteins extracted from the myoblasts of a compound heterozygous p.R136H/p.P140L patient, and the amount of trimeric hPNPase was not reduced in such cells (27). In the compound heterozygous state, the presence of the p.R136H allele might partially compensate for the defects caused by the P140L mutation, resulting in less impact on the overall protein levels. However, given the severity of the patient’s condition - diagnosed with Leigh syndrome and passing away at the age of 2 years (27) - the activity of the protein, whether as an R136H homotrimer or a heterotrimer formed by both variants, is likely to be severely impaired.

Interestingly, while we could not establish stable P140L homozygous lines in spite of multiple attempts, we readily obtained P140S homozygous and P140L/P140S compound heterozygous mutants. Higher editing efficiency of *PNPT1* position 418 in comparison to position 419 of hPNPase ORF (see **Fig. 1B**) may have contributed to this result. However, considering the severity of the disease caused by the P140L mutation, its impact on protein levels in systems as different as bacterial and human cells and its inability to enter native gels indicative of relevant structural defects, it seems likely that in homozygosity, the P140L mutation is lethal in 293T cells, whereas the P140S, although causing slight defects in mitochondrial function, is tolerated. Indeed, serine is small enough to fit in the space occupied by P140 without causing a large perturbation of its surroundings (**Supplementary Figs. S4B-D**).

Unexpectedly, all mutant hPNPases retain the ability to bind the COX probe forming complexes with higher in-gel mobility than the complex 1 formed by the wild-type protein and by the M745T at high concentration. We believe complexes 2 and 3 involve hPNPase forms with altered oligomerization state (i.e. dimers/monomers). The alternative explanation could be that mutant hPNPase preparations are contaminated with a protein able to bind COX, and not RNA20, probe. However, we could not detect any different band in SDS-PAGE of purified mutant *vs*. wt hPNPase. Moreover, in EMSA experiments performed with COX RNA and serial dilutions of the wt hPNPase, some diluted samples occasionally form complexes 2 and 3 (see **Supplementary Fig. S2A**). This is consistent with such complexes being not due to a contaminating protein, which should be serially diluted together with hPNPase, but to a shift in the equilibrium between different hPNPase oligomerization states occurring in some samples. Since the mutated variants do not bind short RNAs like the RNA20 or a 15-mer poly(U) RNA (6), and considering the low RNA sequence specificity of PNPases of different species (3), it seems likely that it is either the length of the RNA and/or the potential presence of secondary structures that are critical for binding by dimeric/monomeric forms of hPNPase, and not a specific sequence in COX RNA. This result may have implications for *in vivo* hPNPase activity. It is likely that the mitochondrial localization signal, which is removed only once the enzyme has reached its mitochondrial destination, impairs trimerization, similar to the destabilizing effect observed with the shorter Histag (6). However, our findings suggest that altering the oligomerization state of hPNPase does not completely abolish its RNA binding activity towards certain mRNAs.

We showed here that human PNPase like the *E. coli* and *B. subtilis* orthologues can bind, albeit with very low efficiency, the ssDNA and trim it, and that all mutations but the M745T impair these activities. In bacteria, DNA binding and degradation by PNPase has been related to its involvement in DNA repair and homologous recombination (40, 41, 58). However, the physiological meaning of DNA binding and processing by PNPase remains to be established not only in the human, but also in the bacterial systems.

## ACKNOWLEDGMENTS

We thank Moira Paroni and Eva Pinatel for helpful discussions, Claudia Bazzini and Cristina Ruberti for assistance with FACS cell sorting and Seahorse experiments, and Claudia Fracassetti and Ilaria Pasquino for technical assistance.

## DATA AVAILABILITY

RNA-Seq raw data are available at GEO repository (accession number GSE277568).

