## Supplementary Fig. S1 for "Pathological PNPase variants with altered RNA binding and degradation activity affect the phenotype of bacterial and human cell models"

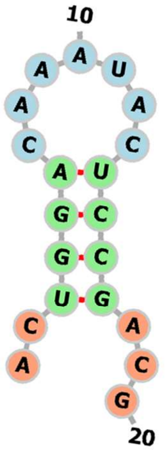


**RNA20**

**ΔG = -2.13 kcal/mol**

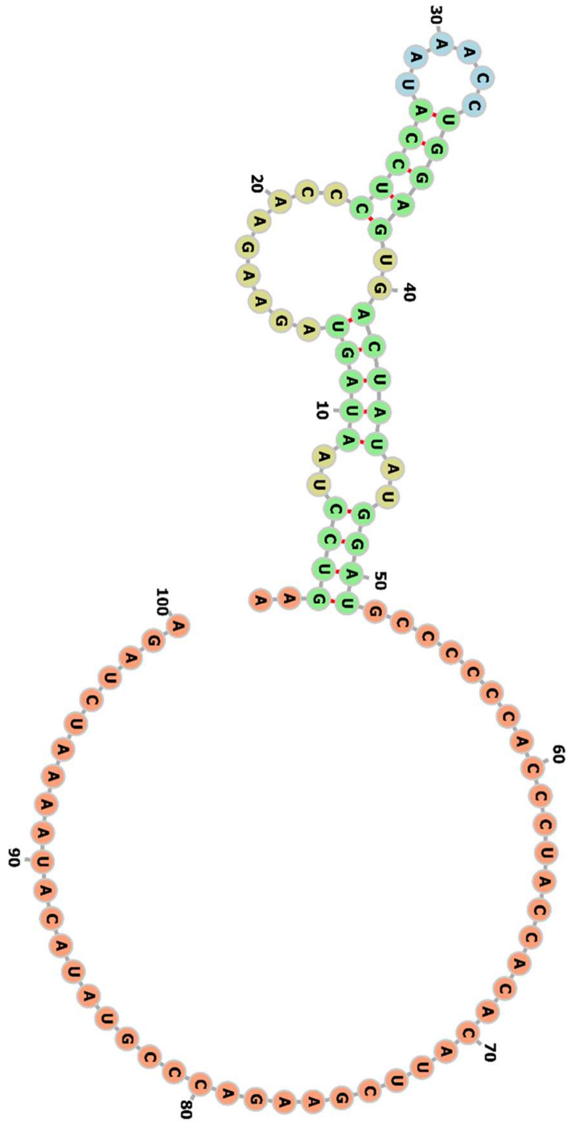


**COX RNA**

**ΔG = -12.2 kcal/mol**

**Supplementary Fig. S1. Predicted secondary structure of RNA20 and COX riboprobes.** The secondary structure and free energy of the thermodynamic ensemble were predicted with RNAFold[9]. According to OligoCalc[10], DNA20 had no potential for hairpin formation or self-annealing.
