## Supplementary Fig. S2 for "Pathological PNPase variants with altered RNA binding and degradation activity affect the phenotype of bacterial and human cell models"

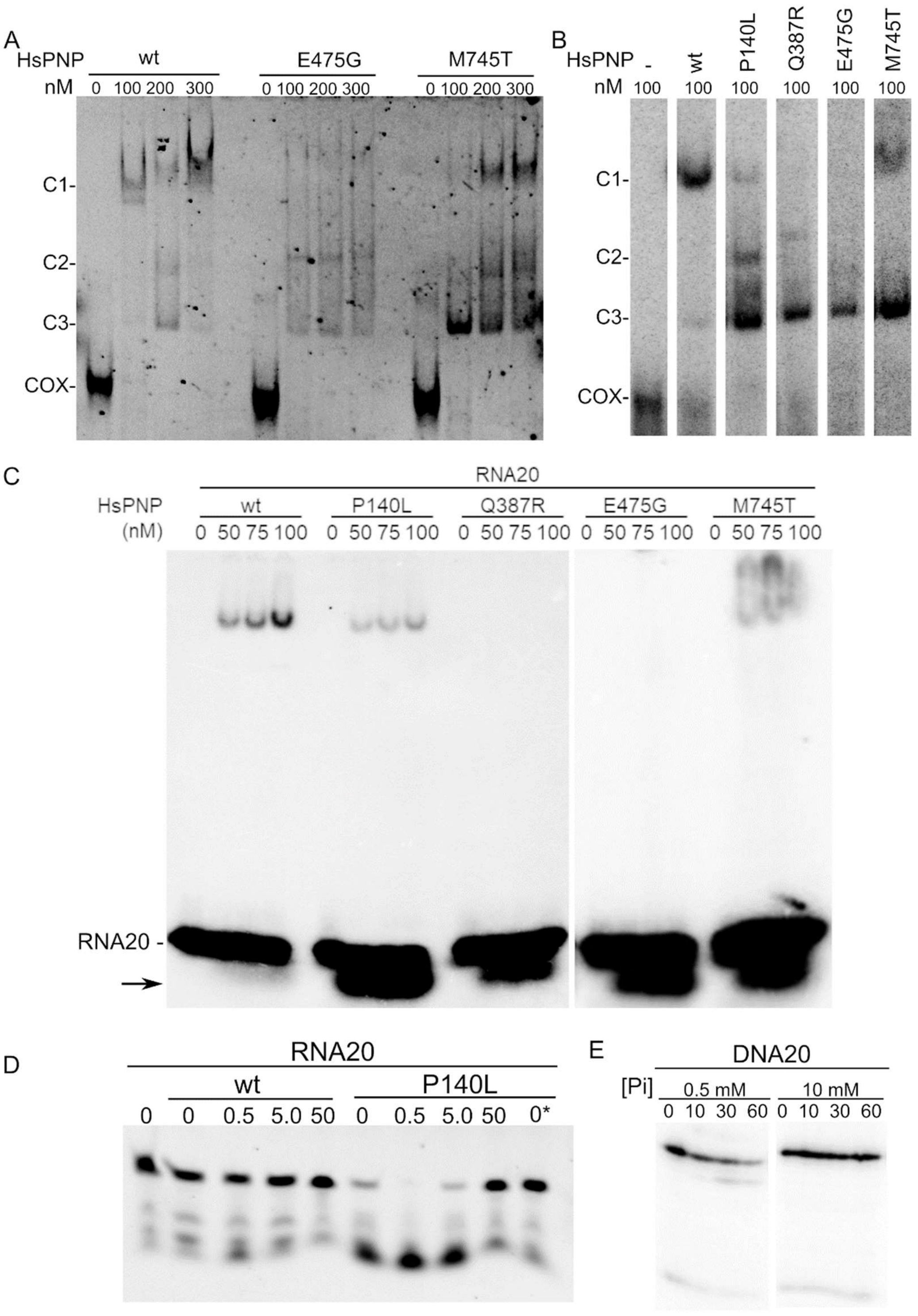


**Supplementary Fig. S2. hPNPase RNA binding and Pi sensitivity of RNA20 and DNA20 degradation activity.** In all panels, hPNPase variants are indicated with the mutated residue. A, B and C. EMSA with fluorescent (A) or radiolabelled (B) COX RNA (0.4 nM) or radiolabelled RNA20 (C; 0.6 nM) incubated with the indicated amount of hPNPase (HsPNP) and run in 5% polyacrylamide native gels. C1, C2 and C3, complex 1, 2 and 3, respectively; COX, unbound COX RNA; RNA20, unbound RNA20; arrow, RNA20 degradation fragments. D. Degradation assay. 10 nM RNA20 fluorescently labelled at the 5'-end was incubated in the EMSA buffer with 50 nM wt or P140L hPNPases for 30 min at 37 °C. Pi amount (mM) is indicated on top of the lanes. 0*, sample incubated in EMSA buffer without NP40. Samples were run in 10% polyacrylamide denaturing gel. E. DNA20 degradation upon incubation at 37 °C for the time indicated in min on top of the lanes with 50 nM wt hPNPase and 0.5 or 10 mM Pi.
