## Supplementary Fig. S3 for "Pathological PNPase variants with altered RNA binding and degradation activity affect the phenotype of bacterial and human cell models"

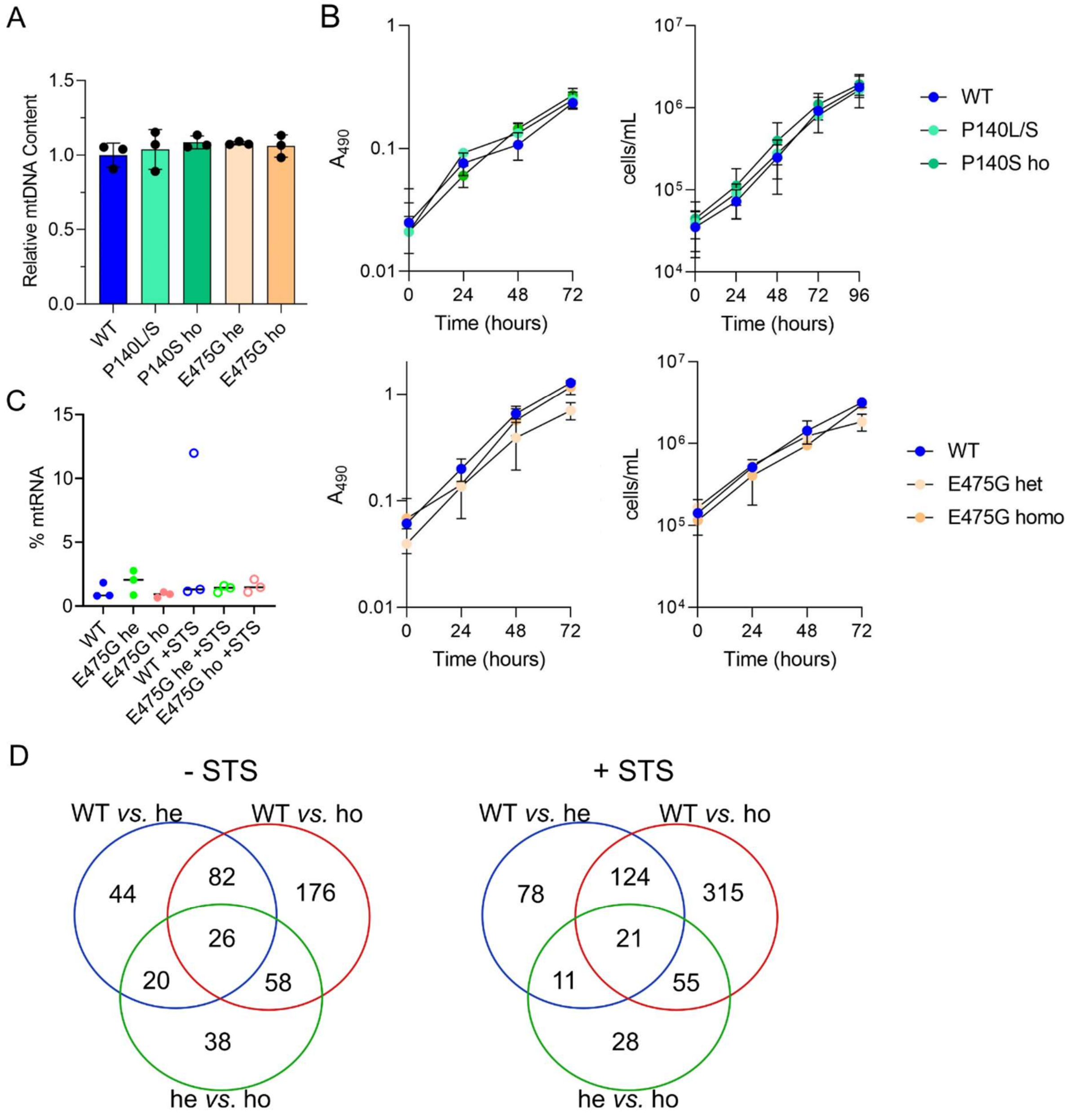


**Supplementary Fig. S3. Phenotypic and transcriptomic analyses of 293T lines with *PNPT1* mutations**. A. Relative mtDNA content determined by qPCR (N = 3). B. Growth rate measured by MTS assay as increasing absorbance at 490 nm (left panels) and by trypan blue exclusion tests (right panels) in 293T mutant cell lines. Symbols indicates the average and the standard deviation of three and two technical replicates for the MTS assay and the trypan blue exclusion test, respectively. Plots are representative of three independent experiments. C. Percentage of mtRNA reads on the total reads mapped on mitochondrial and nuclear genomes. D. Venn diagram of DEGs with FDR<0.01 and |log2(FoldChange)| ≥ 1. wt, 293T cell line; he and ho, 293T cells with the heterozygous and homozygous p.(E475G) mutation, respectively.
