## Supplementary Fig. S4 for "Pathological PNPase variants with altered RNA binding and degradation activity affect the phenotype of bacterial and human cell models"

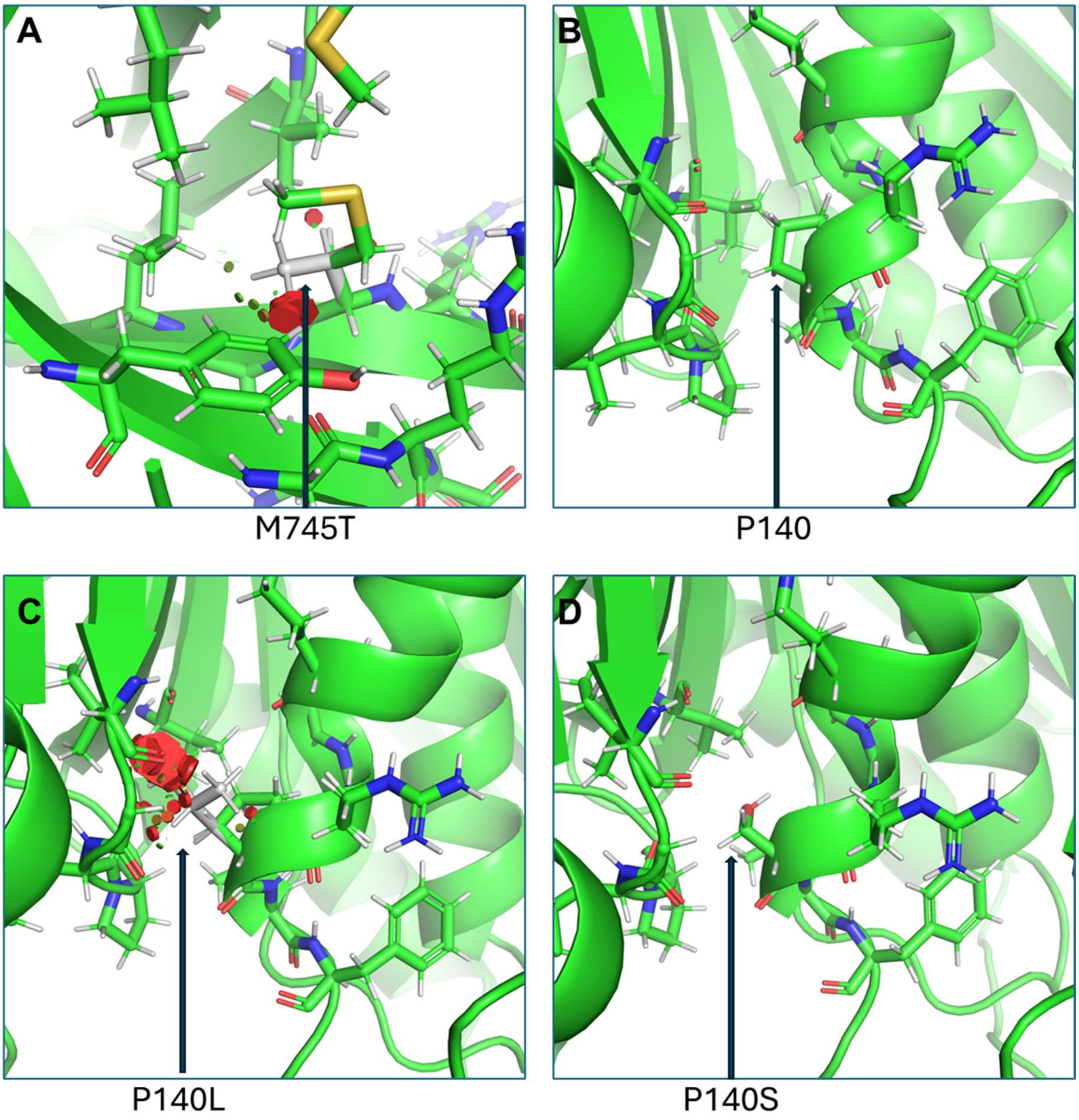


**Supplementary Fig. S4. Zoom in into the effect of M745T and P140L/S mutations**. A. M745 is located in the S1 domain as predicted by AlphaFold2[11]. The mutation to threonine can lead to clashes with Y735. B. P140 as observed in the crystal structure deposited (pdb code 3U1K). C. Clash analysis in case of P140L mutation as observed by a direct substitution with no structural modifications. D. Clash analysis in case of P140S mutation as observed by a direct substitution with no structural modifications.
