## Supplementary Table S1 for "Pathological PNPase variants with altered RNA binding and degradation activity affect the phenotype of bacterial and human cell models"

**Supplementary Table S1.** ***Escherichia coli*** **strains, plasmids, human cell lines and oligonucleotides**

*E. coli* strains

| Strain | Features | Reference |
|---|---|---|
| C-1a | *E. coli* strain C, prototrophic | (1) |
| C-5691 | C-1a *Δpnp-751* | (2) |
| C-6001 | C-1a *Δpnp::PNPT1*$_{Ec}$ | (3) |
| C-6027 | C-1a *Δpnp::PNPT1*$_{Ec}$*- P140L* | This work |
| C-6028 | C-1a *Δpnp:: PNPT1*$_{Ec}$*-Q387R* | This work |
| C-6029 | C-1a *Δpnp:: PNPT1*$_{Ec}$*-E475G* | This work |
| C-6030 | C-1a *Δpnp:: PNPT1*$_{Ec}$*-M745T* | This work |
| SHuffle® T7 | F′ *lac, pro, lacI*$^q$ */ Δ(ara-leu)7697 araD139 fhuA2 lacZ::T7 gene1 Δ(phoA)PvuII phoR ahpC* galE (or U) galK λatt::pNEB3-r1-cDsbC* (Spec$^R$, *lacI*$^q$) *ΔtrxB rpsL150*(Str$^R$) *Δgor Δ(malF)3* | New England Biolabs |

*E. coli* Plasmids

| Name | Features | Reference |
|---|---|---|
| pCP20 | Flippase encoding plasmid | (4) |
| pKD13 | KanR encoding plasmid | (4) |
| pKD46 | λRED plasmid | (4) |
| pET28b-hPNP | pET28b (+) derivative encoding hPNP without the mitochondrial localization signal (MLS) (aa 46-783), with an N-terminal 6x His-tag; KanR; T7 promoter. | (5) |
| pET28b-hPNP$^{P140L}$ | pET28b (+) derivative encoding hPNP P140L without the mitochondrial localization signal (MLS) (aa 46-783), with an N-terminal 6x His-tag; KanR; T7 promoter. | This work |
| pET28b-hPNP$^{Q387R}$ | pET28b (+) derivative encoding hPNP Q387R without the mitochondrial localization signal (MLS) (aa 46-783), with an N-terminal 6x His-tag; KanR; T7 promoter. | This work |

| | | |
|---|---|---|
| pET28b-hPNP$^{E475G}$ | pET28b (+) derivative encoding hPNP E475G without the mitochondrial localization signal (MLS) (aa 46-783), with an N-terminal 6x His-tag; KanR; T7 promoter. | This work |
| pET28b-hPNP$^{M745T}$ | pET28b (+) derivative encoding hPNP M745T without the mitochondrial localization signal (MLS) (aa 46-783), with an N-terminal 6x His-tag; KanR; T7 promoter. | This work |
| pET29b (+) | T7 promoter, kanR | Novagen |
| pET29b-hPNP | pET29b (+) derivative encoding hPNP without the MLS (aa 46-783), with a C-terminal 6x His tag, KanR; T7 promoter. | This work |

Human cell Plasmids

| **Name** | **Features** | **Reference** |
|---|---|---|
| pCMV_ABEmax_P2A_GFP | PCMV encoding ABEmax base editor. AmpR. | (6) |
| pCMV3_BE4max_P2A_GFP | PCMV encoding BE4max base editor. AmpR. | (6) |
| pFYF1320_E475G | PFYF1320 EGFP Site#1 derivative designed with the sgRNA targeting the c.1424A locus in exon 17 of the *PNPT1* human gene. | This work |
| pFYF1320_P140L | PFYF1320 EGFP Site#1 derivative designed with the sgRNA targeting the c.419C locus in exon 5 of the *PNPT1* human gene. | This work |
| pFYF1320_P140L/S | PFYF1320-P140L derivative designed with the sgRNA targeting the c.418T; c.419C locus in exon 5 of the *PNPT1* human gene. | This work |
| pFYF1320_P140L/S-L141L | PFYF1320-P140L derivative designed with the sgRNA targeting the c.418T; c.419C locus in exon 5 of the *PNPT1* human gene. | This work |
| PmCherry | PmCherry-N1-EXO1b derivative without EXO1b. KanR. | (7) |

Human cell lines

| **Name** | **Features** | **Reference** |
|---|---|---|
| 293T | | (8) |
| EK81 | 293T clone with NM_033109.5 (*PNPT1*): c.1424A>G: p.(Glu475Gly) (heterozygous). | This work |

| | | |
|---|---|---|
| EK89 | 293T clone with NM_033109.5 (*PNPT1*): c.1424A>G: p.(Glu475Gly) (heterozygous). | This work |
| EK81.8 | 293T clone with NM_033109.5 (*PNPT1*): c.1424A>G: p.(Glu475Gly). | This work |
| P35 | 293T clone with NM_033109.5 (*PNPT1*): c.418C>T (homozygous), c.419C>T (heterozygous): p.(Pro140Ser/Leu) (compound heterozygous). | This work |
| P102 | 293T clone with NM_033109.5 (*PNPT1*): c.418C>T: p.(Pro140Ser). | This work |

DNA and **RNA** Oligonucleotides

| **Name** | **Sequence (5'-3')** |
|---|---|
| 5a | TCCAAGGGGATTTTTCTTGA |
| 5b | AATTTTTGATTGATCTGGCAACT |
| 16Sa | TGTCGTCAGCTCGTGTCGTGA |
| 16Sb | ATCCCCACCTTCCTCCGGT |
| 17a | TGAAATTGGCAAAGTCACTGG |
| 17b | CTCCTGCCTAATGATCACTCTAT |
| ACTa | CCTGGCACCCAGCACAAT |
| ACTb | GGGCCGGACTCGTCATAC |
| CO-1a | ACAGACCGCAACCTCAACAC |
| CO-1b | GTGTGCTCACACGATAAACCC |
| DNA20 | ACTGGACAAATACTCCGACG |
| GLa | TGCTGTCTCCATGTTTGATGTATCT |
| GLb | TCTCTGCTCCCCACCTCTAAGT |
| pFYF1320_ L141La | ACCGATTAGATCGCTTTTTCC |
| pFYF1320_ L141Lb | GTTTCGTCCTTTCCACAAG |
| pFYF1320_E475Ga | TAGAGTCAAAGTTTTAGAGCTAGAAATAGCAAG |
| pFYF1320_E475Gb | GGACTTCAGACGGTGTTTCGTCCTTTCC |
| pFYF1320_P140La | TCTTTCCAGCGTTTTAGAGCTAGAAATAGCAAG |

| | |
|---|---|
| pFYF1320_P140Lb | GCGGTCTAATCGGTGTTTCGTCCTTTCC |
| pFYF1320_P140L/Sa | ACCGATTAGATCGCTCTTTCC |
| pFYF1320_P140L/Sb | GTTTCGTCCTTTCCACAAG |
| PNTP1a | GCATGTGGCGGAAGTTTAGC |
| PNTP1b | TCACCCTTCTCAGGATCGGT |
| recA-a | CACTGGGCCAGATTGAGAAAC |
| recA-b | CCATCGGCAGACCACCTG |
| **RNA20** | **ACUGGACAAAUACUCCGACG** |
| UURa | CACCCAAGAACAGGGTTTGT |
| UURb | TGGCCATGGGTATGTTGTTA |
