## Supplementary Table S2 for "Pathological PNPase variants with altered RNA binding and degradation activity affect the phenotype of bacterial and human cell models"

**Supplementary Table S2. List of differentially expressed genes**

1. ***Samples not incubated with staurosporine***

| E475G heterozygous cell line *vs.* wt | | | | | |
|---|---|---|---|---|---|
| | **Gene** | **logFC** | **logCPM** | **PValue** | **FDR** |
| **1** | ACHE | 1.47 | 0.98 | 7.82E-08 | 6.13E-06 |
| **2** | ACSM3 | -1.00 | 1.44 | 2.51E-04 | 5.76E-03 |
| **3** | ADAMTS12 | -3.38 | -1.18 | 3.57E-05 | 1.16E-03 |
| **4** | ANXA1 | -2.15 | 2.16 | 3.87E-15 | 1.23E-12 |
| **5** | APOBEC3B | 1.30 | 1.37 | 6.01E-05 | 1.79E-03 |
| **6** | ARC | 1.98 | 2.26 | 5.34E-19 | 2.68E-16 |
| **7** | ARL4C | -1.01 | 3.23 | 7.43E-07 | 4.29E-05 |
| **8** | BCHE | -1.18 | 2.95 | 1.61E-14 | 4.74E-12 |
| **9** | BMP2 | -1.36 | 0.39 | 8.04E-06 | 3.39E-04 |
| **10** | BMP3 | -1.40 | 0.10 | 9.49E-06 | 3.87E-04 |
| **11** | C11orf96 | 1.70 | 3.02 | 3.81E-07 | 2.45E-05 |
| **12** | C6orf132 | -2.61 | -0.34 | 4.68E-09 | 5.03E-07 |
| **13** | C8orf44-SGK3 | -6.71 | 0.30 | 2.07E-04 | 4.93E-03 |
| **14** | CASP10 | -1.84 | 0.80 | 2.64E-14 | 7.10E-12 |
| **15** | CCDC149 | -1.22 | -0.54 | 2.28E-04 | 5.34E-03 |
| **16** | CD68 | 1.03 | 1.88 | 1.74E-04 | 4.27E-03 |
| **17** | CDH12 | 2.26 | 1.95 | 8.50E-21 | 5.30E-18 |
| **18** | CDHR1 | 1.42 | 2.99 | 2.14E-17 | 1.01E-14 |
| **19** | CGA | 2.47 | 2.03 | 1.06E-17 | 5.18E-15 |
| **20** | CHRDL1 | -2.42 | -0.68 | 8.14E-08 | 6.35E-06 |
| **21** | COL12A1 | -1.10 | 2.36 | 1.12E-05 | 4.48E-04 |
| **22** | COL5A1 | -1.37 | 4.40 | 1.45E-28 | 2.05E-25 |
| **23** | CPNE7 | -3.81 | 0.44 | 1.71E-19 | 9.43E-17 |
| **24** | CPS1 | -1.15 | 5.72 | 9.06E-23 | 7.43E-20 |
| **25** | CPT1C | 1.12 | 1.18 | 2.74E-05 | 9.36E-04 |
| **26** | CPXM1 | 1.33 | 0.45 | 1.70E-04 | 4.21E-03 |
| **27** | CRTAC1 | 1.26 | 0.53 | 1.15E-04 | 3.00E-03 |
| **28** | CYB561 | 1.44 | 1.83 | 1.08E-06 | 5.94E-05 |
| **29** | CYBB | 1.86 | 0.07 | 9.94E-06 | 4.01E-04 |
| **30** | CYRIA | -3.35 | -1.49 | 1.79E-07 | 1.27E-05 |
| **31** | DCHS2 | -1.50 | 1.33 | 2.47E-08 | 2.20E-06 |
| **32** | DECR1 | -1.11 | 0.92 | 3.56E-04 | 7.69E-03 |
| **33** | DHRS2 | 1.17 | 1.96 | 2.09E-05 | 7.57E-04 |
| **34** | DMRT2 | -2.41 | -1.06 | 5.67E-06 | 2.47E-04 |
| **35** | DMRT3 | -1.40 | 0.29 | 7.08E-05 | 2.03E-03 |
| **36** | DNAH17 | 1.75 | 2.75 | 7.27E-17 | 3.06E-14 |
| **37** | DNAH17-AS1 | 1.76 | -0.62 | 1.56E-04 | 3.93E-03 |
| **38** | EEF1A2 | 1.51 | 7.12 | 1.72E-30 | 4.46E-27 |
| **39** | EFEMP1 | -1.39 | 1.74 | 7.64E-09 | 7.94E-07 |
| **40** | EGR3 | 1.24 | 0.59 | 4.18E-06 | 1.92E-04 |
| **41** | EMP3 | 1.00 | 4.62 | 2.25E-09 | 2.58E-07 |
| **42** | EPHA4 | -1.10 | 3.74 | 2.27E-15 | 7.39E-13 |

| | | | | | |
|---|---|---|---|---|---|
| **43** | EPHB1 | -1.19 | 1.76 | 3.63E-08 | 3.05E-06 |
| **44** | EPPK1 | -1.01 | 2.22 | 4.61E-04 | 9.44E-03 |
| **45** | ERVMER34-1 | -4.10 | -0.45 | 2.92E-21 | 1.98E-18 |
| **46** | ETV1 | -1.20 | 2.95 | 6.35E-16 | 2.36E-13 |
| **47** | FBXL7 | -1.55 | 0.82 | 4.13E-05 | 1.30E-03 |
| **48** | FGF13 | -1.31 | 1.02 | 9.12E-10 | 1.17E-07 |
| **49** | FOXN4 | 1.40 | 1.00 | 1.57E-06 | 8.39E-05 |
| **50** | GABRQ | -1.67 | -0.11 | 2.12E-05 | 7.64E-04 |
| **51** | GADD45B | 1.46 | 3.23 | 2.45E-14 | 6.71E-12 |
| **52** | GALNT14 | -1.65 | 0.59 | 6.81E-11 | 1.09E-08 |
| **53** | GPR50 | 1.68 | 1.39 | 4.52E-06 | 2.04E-04 |
| **54** | GRIA4 | -1.67 | 0.68 | 6.54E-05 | 1.90E-03 |
| **55** | H1-1 | 1.09 | 4.09 | 7.19E-10 | 9.44E-08 |
| **56** | H2BC10 | 2.66 | -0.69 | 6.89E-06 | 2.94E-04 |
| **57** | H3C11 | 1.64 | 3.22 | 5.70E-10 | 7.87E-08 |
| **58** | HEIH | 1.56 | -0.92 | 2.52E-04 | 5.76E-03 |
| **59** | HLA-DMB | -1.58 | -0.48 | 5.06E-05 | 1.54E-03 |
| **60** | HLA-DOA | -2.20 | 0.71 | 2.34E-10 | 3.55E-08 |
| **61** | HMOX1 | 1.46 | 3.44 | 2.62E-13 | 5.92E-11 |
| **62** | HOXA10-HOXA9 | -1.44 | 1.49 | 4.40E-10 | 6.24E-08 |
| **63** | HOXB-AS3 | -1.13 | 0.02 | 8.87E-05 | 2.44E-03 |
| **64** | IFI44L | -1.75 | -0.10 | 5.34E-07 | 3.26E-05 |
| **65** | IGF2 | 2.39 | 3.99 | 2.51E-19 | 1.31E-16 |
| **66** | IGFBP3 | -1.79 | -0.99 | 1.44E-06 | 7.80E-05 |
| **67** | IGFBP5 | -2.06 | 3.31 | 4.94E-30 | 9.62E-27 |
| **68** | IL11 | 3.12 | -0.81 | 9.25E-10 | 1.17E-07 |
| **69** | IL12RB2 | 3.08 | 3.53 | 1.29E-22 | 1.00E-19 |
| **70** | IL15RA | 1.22 | 1.41 | 4.23E-05 | 1.32E-03 |
| **71** | IL6R | 1.58 | 3.16 | 5.58E-25 | 5.80E-22 |
| **72** | KCNS3 | -1.54 | 0.47 | 4.52E-10 | 6.30E-08 |
| **73** | KLHL41 | -1.23 | 1.07 | 5.92E-05 | 1.77E-03 |
| **74** | KRT17 | 2.69 | -0.75 | 9.55E-06 | 3.88E-04 |
| **75** | LAMB3 | 1.61 | -0.05 | 1.45E-04 | 3.69E-03 |
| **76** | LAPTM5 | 2.42 | -0.56 | 2.23E-08 | 2.02E-06 |
| **77** | LGR5 | -1.13 | 4.66 | 1.75E-19 | 9.43E-17 |
| **78** | LINC01234 | 2.01 | 0.52 | 9.12E-08 | 7.01E-06 |
| **79** | LINGO2 | 1.10 | 3.17 | 1.20E-07 | 8.98E-06 |
| **80** | LOC100128288 | 5.90 | 0.54 | 6.94E-05 | 2.00E-03 |
| **81** | LOC100130992 | 1.47 | -0.25 | 2.28E-04 | 5.33E-03 |
| **82** | LOC124908427 | -2.01 | -0.07 | 1.60E-04 | 4.03E-03 |
| **83** | LRRC24 | 3.58 | -0.50 | 2.16E-04 | 5.09E-03 |
| **84** | LRRTM4 | -4.64 | -1.30 | 1.31E-11 | 2.40E-09 |
| **85** | LSMEM1 | 1.88 | 0.92 | 2.79E-05 | 9.52E-04 |
| **86** | MAGEB2 | 2.45 | 4.31 | 1.08E-59 | 8.38E-56 |
| **87** | MAL2 | -1.97 | 1.18 | 6.89E-14 | 1.79E-11 |
| **88** | MAP2 | -1.38 | 1.39 | 2.87E-10 | 4.25E-08 |

| | | | | | |
|---|---|---|---|---|---|
| **89** | MAP2K6 | -1.07 | 3.90 | 1.64E-14 | 4.74E-12 |
| **90** | MFAP2 | -4.45 | 2.08 | 1.03E-54 | 5.35E-51 |
| **91** | MGAM | 1.53 | 4.00 | 6.90E-05 | 1.99E-03 |
| **92** | MME | -1.31 | 2.89 | 3.10E-14 | 8.18E-12 |
| **93** | NEFM | 2.22 | 9.65 | 1.21E-23 | 1.05E-20 |
| **94** | NMNAT2 | 1.73 | 0.48 | 2.75E-06 | 1.34E-04 |
| **95** | NPR1 | -1.76 | -0.32 | 3.56E-04 | 7.69E-03 |
| **96** | NR0B1 | 1.63 | -1.07 | 3.67E-06 | 1.72E-04 |
| **97** | NRXN3 | -1.14 | 1.61 | 3.77E-05 | 1.21E-03 |
| **98** | OAS3 | -1.12 | 2.09 | 3.40E-08 | 2.88E-06 |
| **99** | OSTN | -1.59 | 2.26 | 1.38E-11 | 2.50E-09 |
| **100** | OXCT1 | -1.65 | 2.96 | 1.18E-20 | 7.08E-18 |
| **101** | P2RX5 | 1.38 | 2.06 | 6.42E-11 | 1.05E-08 |
| **102** | P4HTM | -2.37 | 1.63 | 1.24E-11 | 2.30E-09 |
| **103** | PAK3 | -1.58 | 0.37 | 2.92E-07 | 1.95E-05 |
| **104** | PAPPA | -2.42 | 0.54 | 6.38E-07 | 3.79E-05 |
| **105** | PAX1 | -2.23 | -1.60 | 1.24E-04 | 3.21E-03 |
| **106** | PCYT1B | 1.36 | 4.47 | 7.61E-27 | 9.89E-24 |
| **107** | PDE3A | -3.17 | -0.99 | 1.67E-07 | 1.22E-05 |
| **108** | PDLIM4 | -1.27 | 0.62 | 2.38E-05 | 8.39E-04 |
| **109** | PER2 | -1.14 | 3.91 | 1.22E-21 | 8.64E-19 |
| **110** | PLAGL1 | -2.04 | 1.36 | 2.16E-11 | 3.82E-09 |
| **111** | PLCL1 | 3.07 | 3.98 | 6.77E-118 | 1.06E-113 |
| **112** | PLPPR4 | -1.10 | 0.35 | 4.16E-05 | 1.31E-03 |
| **113** | PLXDC2 | -2.44 | -0.12 | 2.57E-08 | 2.25E-06 |
| **114** | PPP1R15A | 1.10 | 4.18 | 2.18E-08 | 1.99E-06 |
| **115** | PRAME | 1.03 | 3.80 | 1.35E-07 | 9.89E-06 |
| **116** | PRELID1P1 | -1.86 | -0.59 | 4.78E-04 | 9.75E-03 |
| **117** | RELN | -2.86 | 0.31 | 8.11E-16 | 2.94E-13 |
| **118** | RGS7BP | -2.73 | -0.57 | 1.82E-14 | 5.08E-12 |
| **119** | RNA45SN1 | -1.40 | 7.08 | 7.68E-06 | 3.26E-04 |
| **120** | RNA45SN2 | -1.17 | 10.31 | 8.82E-05 | 2.44E-03 |
| **121** | ROR2 | -1.75 | 0.64 | 3.95E-07 | 2.53E-05 |
| **122** | RPRM | -1.32 | 2.46 | 2.53E-13 | 5.81E-11 |
| **123** | RYR1 | 1.01 | 1.78 | 5.12E-06 | 2.27E-04 |
| **124** | SAT1 | 1.89 | 5.04 | 3.72E-26 | 4.46E-23 |
| **125** | SELENOP | -1.11 | 4.79 | 4.51E-17 | 2.01E-14 |
| **126** | SEMA6A | -1.22 | 3.26 | 6.32E-16 | 2.36E-13 |
| **127** | SERPINB8 | 1.04 | 0.30 | 2.73E-04 | 6.18E-03 |
| **128** | SERPINF1 | -1.22 | 1.52 | 4.60E-09 | 4.98E-07 |
| **129** | SFRP1 | -2.74 | 1.93 | 7.05E-30 | 1.10E-26 |
| **130** | SFRP2 | 2.42 | 2.69 | 5.68E-22 | 4.22E-19 |
| **131** | SH2D3C | 1.46 | 0.91 | 6.22E-06 | 2.69E-04 |
| **132** | SLC13A3 | 1.89 | 1.17 | 4.59E-08 | 3.78E-06 |
| **133** | SLC19A3 | -3.35 | -0.58 | 9.64E-14 | 2.38E-11 |
| **134** | SLFN12 | -4.15 | 0.33 | 3.87E-33 | 1.21E-29 |

| | | | | | |
|---|---|---|---|---|---|
| **135** | SLIT1 | 1.06 | 3.64 | 9.84E-10 | 1.24E-07 |
| **136** | SMARCA1 | -1.45 | 4.32 | 9.19E-26 | 1.02E-22 |
| **137** | SNAI2 | -1.06 | 4.06 | 1.77E-08 | 1.65E-06 |
| **138** | SORBS2 | -1.46 | 0.68 | 1.91E-05 | 7.02E-04 |
| **139** | SOX11 | -3.23 | 0.16 | 1.06E-14 | 3.23E-12 |
| **140** | SP8 | -1.03 | 1.27 | 2.75E-04 | 6.21E-03 |
| **141** | SPARC | -3.93 | 2.59 | 7.56E-53 | 2.95E-49 |
| **142** | SPOCK1 | -1.52 | 0.07 | 3.55E-06 | 1.68E-04 |
| **143** | SPRY4 | -1.57 | 1.09 | 1.48E-09 | 1.76E-07 |
| **144** | SRGAP3 | -1.35 | 1.88 | 6.17E-10 | 8.43E-08 |
| **145** | STING1 | -1.59 | 2.29 | 1.80E-14 | 5.08E-12 |
| **146** | SULT1C4 | -3.19 | 0.17 | 7.84E-11 | 1.25E-08 |
| **147** | SYN3 | 1.26 | 2.23 | 1.37E-09 | 1.68E-07 |
| **148** | SYNDIG1 | -2.96 | -1.03 | 8.39E-06 | 3.51E-04 |
| **149** | SYT14 | 1.00 | 2.59 | 4.88E-06 | 2.18E-04 |
| **150** | SYTL5 | 1.02 | 2.77 | 3.38E-10 | 4.89E-08 |
| **151** | TAC1 | 1.11 | 1.75 | 1.54E-06 | 8.30E-05 |
| **152** | TFPI2 | 1.09 | 1.59 | 1.02E-05 | 4.11E-04 |
| **153** | THNSL2 | 1.96 | -0.46 | 2.32E-05 | 8.20E-04 |
| **154** | THSD7A | -1.95 | -0.90 | 2.06E-04 | 4.92E-03 |
| **155** | TMEM145 | 1.01 | 1.25 | 1.22E-04 | 3.16E-03 |
| **156** | TMT1A | -1.38 | 3.73 | 8.65E-24 | 8.43E-21 |
| **157** | TNFRSF12A | 1.57 | 3.47 | 2.69E-12 | 5.51E-10 |
| **158** | TNXB | 1.21 | 1.83 | 2.90E-04 | 6.50E-03 |
| **159** | TOR4A | -1.08 | 1.56 | 1.79E-07 | 1.27E-05 |
| **160** | TPM1-AS | -1.54 | -0.65 | 1.05E-04 | 2.80E-03 |
| **161** | TRPA1 | -2.06 | -0.42 | 3.98E-05 | 1.26E-03 |
| **162** | TRPC4 | -1.24 | 0.41 | 5.08E-07 | 3.12E-05 |
| **163** | TUBA4A | -1.73 | 0.57 | 7.39E-05 | 2.10E-03 |
| **164** | UBE2QL1 | -1.24 | 0.63 | 8.56E-05 | 2.38E-03 |
| **165** | UGT3A2 | -1.38 | 1.85 | 1.08E-08 | 1.08E-06 |
| **166** | VGF | 2.63 | 1.33 | 2.16E-30 | 4.81E-27 |
| **167** | VGLL2 | -1.52 | 0.21 | 1.17E-05 | 4.65E-04 |
| **168** | WSCD1 | 1.00 | 4.67 | 5.30E-11 | 8.88E-09 |
| **169** | XK | 1.12 | 3.75 | 2.94E-17 | 1.35E-14 |
| **170** | YWHABP2 | 1.64 | 1.26 | 8.84E-05 | 2.44E-03 |
| **171** | ZDHHC11 | -1.03 | 1.58 | 3.28E-04 | 7.25E-03 |
| **172** | ZNF22 | -3.03 | 1.08 | 5.64E-30 | 9.77E-27 |
| **173** | ZNF365 | 1.07 | 2.20 | 7.25E-08 | 5.71E-06 |

**homozygous E475G line *vs.* wt**

| | Gene | logFC | logCPM | PValue | FDR |
|---|---|---|---|---|---|
| **1** | ABCA3 | 1.20 | 6.19 | 5.39E-28 | 2.47E-25 |
| **2** | ACHE | 1.70 | 0.98 | 5.27E-10 | 2.83E-08 |
| **3** | ACTA2 | -1.16 | 3.74 | 2.78E-17 | 4.17E-15 |
| **4** | ADAM12 | -2.18 | -0.42 | 9.99E-05 | 1.50E-03 |
| **5** | ADAMTS12 | -5.19 | -1.18 | 6.13E-08 | 2.24E-06 |

| | | | | | |
|---|---|---|---|---|---|
| **6** | ADAMTS16 | -4.08 | 1.48 | 9.93E-47 | 9.67E-44 |
| **7** | ADCYAP1R1 | -1.56 | 2.03 | 7.09E-18 | 1.12E-15 |
| **8** | AFF2 | -1.16 | 4.91 | 3.12E-14 | 3.10E-12 |
| **9** | ALPK3 | -1.59 | -0.58 | 5.57E-04 | 6.11E-03 |
| **10** | AMBN | -2.43 | 1.14 | 1.66E-17 | 2.56E-15 |
| **11** | AMH | 1.07 | 3.79 | 2.97E-06 | 7.01E-05 |
| **12** | ANXA1 | -2.34 | 2.16 | 2.17E-17 | 3.28E-15 |
| **13** | APCDD1 | -1.33 | -0.33 | 6.57E-04 | 7.01E-03 |
| **14** | APOBEC3B | 2.20 | 1.37 | 3.54E-12 | 2.76E-10 |
| **15** | ARC | 1.19 | 2.26 | 1.02E-07 | 3.53E-06 |
| **16** | ARL4C | -1.68 | 3.23 | 9.29E-16 | 1.16E-13 |
| **17** | ASIC3 | 1.29 | -0.25 | 2.54E-05 | 4.62E-04 |
| **18** | ASPHD1 | 1.66 | 0.92 | 1.68E-06 | 4.27E-05 |
| **19** | ATP2B3 | -1.11 | 2.09 | 4.02E-06 | 9.21E-05 |
| **20** | ATP8A2 | -1.04 | 2.16 | 5.95E-07 | 1.73E-05 |
| **21** | B3GALT5 | -4.87 | -0.77 | 4.13E-11 | 2.70E-09 |
| **22** | B4GALNT1 | 1.32 | 4.05 | 1.33E-27 | 5.75E-25 |
| **23** | BCHE | -1.33 | 2.95 | 6.36E-18 | 1.02E-15 |
| **24** | BCL6B | 1.35 | 1.82 | 1.65E-14 | 1.72E-12 |
| **25** | BHLHA15 | 1.32 | 1.24 | 8.19E-07 | 2.27E-05 |
| **26** | BMP2 | -2.36 | 0.39 | 3.14E-13 | 2.74E-11 |
| **27** | BORCS8-MEF2B | 1.36 | 2.05 | 9.40E-07 | 2.55E-05 |
| **28** | BRSK2 | 1.23 | 4.31 | 1.05E-18 | 1.88E-16 |
| **29** | C11orf96 | 1.45 | 3.02 | 1.50E-05 | 2.93E-04 |
| **30** | C2CD4A | 1.44 | -0.82 | 1.97E-05 | 3.70E-04 |
| **31** | C6orf132 | -3.75 | -0.34 | 1.22E-13 | 1.12E-11 |
| **32** | CA2 | -1.74 | 6.37 | 4.57E-23 | 1.34E-20 |
| **33** | CALCB | 1.05 | 1.62 | 5.54E-04 | 6.08E-03 |
| **34** | CALHM2 | 1.24 | 2.93 | 2.22E-12 | 1.77E-10 |
| **35** | CARF | -1.19 | 1.05 | 4.70E-06 | 1.06E-04 |
| **36** | CASKIN1 | 1.20 | 3.42 | 3.17E-12 | 2.48E-10 |
| **37** | CBARP | 1.12 | 2.04 | 2.91E-07 | 8.97E-06 |
| **38** | CCDC149 | -1.88 | -0.54 | 9.58E-08 | 3.34E-06 |
| **39** | CCN2 | -1.44 | 4.18 | 5.57E-17 | 8.20E-15 |
| **40** | CCN3 | 2.03 | 2.12 | 6.06E-27 | 2.50E-24 |
| **41** | CD40 | 2.71 | 0.11 | 3.16E-06 | 7.42E-05 |
| **42** | CDH12 | 2.38 | 1.95 | 2.20E-23 | 6.71E-21 |
| **43** | CDHR1 | 1.86 | 2.99 | 1.89E-29 | 1.02E-26 |
| **44** | CDX2 | -1.00 | 4.60 | 3.10E-19 | 5.97E-17 |
| **45** | CEACAM19 | 1.07 | 1.25 | 1.38E-04 | 1.98E-03 |
| **46** | CES3 | 1.02 | 2.14 | 1.11E-05 | 2.27E-04 |
| **47** | CFD | 1.07 | 2.01 | 5.94E-05 | 9.54E-04 |
| **48** | CHAC1 | 1.55 | 1.88 | 1.18E-12 | 9.83E-11 |
| **49** | CHRD | 1.29 | 0.69 | 2.32E-05 | 4.29E-04 |
| **50** | CHRDL1 | -8.07 | -0.68 | 6.47E-19 | 1.20E-16 |
| **51** | CLSTN2 | -1.66 | 1.47 | 7.85E-09 | 3.45E-07 |

| | | | | | |
|---|---|---|---|---|---|
| **52** | CNTN4 | -1.37 | 1.60 | 8.43E-07 | 2.33E-05 |
| **53** | COL12A1 | -3.17 | 2.36 | 2.89E-29 | 1.50E-26 |
| **54** | COL26A1 | 1.32 | 2.16 | 4.68E-10 | 2.53E-08 |
| **55** | COL4A4 | 1.02 | 3.11 | 3.17E-09 | 1.52E-07 |
| **56** | COL5A1 | -1.73 | 4.40 | 5.40E-44 | 4.68E-41 |
| **57** | CORO6 | 1.30 | 0.15 | 3.18E-07 | 9.72E-06 |
| **58** | CPNE7 | -3.15 | 0.44 | 1.02E-15 | 1.26E-13 |
| **59** | CPT1B | 1.00 | 1.71 | 1.70E-04 | 2.35E-03 |
| **60** | CPT1C | 1.56 | 1.18 | 2.55E-09 | 1.24E-07 |
| **61** | CRIP2 | -1.65 | 0.68 | 5.74E-07 | 1.67E-05 |
| **62** | CRIP3 | 1.43 | 2.28 | 5.90E-09 | 2.66E-07 |
| **63** | CRYGD | -1.00 | 2.22 | 1.31E-04 | 1.90E-03 |
| **64** | CTSO | -1.10 | 1.22 | 2.05E-04 | 2.74E-03 |
| **65** | CUX2 | -1.26 | 0.38 | 1.24E-04 | 1.83E-03 |
| **66** | CYB561 | 1.58 | 1.83 | 7.58E-08 | 2.71E-06 |
| **67** | CYBB | 1.98 | 0.07 | 1.50E-06 | 3.87E-05 |
| **68** | CYRIA | -5.46 | -1.49 | 2.02E-11 | 1.41E-09 |
| **69** | DDIT4 | 1.40 | 4.72 | 1.19E-11 | 8.65E-10 |
| **70** | DECR1 | -2.12 | 0.92 | 1.11E-10 | 6.68E-09 |
| **71** | DEPDC1-AS1 | -1.45 | -0.48 | 1.36E-04 | 1.96E-03 |
| **72** | DLL1 | 1.14 | 2.85 | 6.57E-13 | 5.57E-11 |
| **73** | DMRT2 | -3.71 | -1.06 | 3.29E-10 | 1.82E-08 |
| **74** | DMRT3 | -1.31 | 0.29 | 1.84E-04 | 2.51E-03 |
| **75** | DNAH17 | 1.24 | 2.75 | 3.63E-09 | 1.70E-07 |
| **76** | DTX3 | -1.85 | 0.84 | 1.59E-08 | 6.46E-07 |
| **77** | DUSP8 | 1.23 | 6.02 | 4.50E-20 | 9.87E-18 |
| **78** | DYSF | -2.00 | -0.03 | 1.43E-05 | 2.81E-04 |
| **79** | EBF1 | -1.40 | -0.03 | 2.38E-04 | 3.09E-03 |
| **80** | EBF4 | -1.05 | 1.60 | 1.39E-04 | 1.99E-03 |
| **81** | EDIL3 | -1.53 | 3.16 | 2.90E-19 | 5.66E-17 |
| **82** | EEF1A2 | 2.00 | 7.12 | 4.77E-51 | 5.72E-48 |
| **83** | EFEMP1 | -1.47 | 1.74 | 5.86E-10 | 3.13E-08 |
| **84** | EGR1 | 1.59 | 4.38 | 6.57E-11 | 4.13E-09 |
| **85** | EGR3 | 1.56 | 0.59 | 2.45E-09 | 1.19E-07 |
| **86** | EMILIN2 | -1.06 | 3.84 | 2.13E-20 | 4.88E-18 |
| **87** | EMP3 | 1.57 | 4.62 | 6.38E-21 | 1.58E-18 |
| **88** | ENO2 | 1.01 | 4.68 | 2.59E-10 | 1.46E-08 |
| **89** | EPHB1 | -2.13 | 1.76 | 9.98E-21 | 2.39E-18 |
| **90** | ERAP2 | 1.30 | 3.57 | 3.88E-23 | 1.16E-20 |
| **91** | ERVMER34-1 | -4.12 | -0.45 | 1.59E-22 | 4.35E-20 |
| **92** | ETS1 | -1.30 | 1.63 | 7.18E-12 | 5.43E-10 |
| **93** | ETV4 | 1.45 | 3.51 | 3.79E-16 | 4.92E-14 |
| **94** | FA2H | 1.49 | 0.21 | 7.51E-05 | 1.17E-03 |
| **95** | FAAH2 | -2.26 | 0.52 | 1.13E-10 | 6.76E-09 |
| **96** | FAM131C | 1.34 | 1.91 | 1.25E-05 | 2.51E-04 |
| **97** | FBXL7 | -4.04 | 0.82 | 7.64E-19 | 1.39E-16 |

| | | | | | |
|---|---|---|---|---|---|
| **98** | FCHO1 | 1.43 | 2.18 | 3.70E-11 | 2.44E-09 |
| **99** | FGF13 | -4.40 | 1.02 | 5.28E-51 | 5.88E-48 |
| **100** | FGFBP3 | 1.19 | 2.40 | 1.63E-12 | 1.35E-10 |
| **101** | FLI1 | -2.41 | 1.54 | 4.83E-23 | 1.40E-20 |
| **102** | FN1 | -1.35 | 4.48 | 9.12E-21 | 2.22E-18 |
| **103** | FNDC1 | -1.74 | 0.56 | 6.41E-08 | 2.33E-06 |
| **104** | FOS | 2.29 | 2.94 | 5.33E-28 | 2.47E-25 |
| **105** | FOSB | 2.83 | 3.28 | 6.83E-81 | 1.77E-77 |
| **106** | FOXC2 | -1.51 | -1.06 | 1.80E-04 | 2.47E-03 |
| **107** | FOXN4 | 2.38 | 1.00 | 4.03E-18 | 6.61E-16 |
| **108** | FRMPD3 | 1.27 | 1.52 | 3.05E-09 | 1.47E-07 |
| **109** | FUOM | 1.09 | 3.50 | 5.58E-07 | 1.63E-05 |
| **110** | FXYD6 | -1.79 | 2.06 | 1.08E-13 | 9.98E-12 |
| **111** | GABRE | -3.68 | -0.08 | 2.33E-16 | 3.07E-14 |
| **112** | GABRQ | -3.73 | -0.11 | 1.28E-15 | 1.54E-13 |
| **113** | GALNT14 | -3.58 | 0.59 | 3.81E-31 | 2.48E-28 |
| **114** | GDPD3 | 1.70 | -0.46 | 1.59E-04 | 2.23E-03 |
| **115** | GFI1 | 1.29 | 2.30 | 7.25E-16 | 9.12E-14 |
| **116** | GFRA1 | 1.33 | 4.17 | 1.91E-30 | 1.10E-27 |
| **117** | GHR | -1.11 | 2.62 | 8.48E-11 | 5.26E-09 |
| **118** | GPR3 | 1.04 | 0.91 | 1.44E-05 | 2.83E-04 |
| **119** | GPR37 | 1.14 | 2.47 | 6.85E-09 | 3.06E-07 |
| **120** | GRIA4 | -1.87 | 0.68 | 1.08E-05 | 2.22E-04 |
| **121** | GRIN2D | 1.09 | 2.21 | 4.70E-09 | 2.16E-07 |
| **122** | GRM8 | -2.81 | 0.01 | 7.11E-08 | 2.58E-06 |
| **123** | GSTO2 | 1.93 | 0.19 | 7.39E-07 | 2.08E-05 |
| **124** | H2BC10 | 2.06 | -0.69 | 5.97E-04 | 6.47E-03 |
| **125** | H3C11 | 1.91 | 3.22 | 5.89E-13 | 5.04E-11 |
| **126** | HAGHL | 1.35 | 2.10 | 6.25E-06 | 1.35E-04 |
| **127** | HBQ1 | 1.40 | 0.69 | 9.10E-05 | 1.38E-03 |
| **128** | HEIH | 1.90 | -0.92 | 3.27E-06 | 7.60E-05 |
| **129** | HES7 | 1.86 | -1.11 | 1.53E-04 | 2.16E-03 |
| **130** | HIC1 | 1.18 | 1.48 | 4.65E-09 | 2.15E-07 |
| **131** | HLA-DOA | -2.09 | 0.71 | 5.42E-10 | 2.91E-08 |
| **132** | HOOK1 | 1.13 | 4.17 | 2.08E-14 | 2.11E-12 |
| **133** | HS3ST5 | 1.91 | -0.06 | 7.99E-08 | 2.83E-06 |
| **134** | HTR1D | 1.48 | 0.00 | 6.50E-05 | 1.04E-03 |
| **135** | ICAM5 | 1.73 | 2.38 | 3.08E-10 | 1.72E-08 |
| **136** | ID2 | -1.24 | 6.16 | 2.70E-31 | 1.91E-28 |
| **137** | IFI30 | 1.52 | 3.70 | 7.15E-24 | 2.37E-21 |
| **138** | IFI35 | 1.79 | 1.50 | 1.64E-10 | 9.48E-09 |
| **139** | IGF2 | -1.52 | 3.99 | 3.55E-08 | 1.34E-06 |
| **140** | IGFBP3 | -2.68 | -0.99 | 2.39E-11 | 1.64E-09 |
| **141** | IGFBP5 | -1.45 | 3.31 | 1.09E-16 | 1.53E-14 |
| **142** | IGSF11 | -2.14 | 0.11 | 1.03E-08 | 4.43E-07 |
| **143** | IKZF1 | -2.44 | 0.42 | 9.33E-05 | 1.41E-03 |

| | | | | | |
|---|---|---|---|---|---|
| **144** | IKZF3 | 1.11 | 3.91 | 3.87E-21 | 9.73E-19 |
| **145** | IL12RB2 | 3.07 | 3.53 | 1.56E-22 | 4.34E-20 |
| **146** | IL15RA | 1.51 | 1.41 | 3.85E-07 | 1.16E-05 |
| **147** | IL6R | 1.17 | 3.16 | 2.17E-14 | 2.18E-12 |
| **148** | IRF5 | 1.13 | 2.54 | 2.66E-08 | 1.04E-06 |
| **149** | ITGB8 | -1.08 | 0.43 | 8.23E-06 | 1.74E-04 |
| **150** | ITPR3 | 1.32 | 4.68 | 2.52E-25 | 9.82E-23 |
| **151** | JADE2 | 1.01 | 1.74 | 7.34E-09 | 3.24E-07 |
| **152** | JAG2 | 1.03 | 5.58 | 5.35E-09 | 2.44E-07 |
| **153** | JAKMIP2 | -1.20 | 2.68 | 1.11E-05 | 2.27E-04 |
| **154** | JDP2 | 1.37 | 2.02 | 2.10E-07 | 6.78E-06 |
| **155** | KCNA3 | -1.98 | 0.69 | 7.20E-07 | 2.03E-05 |
| **156** | KCNQ4 | 1.37 | 2.31 | 5.37E-09 | 2.44E-07 |
| **157** | KCNS3 | -2.10 | 0.47 | 5.14E-16 | 6.57E-14 |
| **158** | KIFC2 | 1.00 | 1.90 | 5.23E-05 | 8.59E-04 |
| **159** | KLF7 | 1.72 | 1.35 | 1.50E-10 | 8.73E-09 |
| **160** | KRT222 | -2.35 | -0.68 | 1.25E-08 | 5.23E-07 |
| **161** | L3MBTL4 | 1.13 | 1.99 | 2.13E-08 | 8.43E-07 |
| **162** | LANCL3 | 1.14 | 0.28 | 2.71E-04 | 3.40E-03 |
| **163** | LAPTM5 | -3.54 | -0.56 | 8.89E-06 | 1.87E-04 |
| **164** | LINC00839 | 1.14 | 2.60 | 1.58E-11 | 1.12E-09 |
| **165** | LINC01234 | 1.99 | 0.52 | 1.24E-07 | 4.20E-06 |
| **166** | LINC01551 | 1.00 | 0.54 | 1.52E-04 | 2.15E-03 |
| **167** | LINC02043 | 1.05 | -0.11 | 5.39E-04 | 5.95E-03 |
| **168** | LINC02983 | 1.18 | 0.33 | 1.54E-04 | 2.17E-03 |
| **169** | LINGO1 | 1.07 | 3.79 | 2.13E-15 | 2.46E-13 |
| **170** | LINGO2 | 1.22 | 3.17 | 2.82E-09 | 1.36E-07 |
| **171** | LOC105377622 | 1.28 | 1.36 | 1.79E-08 | 7.24E-07 |
| **172** | LOC90246 | -1.03 | 0.96 | 3.90E-04 | 4.59E-03 |
| **173** | LPAR2 | 1.32 | -0.33 | 8.76E-04 | 8.94E-03 |
| **174** | LRP2 | 1.23 | 4.02 | 1.11E-10 | 6.68E-09 |
| **175** | LRRC24 | 4.19 | -0.50 | 8.76E-06 | 1.84E-04 |
| **176** | LRRTM4 | -8.23 | -1.30 | 1.12E-15 | 1.36E-13 |
| **177** | LSAMP | -2.54 | -0.35 | 1.44E-06 | 3.74E-05 |
| **178** | LTBP4 | 1.12 | 5.53 | 1.93E-09 | 9.57E-08 |
| **179** | MAGEB2 | 2.29 | 4.31 | 1.18E-52 | 1.67E-49 |
| **180** | MAL2 | -5.06 | 1.18 | 2.32E-46 | 2.13E-43 |
| **181** | MAMLD1 | 1.21 | 2.03 | 4.16E-11 | 2.72E-09 |
| **182** | MAP2 | -2.89 | 1.39 | 5.97E-31 | 3.72E-28 |
| **183** | MAP3K15 | 1.68 | 3.28 | 3.48E-31 | 2.36E-28 |
| **184** | MAP6 | -1.45 | 0.42 | 1.58E-04 | 2.22E-03 |
| **185** | MATK | 1.79 | 4.37 | 2.69E-10 | 1.51E-08 |
| **186** | MELTF-AS1 | 1.54 | 0.40 | 3.79E-05 | 6.52E-04 |
| **187** | MFAP2 | -8.04 | 2.08 | 1.07E-82 | 3.33E-79 |
| **188** | MICA-AS1 | 1.24 | -0.30 | 1.62E-05 | 3.14E-04 |
| **189** | MKX | 1.77 | 3.04 | 3.57E-41 | 2.93E-38 |

| | | | | | |
|---|---|---|---|---|---|
| **190** | MLXIPL | 1.45 | 3.28 | 2.39E-16 | 3.14E-14 |
| **191** | MME | -2.01 | 2.89 | 7.12E-29 | 3.47E-26 |
| **192** | MYB | 1.08 | 1.36 | 2.11E-07 | 6.79E-06 |
| **193** | NEFM | 2.59 | 9.65 | 1.08E-30 | 6.50E-28 |
| **194** | NFATC4 | 1.07 | 3.63 | 2.12E-22 | 5.70E-20 |
| **195** | NGEF | -1.26 | 2.27 | 1.49E-06 | 3.85E-05 |
| **196** | NHS | 1.39 | 4.76 | 1.28E-14 | 1.37E-12 |
| **197** | NIBAN1 | 1.45 | 2.37 | 5.04E-14 | 4.91E-12 |
| **198** | NIPAL2 | -1.04 | 3.59 | 6.64E-11 | 4.15E-09 |
| **199** | NKX3-1 | -1.18 | 2.22 | 1.70E-07 | 5.56E-06 |
| **200** | NLRC5 | 1.33 | 2.53 | 1.05E-15 | 1.29E-13 |
| **201** | NMNAT2 | 2.48 | 0.48 | 4.63E-12 | 3.54E-10 |
| **202** | NOS2 | 1.24 | 1.78 | 4.22E-08 | 1.58E-06 |
| **203** | NOXA1 | 1.30 | 0.07 | 2.21E-04 | 2.91E-03 |
| **204** | NPR1 | -2.20 | -0.32 | 1.93E-05 | 3.64E-04 |
| **205** | NPTXR | -1.02 | 2.46 | 3.51E-06 | 8.13E-05 |
| **206** | NRBP2 | 1.18 | 2.25 | 2.62E-10 | 1.48E-08 |
| **207** | NSUN5P1 | 1.06 | 3.25 | 6.04E-08 | 2.22E-06 |
| **208** | NSUN5P2 | 1.33 | 2.38 | 5.50E-05 | 8.95E-04 |
| **209** | NUP153-AS1 | 1.09 | 0.17 | 2.68E-04 | 3.36E-03 |
| **210** | OPRD1 | -1.24 | 3.37 | 1.10E-20 | 2.59E-18 |
| **211** | OPTN | 1.30 | 2.55 | 6.77E-06 | 1.45E-04 |
| **212** | OSTN | -3.16 | 2.26 | 4.57E-33 | 3.39E-30 |
| **213** | OXCT1 | -1.87 | 2.96 | 1.44E-25 | 5.74E-23 |
| **214** | P2RX5 | 1.39 | 2.06 | 3.54E-11 | 2.37E-09 |
| **215** | P4HTM | -2.83 | 1.63 | 3.30E-15 | 3.75E-13 |
| **216** | PAK3 | -2.97 | 0.37 | 3.96E-18 | 6.57E-16 |
| **217** | PANX2 | 1.21 | 2.12 | 3.70E-09 | 1.73E-07 |
| **218** | PAPPA | -2.77 | 0.54 | 3.03E-08 | 1.16E-06 |
| **219** | PARP12 | 1.12 | 2.87 | 3.96E-11 | 2.61E-09 |
| **220** | PAX1 | -3.88 | -1.60 | 8.26E-09 | 3.61E-07 |
| **221** | PCDH19 | -1.42 | 1.55 | 9.09E-12 | 6.78E-10 |
| **222** | PCDHB2 | -1.71 | -1.08 | 4.03E-04 | 4.70E-03 |
| **223** | PCK2 | 1.40 | 4.11 | 4.77E-11 | 3.07E-09 |
| **224** | PCYT1B | 1.29 | 4.47 | 3.29E-24 | 1.14E-21 |
| **225** | PDE3A | -6.94 | -0.99 | 5.33E-15 | 5.85E-13 |
| **226** | PDK4 | 1.46 | 0.61 | 3.83E-06 | 8.84E-05 |
| **227** | PGGHG | 1.08 | 2.37 | 2.53E-04 | 3.23E-03 |
| **228** | PHF21B | 1.25 | 1.31 | 2.13E-07 | 6.83E-06 |
| **229** | PIK3AP1 | 1.13 | 3.65 | 1.37E-19 | 2.85E-17 |
| **230** | PILRB | 1.51 | 1.23 | 4.29E-04 | 4.94E-03 |
| **231** | PIR | 1.07 | 3.68 | 7.55E-17 | 1.10E-14 |
| **232** | PLAGL1 | -2.86 | 1.36 | 1.18E-18 | 2.05E-16 |
| **233** | PLCL1 | 2.86 | 3.98 | 4.00E-102 | 2.08E-98 |
| **234** | PLIN2 | 1.16 | 1.90 | 2.05E-08 | 8.17E-07 |
| **235** | PLPP4 | -1.97 | 0.60 | 3.01E-08 | 1.15E-06 |

| | | | | | |
|---|---|---|---|---|---|
| **236** | PLXDC2 | -7.42 | -0.12 | 8.07E-24 | 2.58E-21 |
| **237** | PNMA2 | 1.05 | 5.30 | 2.53E-20 | 5.64E-18 |
| **238** | PODXL | 1.03 | 6.43 | 5.13E-17 | 7.62E-15 |
| **239** | PPP1R1A | 1.14 | 0.34 | 4.60E-05 | 7.66E-04 |
| **240** | PRAME | 1.51 | 3.80 | 1.42E-14 | 1.50E-12 |
| **241** | PRKAG2-AS1 | 1.03 | 1.16 | 2.16E-04 | 2.86E-03 |
| **242** | PRKCB | -1.97 | 1.88 | 1.57E-18 | 2.65E-16 |
| **243** | PRRT4 | 1.10 | 2.65 | 3.08E-09 | 1.47E-07 |
| **244** | PTN | -6.01 | -1.41 | 1.99E-04 | 2.67E-03 |
| **245** | PUDP | -11.83 | 2.84 | 2.27E-132 | 1.77E-128 |
| **246** | PWWP2B | 1.04 | 3.30 | 6.43E-04 | 6.87E-03 |
| **247** | RAB26 | 1.19 | 1.10 | 3.39E-04 | 4.07E-03 |
| **248** | RASSF2 | 1.35 | 2.78 | 1.18E-11 | 8.62E-10 |
| **249** | RASSF4 | 1.21 | 1.17 | 2.47E-07 | 7.84E-06 |
| **250** | RBP7 | 1.21 | 0.48 | 4.55E-04 | 5.21E-03 |
| **251** | RELN | -5.44 | 0.31 | 9.68E-30 | 5.39E-27 |
| **252** | REPIN1-AS1 | 1.10 | 1.15 | 1.88E-04 | 2.56E-03 |
| **253** | RGS7BP | -3.79 | -0.57 | 1.12E-21 | 2.92E-19 |
| **254** | RHOF | 1.32 | 1.89 | 2.48E-07 | 7.85E-06 |
| **255** | RHPN1 | 1.04 | 3.22 | 4.27E-05 | 7.20E-04 |
| **256** | RIMS3 | 1.15 | 4.10 | 8.81E-17 | 1.26E-14 |
| **257** | RNF180 | -3.82 | -0.39 | 1.86E-14 | 1.91E-12 |
| **258** | ROR2 | -1.51 | 0.64 | 9.60E-06 | 1.99E-04 |
| **259** | RPS6KL1 | 1.01 | 3.58 | 1.63E-16 | 2.23E-14 |
| **260** | RTN1 | -1.07 | 3.11 | 2.63E-09 | 1.28E-07 |
| **261** | RYR2 | -1.26 | 3.70 | 1.83E-09 | 9.16E-08 |
| **262** | SBK1 | 1.02 | 2.50 | 6.24E-07 | 1.79E-05 |
| **263** | SCARA5 | 1.15 | 0.72 | 2.28E-04 | 2.99E-03 |
| **264** | SCIN | -1.33 | 0.17 | 7.91E-05 | 1.23E-03 |
| **265** | SCN4B | -1.32 | 1.54 | 3.60E-10 | 1.98E-08 |
| **266** | SELENOP | -1.37 | 4.79 | 5.83E-25 | 2.22E-22 |
| **267** | SEMA6A | -1.24 | 3.26 | 1.97E-16 | 2.65E-14 |
| **268** | SERPINF1 | -1.73 | 1.52 | 1.30E-15 | 1.54E-13 |
| **269** | SFRP1 | -5.65 | 1.93 | 1.92E-69 | 4.27E-66 |
| **270** | SFRP2 | 3.95 | 2.69 | 2.74E-56 | 4.74E-53 |
| **271** | SFXN3 | 1.07 | 4.18 | 2.30E-20 | 5.21E-18 |
| **272** | SH2B2 | 1.01 | 2.42 | 4.23E-08 | 1.58E-06 |
| **273** | SH2D3C | 2.26 | 0.91 | 8.81E-14 | 8.27E-12 |
| **274** | SHANK1 | 1.03 | 0.18 | 8.81E-04 | 8.97E-03 |
| **275** | SLC13A3 | 2.24 | 1.17 | 1.25E-11 | 8.91E-10 |
| **276** | SLC19A3 | -9.70 | -0.58 | 6.51E-29 | 3.27E-26 |
| **277** | SLC30A3 | 1.23 | 1.03 | 1.20E-07 | 4.10E-06 |
| **278** | SLC38A4 | -1.60 | -0.68 | 1.06E-04 | 1.58E-03 |
| **279** | SLC6A11 | 1.01 | 5.26 | 5.36E-24 | 1.82E-21 |
| **280** | SLC6A9 | 1.30 | 3.66 | 1.45E-10 | 8.46E-09 |
| **281** | SLCO3A1 | -1.98 | -0.56 | 1.95E-06 | 4.89E-05 |

| | | | | | |
|---|---|---|---|---|---|
| **282** | SLFN12 | -8.14 | 0.33 | 1.18E-51 | 1.53E-48 |
| **283** | SLIT1 | 1.53 | 3.64 | 7.08E-19 | 1.30E-16 |
| **284** | SMAD6 | -1.07 | 2.97 | 2.43E-10 | 1.39E-08 |
| **285** | SMARCA1 | -2.12 | 4.32 | 2.74E-49 | 2.85E-46 |
| **286** | SMARCD3 | 1.11 | 4.36 | 1.08E-11 | 7.99E-10 |
| **287** | SNAI2 | -1.96 | 4.06 | 8.11E-24 | 2.58E-21 |
| **288** | SNED1 | -1.19 | 1.59 | 2.02E-06 | 5.05E-05 |
| **289** | SNHG9 | 1.68 | 1.18 | 1.61E-04 | 2.25E-03 |
| **290** | SNORD15B | 1.28 | 2.42 | 1.82E-04 | 2.49E-03 |
| **291** | SORBS2 | -1.40 | 0.68 | 4.73E-05 | 7.85E-04 |
| **292** | SOX11 | -5.61 | 0.16 | 6.10E-27 | 2.50E-24 |
| **293** | SOX8 | -1.12 | 1.66 | 4.57E-06 | 1.03E-04 |
| **294** | SPARC | -8.55 | 2.59 | 9.99E-98 | 3.89E-94 |
| **295** | SPOCK1 | -4.11 | 0.07 | 1.33E-20 | 3.09E-18 |
| **296** | SQOR | 1.21 | 0.52 | 4.10E-04 | 4.77E-03 |
| **297** | SRRM3 | 1.00 | 1.10 | 3.18E-06 | 7.43E-05 |
| **298** | STING1 | -1.57 | 2.29 | 3.98E-14 | 3.90E-12 |
| **299** | STS | -9.77 | 3.18 | 4.99E-138 | 7.78E-134 |
| **300** | STXBP2 | 1.03 | 3.33 | 2.11E-07 | 6.78E-06 |
| **301** | SULT1A1 | -1.71 | 0.57 | 2.79E-06 | 6.62E-05 |
| **302** | SULT1C4 | -6.86 | 0.17 | 1.08E-21 | 2.85E-19 |
| **303** | SULT4A1 | 1.44 | 3.56 | 1.96E-19 | 3.91E-17 |
| **304** | SYN3 | 1.03 | 2.23 | 7.03E-07 | 1.99E-05 |
| **305** | SYNDIG1 | -8.43 | -1.03 | 6.44E-13 | 5.49E-11 |
| **306** | SYNGAP1 | 1.04 | 2.93 | 3.72E-07 | 1.13E-05 |
| **307** | TENT5A | -1.22 | 3.72 | 1.93E-19 | 3.90E-17 |
| **308** | TERT | 1.51 | 1.57 | 2.53E-11 | 1.72E-09 |
| **309** | TGFBR3L | 1.08 | 1.67 | 1.09E-06 | 2.92E-05 |
| **310** | THSD7A | -1.98 | -0.90 | 1.85E-04 | 2.52E-03 |
| **311** | TIGD1 | -1.01 | 3.46 | 6.72E-05 | 1.07E-03 |
| **312** | TLR3 | 1.23 | -0.46 | 5.82E-04 | 6.34E-03 |
| **313** | TMC5 | 1.30 | 0.91 | 9.19E-07 | 2.50E-05 |
| **314** | TMEM132E | 1.74 | 0.97 | 7.36E-08 | 2.64E-06 |
| **315** | TMEM145 | 1.08 | 1.25 | 3.26E-05 | 5.76E-04 |
| **316** | TMEM38A | 1.03 | 2.63 | 1.96E-05 | 3.68E-04 |
| **317** | TNC | 1.04 | 9.63 | 1.45E-21 | 3.71E-19 |
| **318** | TNXB | 1.65 | 1.83 | 2.74E-07 | 8.50E-06 |
| **319** | TOMM40P1 | 1.26 | 1.62 | 3.24E-06 | 7.55E-05 |
| **320** | TRIP6 | 1.01 | 7.05 | 1.06E-08 | 4.53E-07 |
| **321** | TSPAN7 | 1.13 | 5.70 | 3.34E-19 | 6.34E-17 |
| **322** | TUBA4A | -1.98 | 0.57 | 7.97E-06 | 1.69E-04 |
| **323** | UBE2QL1 | -2.84 | 0.63 | 2.22E-15 | 2.55E-13 |
| **324** | UCP2 | 1.01 | 3.83 | 7.96E-08 | 2.83E-06 |
| **325** | UGT3A2 | -1.15 | 1.85 | 1.31E-06 | 3.44E-05 |
| **326** | ULBP1 | 1.08 | 2.58 | 2.84E-08 | 1.10E-06 |
| **327** | UNCX | -2.09 | -1.16 | 7.37E-05 | 1.15E-03 |

| | | | | | |
|---|---|---|---|---|---|
| **328** | USP32P2 | -1.29 | 0.33 | 2.17E-04 | 2.86E-03 |
| **329** | USP40 | -1.15 | 4.19 | 2.13E-19 | 4.19E-17 |
| **330** | VAX2 | -1.01 | 0.95 | 1.85E-04 | 2.52E-03 |
| **331** | VGF | 1.46 | 1.33 | 1.83E-09 | 9.16E-08 |
| **332** | VGLL2 | -1.51 | 0.21 | 1.21E-05 | 2.45E-04 |
| **333** | WNT8B | 1.14 | 0.29 | 5.38E-05 | 8.80E-04 |
| **334** | WSCD1 | 1.49 | 4.67 | 1.31E-22 | 3.72E-20 |
| **335** | WWC3 | -2.02 | 4.13 | 4.17E-54 | 6.50E-51 |
| **336** | XK | 1.20 | 3.75 | 1.28E-19 | 2.69E-17 |
| **337** | ZCCHC12 | -2.73 | 2.93 | 2.21E-24 | 7.82E-22 |
| **338** | ZNF22 | -6.83 | 1.08 | 2.44E-68 | 4.75E-65 |
| **339** | ZNF365 | 1.26 | 2.20 | 2.45E-10 | 1.40E-08 |
| **340** | ZNF43 | -5.41 | 0.58 | 3.94E-34 | 3.07E-31 |
| **341** | ZNF469 | -1.47 | 0.35 | 6.94E-04 | 7.33E-03 |
| **342** | ZNF595 | -3.17 | -0.69 | 2.48E-05 | 4.54E-04 |
| **343** | ZNF649 | -1.23 | 2.94 | 3.65E-11 | 2.43E-09 |
| **344** | ZNF883 | -1.38 | 3.05 | 6.01E-12 | 4.57E-10 |
| **345** | ZSWIM8 | 1.00 | 5.62 | 1.75E-11 | 1.24E-09 |
| **Homozygous vs. heterozygous E475G cell line** | | | | | |
| | **Gene** | **logFC** | **logCPM** | **PValue** | **FDR** |
| **1** | ACTA2 | -1.02 | 3.74 | 1.05E-13 | 4.08E-11 |
| **2** | ADAMTS16 | -3.36 | 1.48 | 1.78E-30 | 3.46E-27 |
| **3** | AMBN | -1.51 | 1.14 | 6.47E-07 | 5.34E-05 |
| **4** | ANGPT1 | 1.76 | -0.79 | 2.63E-04 | 7.28E-03 |
| **5** | APLN | 1.38 | -0.80 | 3.25E-04 | 8.59E-03 |
| **6** | ATP2B3 | -1.17 | 2.09 | 1.05E-06 | 8.06E-05 |
| **7** | ATP8A2 | -1.11 | 2.16 | 7.87E-08 | 8.34E-06 |
| **8** | B3GALT5 | -3.51 | -0.77 | 3.43E-06 | 2.22E-04 |
| **9** | B3GAT1 | -1.32 | -0.42 | 2.60E-04 | 7.22E-03 |
| **10** | BMP3 | 2.03 | 0.10 | 3.07E-11 | 6.74E-09 |
| **11** | C8orf44-SGK3 | 8.53 | 0.30 | 7.87E-07 | 6.33E-05 |
| **12** | CA2 | -1.03 | 6.37 | 2.70E-09 | 4.34E-07 |
| **13** | CARF | -1.06 | 1.05 | 5.54E-05 | 2.24E-03 |
| **14** | CASP10 | 1.84 | 0.80 | 1.93E-14 | 7.90E-12 |
| **15** | CCN3 | 1.03 | 2.12 | 4.02E-09 | 6.21E-07 |
| **16** | CCT5P1 | -3.34 | -0.71 | 8.15E-05 | 2.99E-03 |
| **17** | CGA | -2.38 | 2.03 | 8.98E-18 | 5.83E-15 |
| **18** | CHAC1 | 1.43 | 1.88 | 2.96E-11 | 6.59E-09 |
| **19** | CHRDL1 | -5.67 | -0.68 | 2.86E-05 | 1.32E-03 |
| **20** | CLSTN2 | -1.21 | 1.47 | 3.33E-05 | 1.50E-03 |
| **21** | COL12A1 | -2.07 | 2.36 | 2.99E-13 | 1.08E-10 |
| **22** | CORO6 | 1.25 | 0.15 | 8.18E-07 | 6.48E-05 |
| **23** | CRIP3 | 1.27 | 2.28 | 1.86E-07 | 1.81E-05 |
| **24** | DDIT4 | 1.09 | 4.72 | 1.09E-07 | 1.09E-05 |
| **25** | DEPDC1-AS1 | -1.48 | -0.48 | 8.28E-05 | 3.03E-03 |
| **26** | DHRS3 | 1.12 | 0.85 | 5.17E-05 | 2.13E-03 |

| | | | | | |
|---|---|---|---|---|---|
| **27** | DTX3 | -1.48 | 0.84 | 8.63E-06 | 4.77E-04 |
| **28** | DYNC1I1 | -1.36 | 0.17 | 1.38E-05 | 7.07E-04 |
| **29** | EDIL3 | -1.20 | 3.16 | 2.73E-12 | 7.33E-10 |
| **30** | EGFR | 1.05 | 4.59 | 1.39E-17 | 8.68E-15 |
| **31** | EGR1 | 1.04 | 4.38 | 1.40E-05 | 7.14E-04 |
| **32** | EP300-AS1 | 1.12 | 1.34 | 1.79E-05 | 8.88E-04 |
| **33** | ERAP2 | 1.43 | 3.57 | 2.03E-27 | 2.87E-24 |
| **34** | ERBB3 | 1.01 | 1.11 | 5.78E-06 | 3.48E-04 |
| **35** | ETS1 | -1.32 | 1.63 | 2.13E-12 | 5.93E-10 |
| **36** | ETV1 | 1.55 | 2.95 | 4.84E-26 | 5.39E-23 |
| **37** | ETV4 | 1.93 | 3.51 | 3.68E-26 | 4.42E-23 |
| **38** | ETV5 | 1.36 | 4.73 | 1.74E-39 | 4.51E-36 |
| **39** | FAAH2 | -1.95 | 0.52 | 4.43E-08 | 5.11E-06 |
| **40** | FBXL7 | -2.49 | 0.82 | 2.07E-07 | 1.98E-05 |
| **41** | FGF13 | -3.09 | 1.02 | 1.15E-21 | 9.46E-19 |
| **42** | FLI1 | -1.66 | 1.54 | 1.71E-11 | 4.22E-09 |
| **43** | FOS | 2.16 | 2.94 | 1.44E-25 | 1.50E-22 |
| **44** | FOSB | 2.03 | 3.28 | 3.35E-48 | 1.30E-44 |
| **45** | FRMPD3 | 1.16 | 1.52 | 4.74E-08 | 5.39E-06 |
| **46** | FXYD6 | -1.04 | 2.06 | 2.38E-05 | 1.13E-03 |
| **47** | GABRE | -3.33 | -0.08 | 4.84E-13 | 1.61E-10 |
| **48** | GABRQ | -2.06 | -0.11 | 4.16E-05 | 1.80E-03 |
| **49** | GALNT14 | -1.93 | 0.59 | 4.89E-08 | 5.48E-06 |
| **50** | GAP43 | 1.10 | 2.39 | 2.48E-07 | 2.34E-05 |
| **51** | GJB2 | 1.01 | 0.92 | 3.21E-04 | 8.53E-03 |
| **52** | GPR37 | 1.16 | 2.47 | 2.69E-09 | 4.34E-07 |
| **53** | GPR50 | -2.32 | 1.39 | 7.84E-10 | 1.42E-07 |
| **54** | GRID2IP | 1.31 | 0.62 | 3.27E-04 | 8.63E-03 |
| **55** | HAGHL | 1.18 | 2.10 | 5.80E-05 | 2.31E-03 |
| **56** | HELZ2 | 1.15 | 4.11 | 1.13E-05 | 6.02E-04 |
| **57** | HES7 | 1.82 | -1.11 | 1.09E-04 | 3.72E-03 |
| **58** | HMOX1 | -1.06 | 3.44 | 6.12E-08 | 6.67E-06 |
| **59** | HOXA10-HOXA9 | 1.21 | 1.49 | 1.53E-07 | 1.50E-05 |
| **60** | HS3ST5 | 1.39 | -0.06 | 4.58E-05 | 1.92E-03 |
| **61** | HSPA6 | -2.78 | -0.87 | 3.27E-08 | 3.92E-06 |
| **62** | ICAM5 | 1.24 | 2.38 | 3.63E-06 | 2.32E-04 |
| **63** | IFI35 | 1.67 | 1.50 | 2.59E-09 | 4.24E-07 |
| **64** | IFI44L | 1.37 | -0.10 | 1.03E-04 | 3.60E-03 |
| **65** | IFIH1 | 1.54 | 0.44 | 7.46E-09 | 1.07E-06 |
| **66** | IGF2 | -3.91 | 3.99 | 7.43E-42 | 2.32E-38 |
| **67** | ITPR3 | 1.46 | 4.68 | 2.10E-30 | 3.63E-27 |
| **68** | JDP2 | 1.56 | 2.02 | 2.17E-09 | 3.64E-07 |
| **69** | KCNA3 | -1.55 | 0.69 | 1.06E-04 | 3.66E-03 |
| **70** | KCND3 | -1.50 | 1.53 | 6.60E-06 | 3.87E-04 |
| **71** | KIT | 1.09 | 3.88 | 7.02E-25 | 6.84E-22 |
| **72** | KLHL41 | 1.62 | 1.07 | 7.56E-08 | 8.08E-06 |

| | | | | | |
|---|---|---|---|---|---|
| **73** | KRT17 | -2.62 | -0.75 | 7.90E-06 | 4.50E-04 |
| **74** | KRT222 | -1.90 | -0.68 | 5.72E-06 | 3.45E-04 |
| **75** | LAMA4 | 1.02 | 4.81 | 3.09E-27 | 4.01E-24 |
| **76** | LAPTM5 | -5.97 | -0.56 | 2.03E-21 | 1.58E-18 |
| **77** | LOC124908427 | 2.26 | -0.07 | 1.33E-05 | 6.95E-04 |
| **78** | LOC128071547 | 8.63 | -0.38 | 2.92E-06 | 1.94E-04 |
| **79** | LRP2 | 1.27 | 4.02 | 3.31E-11 | 7.16E-09 |
| **80** | MAL2 | -3.09 | 1.18 | 5.74E-15 | 2.49E-12 |
| **81** | MAP2 | -1.52 | 1.39 | 2.27E-08 | 2.88E-06 |
| **82** | MAP6 | -1.58 | 0.42 | 3.77E-05 | 1.66E-03 |
| **83** | MATK | 1.50 | 4.37 | 8.49E-08 | 8.88E-06 |
| **84** | MFAP2 | -3.59 | 2.08 | 1.07E-07 | 1.09E-05 |
| **85** | MIR3648-1 | 2.29 | 2.25 | 4.59E-05 | 1.92E-03 |
| **86** | MKX | 1.01 | 3.04 | 8.07E-16 | 4.34E-13 |
| **87** | MMP24 | -1.20 | 1.43 | 1.29E-08 | 1.78E-06 |
| **88** | MYOM2 | 1.62 | 4.84 | 4.43E-55 | 2.30E-51 |
| **89** | NOTCH3 | 1.14 | 5.14 | 2.15E-18 | 1.52E-15 |
| **90** | NR0B1 | -1.84 | -1.07 | 1.24E-07 | 1.24E-05 |
| **91** | OAS3 | 1.26 | 2.09 | 2.74E-10 | 5.28E-08 |
| **92** | OSTN | -1.57 | 2.26 | 1.46E-08 | 1.97E-06 |
| **93** | PAK3 | -1.39 | 0.37 | 1.02E-04 | 3.58E-03 |
| **94** | PANX2 | 1.13 | 2.12 | 3.57E-08 | 4.22E-06 |
| **95** | PCK2 | 1.01 | 4.11 | 1.48E-06 | 1.09E-04 |
| **96** | PDE1B | 1.17 | 2.89 | 1.27E-11 | 3.25E-09 |
| **97** | PIK3R5 | 1.09 | 0.28 | 2.40E-04 | 6.91E-03 |
| **98** | PLEKHA2 | 1.08 | 2.49 | 1.50E-12 | 4.42E-10 |
| **99** | PLXDC2 | -4.99 | -0.12 | 3.23E-08 | 3.90E-06 |
| **100** | PRKCB | -1.64 | 1.88 | 4.85E-13 | 1.61E-10 |
| **101** | PTPRZ1 | 1.17 | 2.87 | 4.02E-15 | 1.84E-12 |
| **102** | PUDP | -11.77 | 2.84 | 1.59E-130 | 1.24E-126 |
| **103** | RBP7 | 1.30 | 0.48 | 2.55E-04 | 7.11E-03 |
| **104** | RELN | -2.58 | 0.31 | 4.09E-05 | 1.78E-03 |
| **105** | RIMS3 | 1.12 | 4.10 | 4.42E-16 | 2.46E-13 |
| **106** | RNA45SN1 | 2.33 | 7.08 | 5.46E-13 | 1.77E-10 |
| **107** | RNA45SN2 | 1.84 | 10.31 | 1.65E-09 | 2.86E-07 |
| **108** | RNA45SN3 | 2.07 | 2.82 | 5.07E-06 | 3.13E-04 |
| **109** | RNF180 | -2.99 | -0.39 | 7.54E-09 | 1.07E-06 |
| **110** | SAT1 | -1.26 | 5.04 | 4.10E-13 | 1.45E-10 |
| **111** | SCN3B | -1.24 | 0.65 | 9.42E-06 | 5.17E-04 |
| **112** | SERPINB8 | -1.21 | 0.30 | 2.94E-05 | 1.35E-03 |
| **113** | SFRP1 | -2.91 | 1.93 | 4.98E-15 | 2.22E-12 |
| **114** | SFRP2 | 1.53 | 2.69 | 6.13E-13 | 1.95E-10 |
| **115** | SLC19A3 | -6.37 | -0.58 | 1.96E-07 | 1.89E-05 |
| **116** | SLC35D3 | 1.58 | -0.40 | 1.75E-04 | 5.32E-03 |
| **117** | SLC6A9 | 1.00 | 3.66 | 6.17E-07 | 5.12E-05 |
| **118** | SLCO3A1 | -1.70 | -0.56 | 6.32E-05 | 2.44E-03 |

| | | | | | |
|---|---|---|---|---|---|
| **119** | SLFN12 | -4.00 | 0.33 | 5.03E-05 | 2.07E-03 |
| **120** | SMPDL3B | 1.26 | 2.97 | 1.80E-11 | 4.37E-09 |
| **121** | SNORD15B | 1.71 | 2.42 | 1.10E-06 | 8.38E-05 |
| **122** | SOX11 | -2.39 | 0.16 | 1.66E-04 | 5.12E-03 |
| **123** | SPARC | -4.62 | 2.59 | 8.39E-16 | 4.36E-13 |
| **124** | SPOCK1 | -2.59 | 0.07 | 5.45E-07 | 4.54E-05 |
| **125** | SPRY4 | 1.50 | 1.09 | 5.60E-09 | 8.39E-07 |
| **126** | STAT6 | 1.15 | 4.18 | 1.45E-22 | 1.25E-19 |
| **127** | STS | -10.06 | 3.18 | 9.33E-154 | 1.45E-149 |
| **128** | SULT1C4 | -3.67 | 0.17 | 4.76E-05 | 1.97E-03 |
| **129** | TERT | 1.13 | 1.57 | 2.70E-07 | 2.54E-05 |
| **130** | TFPI2 | -1.52 | 1.59 | 1.15E-09 | 2.04E-07 |
| **131** | TMEFF2 | -1.06 | 0.47 | 1.83E-04 | 5.54E-03 |
| **132** | TMEM171 | 1.02 | 0.52 | 1.12E-04 | 3.81E-03 |
| **133** | UBE2QL1 | -1.59 | 0.63 | 2.28E-05 | 1.09E-03 |
| **134** | VCAN | 1.06 | 6.10 | 2.53E-11 | 5.73E-09 |
| **135** | VGF | -1.17 | 1.33 | 5.73E-09 | 8.50E-07 |
| **136** | WDR31 | -1.00 | 2.44 | 1.33E-10 | 2.70E-08 |
| **137** | WWC3 | -1.11 | 4.13 | 1.67E-17 | 1.00E-14 |
| **138** | ZCCHC12 | -3.38 | 2.93 | 1.64E-35 | 3.64E-32 |
| **139** | ZNF22 | -3.81 | 1.08 | 1.94E-15 | 9.43E-13 |
| **140** | ZNF43 | -4.79 | 0.58 | 2.24E-24 | 2.06E-21 |
| **141** | ZNF469 | -1.73 | 0.35 | 6.75E-05 | 2.57E-03 |
| **142** | ZNF66 | -2.06 | -0.35 | 1.18E-04 | 3.96E-03 |

### 2. *Samples incubated with staurosporine*

| E475G heterozygous cell line *vs.* wt | | | | | |
|---|---|---|---|---|---|
| | **Gene** | **logFC** | **logCPM** | **PValue** | **FDR** |
| **1** | ABCA3 | 1.03 | 6.19 | 2.47E-21 | 2.14E-18 |
| **2** | ACHE | 1.43 | 0.98 | 1.18E-06 | 4.80E-05 |
| **3** | ACSM3 | -1.09 | 1.44 | 1.60E-05 | 4.20E-04 |
| **4** | ADAMTS12 | -2.74 | -1.18 | 3.72E-04 | 4.99E-03 |
| **5** | ALG1L11P | 1.51 | -0.14 | 6.93E-04 | 8.19E-03 |
| **6** | AMBN | -1.58 | 1.14 | 1.14E-10 | 1.32E-08 |
| **7** | ANKRD34B | -1.64 | -0.17 | 3.60E-04 | 4.84E-03 |
| **8** | ANXA1 | -2.60 | 2.16 | 1.25E-20 | 8.15E-18 |
| **9** | APOBEC3B | 1.20 | 1.37 | 7.19E-05 | 1.39E-03 |
| **10** | ARC | 2.65 | 2.26 | 2.15E-29 | 4.78E-26 |
| **11** | ARL4C | -1.26 | 3.23 | 2.61E-08 | 1.75E-06 |
| **12** | ASPHD1 | 1.09 | 0.92 | 2.87E-04 | 4.01E-03 |
| **13** | ATP5IF1 | -1.10 | 5.16 | 7.71E-04 | 8.86E-03 |
| **14** | BCL6B | 1.83 | 1.82 | 2.09E-07 | 1.08E-05 |
| **15** | BEGAIN | 1.17 | 1.29 | 6.88E-05 | 1.34E-03 |
| **16** | BIN1 | -1.11 | 2.31 | 2.79E-05 | 6.58E-04 |
| **17** | BTBD19 | -2.33 | -0.27 | 4.45E-07 | 2.11E-05 |
| **18** | C11orf96 | 1.71 | 3.02 | 1.07E-07 | 6.07E-06 |
| **19** | CASP10 | -1.49 | 0.80 | 7.26E-06 | 2.17E-04 |
| **20** | CCDC80 | -1.33 | 1.64 | 3.60E-07 | 1.75E-05 |
| **21** | CCN2 | -1.40 | 4.18 | 1.30E-21 | 1.20E-18 |
| **22** | CD40 | 2.27 | 0.11 | 6.42E-05 | 1.27E-03 |
| **23** | CDC26P1 | -1.20 | 2.20 | 8.45E-04 | 9.49E-03 |
| **24** | CDH12 | 1.52 | 1.95 | 4.58E-12 | 6.61E-10 |
| **25** | CDHR1 | 1.24 | 2.99 | 3.11E-15 | 7.81E-13 |
| **26** | CDKN1A | 1.47 | 4.71 | 1.43E-22 | 1.56E-19 |
| **27** | CFAP298 | -1.12 | 4.52 | 2.47E-04 | 3.59E-03 |
| **28** | CHMP4BP1 | -1.88 | 0.98 | 7.69E-04 | 8.85E-03 |
| **29** | CHRDL1 | -3.07 | -0.68 | 1.56E-12 | 2.45E-10 |
| **30** | CHST8 | 1.49 | 0.54 | 1.53E-04 | 2.47E-03 |
| **31** | COL27A1 | -1.01 | 4.22 | 1.87E-05 | 4.76E-04 |
| **32** | COL5A1 | -1.18 | 4.40 | 1.50E-22 | 1.56E-19 |
| **33** | COLEC12 | -1.14 | 4.01 | 4.11E-15 | 9.71E-13 |
| **34** | CORIN | 1.70 | 0.31 | 1.34E-10 | 1.54E-08 |
| **35** | CPNE7 | -3.02 | 0.44 | 9.08E-15 | 2.08E-12 |
| **36** | CPS1 | -1.05 | 5.72 | 7.87E-20 | 4.25E-17 |
| **37** | CRTAC1 | 1.32 | 0.53 | 1.64E-05 | 4.30E-04 |
| **38** | CSRNP1 | 1.32 | 4.16 | 1.07E-16 | 3.72E-14 |
| **39** | CYB561 | 1.58 | 1.83 | 1.43E-07 | 7.66E-06 |
| **40** | CYBB | 1.32 | 0.07 | 4.57E-04 | 5.85E-03 |
| **41** | CYRIA | -2.57 | -1.49 | 2.44E-04 | 3.55E-03 |
| **42** | DECR1 | -1.42 | 0.92 | 9.17E-07 | 3.85E-05 |
| **43** | DEFB109D | -3.26 | -0.73 | 2.13E-05 | 5.29E-04 |

| | | | | | |
|---|---|---|---|---|---|
| **44** | DHX32 | 1.21 | 4.38 | 2.25E-15 | 5.85E-13 |
| **45** | DLX5 | -1.03 | 1.73 | 2.65E-04 | 3.77E-03 |
| **46** | DNAH17 | 1.90 | 2.75 | 2.24E-19 | 1.13E-16 |
| **47** | DNAH17-AS1 | 2.13 | -0.62 | 7.42E-07 | 3.26E-05 |
| **48** | DTX3 | -1.11 | 0.84 | 1.75E-04 | 2.73E-03 |
| **49** | DUSP5P1 | -5.46 | -0.98 | 1.09E-04 | 1.91E-03 |
| **50** | EEF1A2 | 1.39 | 7.12 | 9.07E-27 | 1.77E-23 |
| **51** | EFEMP1 | -1.82 | 1.74 | 1.35E-14 | 3.02E-12 |
| **52** | EGR1 | 1.06 | 4.38 | 2.06E-05 | 5.16E-04 |
| **53** | EGR3 | 1.52 | 0.59 | 2.03E-05 | 5.09E-04 |
| **54** | EMP3 | 1.18 | 4.62 | 7.40E-13 | 1.23E-10 |
| **55** | EMX2 | -1.14 | 1.57 | 3.40E-04 | 4.61E-03 |
| **56** | EPHB1 | -1.38 | 1.76 | 3.34E-09 | 2.84E-07 |
| **57** | FAM83G | 1.35 | 4.26 | 2.80E-17 | 1.15E-14 |
| **58** | FBXL7 | -2.24 | 0.82 | 1.51E-09 | 1.44E-07 |
| **59** | FCGBP | 1.49 | 0.97 | 1.38E-05 | 3.70E-04 |
| **60** | FGF13 | -1.13 | 1.02 | 3.06E-07 | 1.52E-05 |
| **61** | FHDC1 | 1.62 | 0.48 | 4.92E-05 | 1.03E-03 |
| **62** | FLG | 1.08 | 3.88 | 1.79E-05 | 4.63E-04 |
| **63** | FLRT3 | -1.18 | 0.96 | 7.97E-04 | 9.09E-03 |
| **64** | FN3K | -1.37 | -0.29 | 4.81E-04 | 6.08E-03 |
| **65** | FOSL1 | 1.13 | 3.22 | 1.27E-04 | 2.13E-03 |
| **66** | FOXCUT | 1.19 | 2.18 | 1.79E-04 | 2.79E-03 |
| **67** | FXYD6 | -1.05 | 2.06 | 1.19E-06 | 4.83E-05 |
| **68** | FZD9 | 1.26 | 2.64 | 8.87E-04 | 9.87E-03 |
| **69** | GALNT14 | -1.45 | 0.59 | 1.07E-08 | 8.22E-07 |
| **70** | GHR | -1.25 | 2.62 | 2.19E-11 | 2.90E-09 |
| **71** | GRM8 | -1.78 | 0.01 | 3.21E-04 | 4.42E-03 |
| **72** | GSTO2 | 1.05 | 0.19 | 4.51E-04 | 5.78E-03 |
| **73** | H2BC10 | 2.67 | -0.69 | 7.50E-05 | 1.44E-03 |
| **74** | H3C11 | 1.26 | 3.22 | 1.87E-06 | 7.01E-05 |
| **75** | HAND1 | -1.26 | 3.92 | 1.15E-08 | 8.72E-07 |
| **76** | HARBI1 | 1.37 | 1.04 | 1.97E-04 | 3.01E-03 |
| **77** | HHLA3 | -1.19 | 3.36 | 4.26E-05 | 9.28E-04 |
| **78** | HLA-DOA | -1.83 | 0.71 | 5.44E-09 | 4.44E-07 |
| **79** | HMOX1 | 1.61 | 3.44 | 6.96E-15 | 1.62E-12 |
| **80** | HNRNPH1 | -1.10 | 8.51 | 4.29E-06 | 1.44E-04 |
| **81** | HOXA1 | -1.94 | 1.88 | 1.37E-05 | 3.69E-04 |
| **82** | HOXA10-HOXA9 | -2.06 | 1.49 | 3.78E-14 | 7.85E-12 |
| **83** | HOXA4 | -1.06 | 1.56 | 2.68E-04 | 3.80E-03 |
| **84** | HOXB3 | -1.02 | 2.50 | 3.38E-04 | 4.60E-03 |
| **85** | HOXB6 | -1.29 | 3.45 | 1.85E-15 | 5.07E-13 |
| **86** | HOXB8 | -1.45 | 3.37 | 2.55E-07 | 1.30E-05 |
| **87** | HOXB-AS3 | -1.25 | 0.02 | 7.39E-04 | 8.58E-03 |
| **88** | HSBP1L1 | 1.47 | 1.04 | 2.11E-08 | 1.48E-06 |
| **89** | HSPA6 | 2.67 | -0.87 | 6.25E-06 | 1.93E-04 |

| | | | | | |
|---|---|---|---|---|---|
| **90** | IFI44L | -2.04 | -0.10 | 1.32E-07 | 7.12E-06 |
| **91** | IFITM2 | -1.13 | 5.02 | 5.26E-06 | 1.67E-04 |
| **92** | IFITM3 | -1.14 | 0.96 | 4.40E-06 | 1.47E-04 |
| **93** | IGF2 | 1.56 | 3.99 | 8.80E-10 | 8.85E-08 |
| **94** | IGFBP5 | -1.95 | 3.31 | 9.32E-25 | 1.45E-21 |
| **95** | IL11 | 2.06 | -0.81 | 5.60E-05 | 1.14E-03 |
| **96** | IL12RB2 | 2.60 | 3.53 | 2.47E-17 | 1.04E-14 |
| **97** | IL15RA | 1.29 | 1.41 | 5.33E-04 | 6.57E-03 |
| **98** | IL6R | 1.21 | 3.16 | 1.51E-13 | 2.83E-11 |
| **99** | INSIG1-DT | 1.32 | 0.89 | 4.99E-05 | 1.05E-03 |
| **100** | IRF5 | 1.00 | 2.54 | 9.88E-07 | 4.07E-05 |
| **101** | ITGA7 | 1.05 | 2.51 | 2.43E-04 | 3.54E-03 |
| **102** | JAKMIP2 | -1.30 | 2.68 | 5.31E-06 | 1.68E-04 |
| **103** | KCNS3 | -1.08 | 0.47 | 7.86E-04 | 8.99E-03 |
| **104** | KIFC2 | 1.01 | 1.90 | 3.44E-05 | 7.76E-04 |
| **105** | KIT | -1.28 | 3.88 | 6.19E-21 | 4.60E-18 |
| **106** | KLHL41 | -1.31 | 1.07 | 4.51E-06 | 1.49E-04 |
| **107** | KRT17 | 1.63 | -0.75 | 8.15E-04 | 9.24E-03 |
| **108** | KRT222 | -1.48 | -0.68 | 5.54E-05 | 1.13E-03 |
| **109** | L3MBTL4 | 1.10 | 1.99 | 6.35E-08 | 3.84E-06 |
| **110** | LAMB3 | 1.67 | -0.05 | 1.28E-05 | 3.48E-04 |
| **111** | LAPTM5 | 3.31 | -0.56 | 3.41E-15 | 8.32E-13 |
| **112** | LGALS3 | -1.82 | 0.24 | 2.97E-06 | 1.05E-04 |
| **113** | LINC01234 | 1.74 | 0.52 | 3.10E-04 | 4.29E-03 |
| **114** | LINC03011 | 1.48 | 0.13 | 1.60E-06 | 6.22E-05 |
| **115** | LINGO2 | 1.39 | 3.17 | 1.09E-14 | 2.47E-12 |
| **116** | LOC100130992 | 1.82 | -0.25 | 1.85E-04 | 2.86E-03 |
| **117** | LOC100507412 | -1.21 | 2.03 | 6.37E-06 | 1.96E-04 |
| **118** | LOC101928126 | -2.03 | -0.42 | 3.46E-04 | 4.68E-03 |
| **119** | LOC124906720 | 3.95 | -1.46 | 1.17E-04 | 2.02E-03 |
| **120** | LOC642969 | 1.87 | 1.46 | 4.43E-04 | 5.72E-03 |
| **121** | LOC644462 | 1.09 | 0.71 | 3.89E-04 | 5.18E-03 |
| **122** | LRRCC1 | -1.00 | 2.74 | 1.81E-05 | 4.64E-04 |
| **123** | LRRTM4 | -4.32 | -1.30 | 9.40E-09 | 7.36E-07 |
| **124** | LTBP4 | 1.02 | 5.53 | 2.91E-08 | 1.92E-06 |
| **125** | MACROD2 | 1.10 | 1.54 | 2.17E-06 | 8.00E-05 |
| **126** | MAFA | 1.47 | 1.04 | 1.18E-04 | 2.02E-03 |
| **127** | MAGEB2 | 2.98 | 4.31 | 3.61E-87 | 2.82E-83 |
| **128** | MAL2 | -2.50 | 1.18 | 3.13E-19 | 1.53E-16 |
| **129** | MAMSTR | 2.03 | -0.06 | 3.75E-04 | 5.01E-03 |
| **130** | MAOB | -1.79 | 0.17 | 3.72E-06 | 1.28E-04 |
| **131** | MAP2 | -1.16 | 1.39 | 7.57E-08 | 4.45E-06 |
| **132** | MFAP2 | -4.66 | 2.08 | 2.38E-65 | 9.26E-62 |
| **133** | MICA-AS1 | 2.88 | -0.30 | 6.91E-06 | 2.08E-04 |
| **134** | MSL3P1 | -1.35 | 2.14 | 2.02E-09 | 1.83E-07 |
| **135** | MSS51 | 1.51 | 0.29 | 7.00E-04 | 8.25E-03 |

| | | | | | |
|---|---|---|---|---|---|
| **136** | MT2A | -1.25 | 3.96 | 4.31E-07 | 2.05E-05 |
| **137** | MXRA8 | -1.32 | 0.69 | 4.17E-04 | 5.48E-03 |
| **138** | MYO1G | -1.02 | 0.03 | 3.64E-04 | 4.89E-03 |
| **139** | NCL | -1.27 | 9.48 | 9.53E-06 | 2.74E-04 |
| **140** | NEFM | 1.57 | 9.65 | 4.18E-13 | 7.24E-11 |
| **141** | NMNAT2 | 1.57 | 0.48 | 3.07E-06 | 1.08E-04 |
| **142** | NOS2 | 1.17 | 1.78 | 5.08E-06 | 1.63E-04 |
| **143** | NPR1 | -1.66 | -0.32 | 5.17E-04 | 6.42E-03 |
| **144** | NR2F1 | -1.02 | 4.66 | 8.98E-18 | 3.89E-15 |
| **145** | NUTM2A | 1.22 | 2.04 | 4.24E-05 | 9.25E-04 |
| **146** | OAS3 | -1.29 | 2.09 | 1.81E-11 | 2.44E-09 |
| **147** | OLMALINC | -1.25 | 2.08 | 1.63E-06 | 6.28E-05 |
| **148** | OSTN | -1.65 | 2.26 | 4.48E-13 | 7.60E-11 |
| **149** | OXCT1 | -1.76 | 2.96 | 2.07E-25 | 3.59E-22 |
| **150** | P2RX5 | 1.57 | 2.06 | 7.27E-14 | 1.47E-11 |
| **151** | P4HTM | -1.34 | 1.63 | 4.16E-05 | 9.15E-04 |
| **152** | PAK3 | -1.40 | 0.37 | 1.02E-04 | 1.81E-03 |
| **153** | PAPPA | -3.14 | 0.54 | 2.04E-09 | 1.84E-07 |
| **154** | PAX1 | -6.43 | -1.60 | 6.78E-06 | 2.06E-04 |
| **155** | PCDH18 | -1.04 | 2.67 | 1.02E-05 | 2.89E-04 |
| **156** | PCYT1B | 1.22 | 4.47 | 8.32E-22 | 8.11E-19 |
| **157** | PDE3A | -3.30 | -0.99 | 3.00E-06 | 1.06E-04 |
| **158** | PDE4B | -1.43 | -0.19 | 9.14E-05 | 1.67E-03 |
| **159** | PGAM4 | 1.60 | 0.76 | 8.84E-05 | 1.64E-03 |
| **160** | PGR | 1.32 | 4.52 | 3.42E-21 | 2.80E-18 |
| **161** | PHEX | 1.11 | 0.95 | 6.78E-04 | 8.05E-03 |
| **162** | PITPNM3 | 1.14 | 0.94 | 8.33E-05 | 1.56E-03 |
| **163** | PLCL1 | 3.50 | 3.98 | 7.95E-154 | 1.24E-149 |
| **164** | PLPP4 | -1.15 | 0.60 | 3.99E-04 | 5.30E-03 |
| **165** | PLXDC2 | -2.17 | -0.12 | 1.38E-06 | 5.43E-05 |
| **166** | PPP2R2C | 1.17 | 3.06 | 3.10E-13 | 5.55E-11 |
| **167** | PTMAP9 | -1.62 | 5.04 | 3.60E-04 | 4.84E-03 |
| **168** | PTN | -4.68 | -1.41 | 1.59E-06 | 6.17E-05 |
| **169** | PYCARD-AS1 | -1.31 | 1.59 | 2.02E-05 | 5.08E-04 |
| **170** | RAB30-DT | 1.17 | 1.59 | 1.83E-06 | 6.91E-05 |
| **171** | RAB35-AS1 | -1.54 | 1.27 | 7.12E-05 | 1.39E-03 |
| **172** | RASA4B | 1.14 | 2.82 | 1.64E-11 | 2.23E-09 |
| **173** | RASD2 | 1.86 | -0.15 | 2.72E-04 | 3.85E-03 |
| **174** | RASL11A | -1.00 | 3.16 | 5.56E-12 | 7.88E-10 |
| **175** | RELN | -2.61 | 0.31 | 2.17E-14 | 4.63E-12 |
| **176** | RELT | 1.09 | 3.96 | 1.06E-09 | 1.06E-07 |
| **177** | RGS2 | -1.07 | 3.60 | 8.50E-12 | 1.19E-09 |
| **178** | ROR2 | -1.39 | 0.64 | 2.37E-04 | 3.47E-03 |
| **179** | RPL21P75 | -1.02 | 2.59 | 1.56E-05 | 4.10E-04 |
| **180** | RPL22L1 | -1.01 | 5.71 | 2.01E-09 | 1.83E-07 |
| **181** | RPL36AP37 | -3.14 | 5.60 | 8.19E-04 | 9.26E-03 |

| | | | | | |
|---|---|---|---|---|---|
| **182** | RPRM | -1.20 | 2.46 | 7.20E-09 | 5.75E-07 |
| **183** | SBF2-AS1 | 1.12 | 1.62 | 2.54E-06 | 9.26E-05 |
| **184** | SCARA5 | 1.02 | 0.72 | 1.22E-04 | 2.06E-03 |
| **185** | SCIN | -1.45 | 0.17 | 6.62E-06 | 2.02E-04 |
| **186** | SERPINB8 | 1.74 | 0.30 | 2.64E-07 | 1.33E-05 |
| **187** | SERPINF1 | -1.65 | 1.52 | 1.72E-16 | 5.60E-14 |
| **188** | SERTAD1 | 1.05 | 1.76 | 2.50E-04 | 3.62E-03 |
| **189** | SFRP1 | -2.82 | 1.93 | 5.39E-31 | 1.40E-27 |
| **190** | SFRP2 | 2.39 | 2.69 | 6.82E-21 | 4.83E-18 |
| **191** | SH2B2 | 1.15 | 2.42 | 1.28E-07 | 7.03E-06 |
| **192** | SLC10A3 | 1.03 | 3.18 | 9.04E-07 | 3.82E-05 |
| **193** | SLC13A3 | 1.07 | 1.17 | 7.42E-05 | 1.42E-03 |
| **194** | SLFN12 | -3.13 | 0.33 | 1.67E-20 | 1.04E-17 |
| **195** | SMAD6 | -1.65 | 2.97 | 1.80E-19 | 9.35E-17 |
| **196** | SMARCA1 | -1.79 | 4.32 | 8.24E-39 | 2.57E-35 |
| **197** | SMIM10 | -1.33 | 1.41 | 4.91E-05 | 1.03E-03 |
| **198** | SNAI2 | -1.80 | 4.06 | 3.76E-21 | 2.93E-18 |
| **199** | SNCB | -1.13 | 0.90 | 2.51E-04 | 3.64E-03 |
| **200** | SOX11 | -3.59 | 0.16 | 4.25E-13 | 7.27E-11 |
| **201** | SP6 | 1.94 | 0.02 | 4.33E-05 | 9.35E-04 |
| **202** | SPARC | -4.64 | 2.59 | 1.65E-70 | 8.56E-67 |
| **203** | SPOCK1 | -2.04 | 0.07 | 1.12E-10 | 1.31E-08 |
| **204** | STING1 | -1.33 | 2.29 | 2.58E-10 | 2.83E-08 |
| **205** | SULT1C4 | -4.44 | 0.17 | 1.26E-15 | 3.52E-13 |
| **206** | SYN3 | 1.18 | 2.23 | 4.18E-09 | 3.54E-07 |
| **207** | SYT14 | 1.13 | 2.59 | 1.24E-06 | 5.00E-05 |
| **208** | SYTL5 | 1.10 | 2.77 | 1.42E-11 | 1.98E-09 |
| **209** | TAC1 | 1.40 | 1.75 | 4.89E-09 | 4.05E-07 |
| **210** | TAGLN | -1.18 | 0.57 | 8.42E-05 | 1.57E-03 |
| **211** | TEPSIN | 1.02 | 3.38 | 1.40E-05 | 3.75E-04 |
| **212** | THNSL2 | 1.84 | -0.46 | 9.28E-05 | 1.69E-03 |
| **213** | TIGD1 | -1.08 | 3.46 | 1.98E-05 | 5.02E-04 |
| **214** | TLCD3B | 1.12 | 0.63 | 1.60E-04 | 2.55E-03 |
| **215** | TMEM132E | 1.41 | 0.97 | 7.38E-05 | 1.42E-03 |
| **216** | TMEM200B | 2.49 | -0.57 | 3.46E-05 | 7.79E-04 |
| **217** | TMT1A | -1.20 | 3.73 | 7.78E-18 | 3.47E-15 |
| **218** | TNFRSF12A | 1.53 | 3.47 | 5.92E-11 | 7.21E-09 |
| **219** | TNFSF9 | 2.14 | 0.87 | 3.14E-06 | 1.10E-04 |
| **220** | TOMM40P2 | 2.24 | 1.61 | 4.62E-04 | 5.89E-03 |
| **221** | TUBA4A | -1.60 | 0.57 | 4.69E-04 | 5.96E-03 |
| **222** | UBE2QL1 | -1.18 | 0.63 | 1.12E-04 | 1.95E-03 |
| **223** | UGT3A2 | -1.33 | 1.85 | 1.12E-08 | 8.55E-07 |
| **224** | USP32P2 | -1.92 | 0.33 | 5.31E-06 | 1.68E-04 |
| **225** | VGF | 2.10 | 1.33 | 1.88E-20 | 1.12E-17 |
| **226** | WSCD1 | 1.20 | 4.67 | 1.54E-14 | 3.32E-12 |
| **227** | XK | 1.14 | 3.75 | 7.32E-16 | 2.11E-13 |

| | | | | | |
|---|---|---|---|---|---|
| **228** | ZDHHC11 | -1.25 | 1.58 | 3.12E-05 | 7.17E-04 |
| **229** | ZNF22 | -3.39 | 1.08 | 7.79E-24 | 1.01E-20 |
| **230** | ZNF365 | 1.50 | 2.20 | 2.50E-09 | 2.20E-07 |
| **231** | ZNF43 | -1.42 | 0.58 | 6.04E-07 | 2.75E-05 |
| **232** | ZNF830 | -1.03 | 2.58 | 8.53E-04 | 9.57E-03 |
| **233** | ZNF883 | -1.73 | 3.05 | 3.29E-15 | 8.15E-13 |
| **234** | ZSCAN16-AS1 | -1.18 | 2.19 | 5.92E-06 | 1.85E-04 |

**E475G homozygous cell line *vs.* wt**

| | **Gene** | **logFC** | **logCPM** | **PValue** | **FDR** |
|---|---|---|---|---|---|
| **1** | A1BG-AS1 | 1,23 | 0,91 | 4,41E-04 | 3,49E-03 |
| **2** | AARSD1 | -1,19 | 2,87 | 3,78E-04 | 3,07E-03 |
| **3** | ABCA3 | 1,05 | 6,19 | 3,06E-22 | 7,56E-20 |
| **4** | ABCC2 | 1,39 | 1,07 | 7,61E-05 | 8,16E-04 |
| **5** | ABLIM3 | -1,14 | 0,08 | 2,87E-04 | 2,45E-03 |
| **6** | ACHE | 1,15 | 0,98 | 1,41E-04 | 1,36E-03 |
| **7** | ADAM12 | -4,28 | -0,42 | 1,34E-10 | 6,87E-09 |
| **8** | ADAM8 | 2,03 | -0,68 | 1,70E-04 | 1,60E-03 |
| **9** | ADAMTS12 | -3,03 | -1,18 | 4,49E-05 | 5,25E-04 |
| **10** | ADAMTS16 | -3,84 | 1,48 | 1,68E-39 | 1,64E-36 |
| **11** | ADCYAP1R1 | -1,05 | 2,03 | 1,24E-06 | 2,44E-05 |
| **12** | ADSS1 | -1,27 | -0,40 | 3,67E-04 | 2,99E-03 |
| **13** | AFF2 | -1,05 | 4,91 | 2,35E-12 | 1,61E-10 |
| **14** | AGAP2 | 1,44 | -0,06 | 7,46E-05 | 8,03E-04 |
| **15** | ALDH3B1 | 1,47 | -0,40 | 3,82E-04 | 3,10E-03 |
| **16** | ALG1L13P | 1,07 | 2,44 | 1,11E-04 | 1,11E-03 |
| **17** | AMBN | -2,91 | 1,14 | 6,08E-26 | 2,43E-23 |
| **18** | ANKRD1 | -1,84 | -0,35 | 6,86E-05 | 7,46E-04 |
| **19** | ANKRD11P2 | 1,59 | 1,46 | 1,03E-04 | 1,05E-03 |
| **20** | ANO7 | -1,51 | 0,99 | 1,94E-05 | 2,57E-04 |
| **21** | ANXA1 | -2,07 | 2,16 | 5,42E-15 | 5,82E-13 |
| **22** | APCDD1L | 1,28 | 0,39 | 3,08E-04 | 2,60E-03 |
| **23** | APOBEC3B | 1,74 | 1,37 | 6,77E-09 | 2,49E-07 |
| **24** | ARAP1-AS2 | 2,43 | -0,39 | 3,88E-04 | 3,14E-03 |
| **25** | ARC | 1,58 | 2,26 | 8,31E-11 | 4,41E-09 |
| **26** | ARHGAP22 | 1,03 | 3,78 | 3,07E-06 | 5,28E-05 |
| **27** | ARHGAP27P1 | 1,78 | 0,60 | 4,10E-05 | 4,87E-04 |
| **28** | ARL4C | -1,53 | 3,23 | 4,32E-11 | 2,39E-09 |
| **29** | ARRDC2 | 1,24 | 2,23 | 3,25E-04 | 2,73E-03 |
| **30** | ARX | 2,31 | -0,81 | 3,30E-05 | 4,07E-04 |
| **31** | ASPHD1 | 1,02 | 0,92 | 6,16E-04 | 4,60E-03 |
| **32** | ATP2B3 | -1,70 | 2,09 | 4,02E-12 | 2,63E-10 |
| **33** | ATP5IF1 | -1,10 | 5,16 | 7,63E-04 | 5,49E-03 |
| **34** | ATP5PF | -1,18 | 5,63 | 4,27E-09 | 1,66E-07 |
| **35** | ATP8A2 | -1,09 | 2,16 | 2,10E-07 | 5,16E-06 |
| **36** | B3GALT5 | -2,56 | -0,77 | 1,16E-04 | 1,15E-03 |

| | | | | | |
|---|---|---|---|---|---|
| **37** | B4GALNT1 | 1,02 | 4,05 | 1,15E-16 | 1,61E-14 |
| **38** | BCHE | -1,16 | 2,95 | 7,75E-11 | 4,15E-09 |
| **39** | BCL6B | 2,76 | 1,82 | 2,19E-18 | 4,06E-16 |
| **40** | BCYRN1 | -3,57 | 2,76 | 7,86E-04 | 5,62E-03 |
| **41** | BEND3P3 | 1,06 | 4,11 | 4,61E-05 | 5,36E-04 |
| **42** | BHLHA15 | 1,84 | 1,24 | 7,67E-07 | 1,61E-05 |
| **43** | BIN1 | -1,11 | 2,31 | 2,64E-05 | 3,37E-04 |
| **44** | BLOC1S5-TXNDC5 | -9,32 | -0,26 | 1,16E-03 | 7,70E-03 |
| **45** | BMP2 | -2,86 | 0,39 | 1,47E-08 | 5,03E-07 |
| **46** | BRSK1 | 1,34 | 2,62 | 7,96E-07 | 1,66E-05 |
| **47** | BRSK2 | 1,52 | 4,31 | 9,85E-26 | 3,84E-23 |
| **48** | C1QL1 | -1,10 | 2,38 | 1,28E-06 | 2,52E-05 |
| **49** | C4orf19 | 1,75 | 0,31 | 1,77E-07 | 4,43E-06 |
| **50** | CA2 | -1,49 | 6,37 | 2,93E-17 | 4,74E-15 |
| **51** | CAMK1D | 1,08 | 4,35 | 1,10E-28 | 5,36E-26 |
| **52** | CASKIN1 | 1,14 | 3,42 | 3,62E-11 | 2,04E-09 |
| **53** | CBFA2T3 | -1,83 | 0,29 | 2,16E-05 | 2,84E-04 |
| **54** | CCDC144BP | -2,00 | 1,67 | 1,61E-05 | 2,19E-04 |
| **55** | CCN1 | -1,23 | 7,98 | 2,85E-37 | 2,11E-34 |
| **56** | CCN2 | -2,88 | 4,18 | 8,25E-74 | 2,14E-70 |
| **57** | CCN3 | 1,68 | 2,12 | 1,62E-16 | 2,20E-14 |
| **58** | CCR6 | 1,68 | -0,21 | 1,05E-04 | 1,06E-03 |
| **59** | CCZ1P1 | 1,38 | 2,32 | 1,05E-06 | 2,12E-05 |
| **60** | CD40 | 2,18 | 0,11 | 1,32E-04 | 1,28E-03 |
| **61** | CDC26 | -1,15 | 3,61 | 2,32E-06 | 4,20E-05 |
| **62** | CDH12 | 1,95 | 1,95 | 2,23E-19 | 4,45E-17 |
| **63** | CDHR1 | 1,53 | 2,99 | 6,07E-23 | 1,58E-20 |
| **64** | CEBPA-DT | 1,97 | 0,22 | 1,13E-03 | 7,54E-03 |
| **65** | CENPBD1P | 1,24 | 1,89 | 7,43E-07 | 1,57E-05 |
| **66** | CEP63 | -1,04 | 3,01 | 7,13E-04 | 5,20E-03 |
| **67** | CFAP298 | -1,03 | 4,52 | 7,49E-04 | 5,41E-03 |
| **68** | CFDP1 | -1,44 | 3,87 | 2,83E-04 | 2,43E-03 |
| **69** | CHMP4A | -1,11 | 2,38 | 1,58E-03 | 9,87E-03 |
| **70** | CHP1P2 | 1,08 | 1,84 | 4,94E-04 | 3,84E-03 |
| **71** | CHRDL1 | -8,91 | -0,68 | 2,48E-24 | 7,74E-22 |
| **72** | CHST8 | 1,37 | 0,54 | 5,78E-04 | 4,37E-03 |
| **73** | CIITA | 1,04 | 1,65 | 4,32E-05 | 5,09E-04 |
| **74** | CLCN4 | -1,13 | 3,57 | 6,15E-11 | 3,34E-09 |
| **75** | CLSTN2 | -1,99 | 1,47 | 2,63E-13 | 2,17E-11 |
| **76** | COL12A1 | -2,72 | 2,36 | 2,26E-23 | 6,07E-21 |
| **77** | COL26A1 | 1,34 | 2,16 | 3,05E-11 | 1,76E-09 |
| **78** | COL5A1 | -1,77 | 4,40 | 1,02E-46 | 1,32E-43 |
| **79** | COL7A1 | 1,02 | 0,47 | 1,47E-03 | 9,34E-03 |
| **80** | COL9A3 | -1,24 | 0,08 | 1,11E-03 | 7,44E-03 |
| **81** | CORIN | 1,70 | 0,31 | 1,60E-10 | 8,02E-09 |
| **82** | CPNE7 | -3,31 | 0,44 | 7,55E-17 | 1,11E-14 |

| | | | | | |
|---|---|---|---|---|---|
| **83** | CRIP2 | -1,31 | 0,68 | 1,72E-05 | 2,32E-04 |
| **84** | CRLF1 | -1,19 | 0,02 | 1,41E-03 | 9,00E-03 |
| **85** | CRYGD | -1,08 | 2,22 | 3,34E-05 | 4,11E-04 |
| **86** | CSPG4P11 | 1,08 | 2,75 | 5,47E-05 | 6,14E-04 |
| **87** | CTH | 1,00 | 3,47 | 4,90E-10 | 2,29E-08 |
| **88** | CYB561 | 1,62 | 1,83 | 6,97E-08 | 1,97E-06 |
| **89** | CYBB | 1,23 | 0,07 | 9,19E-04 | 6,37E-03 |
| **90** | CYRIA | -3,74 | -1,49 | 1,34E-06 | 2,61E-05 |
| **91** | DECR1 | -2,33 | 0,92 | 2,49E-14 | 2,39E-12 |
| **92** | DGAT2 | 1,05 | 1,05 | 8,80E-05 | 9,18E-04 |
| **93** | DHX32 | 1,23 | 4,38 | 4,03E-16 | 5,19E-14 |
| **94** | DKK1 | 2,01 | -0,33 | 2,83E-04 | 2,43E-03 |
| **95** | DMRT2 | -6,03 | -1,06 | 7,71E-06 | 1,15E-04 |
| **96** | DMRT3 | -2,46 | 0,29 | 2,50E-04 | 2,20E-03 |
| **97** | DNAH17 | 1,53 | 2,75 | 5,12E-13 | 3,95E-11 |
| **98** | DNAH17-AS1 | 1,46 | -0,62 | 9,66E-04 | 6,65E-03 |
| **99** | DNAJC30 | 1,14 | 3,44 | 9,77E-12 | 6,07E-10 |
| **100** | DPP7 | -1,24 | 0,79 | 3,03E-06 | 5,22E-05 |
| **101** | DTNB-AS1 | 3,70 | -0,29 | 9,37E-05 | 9,68E-04 |
| **102** | DTX3 | -2,73 | 0,84 | 5,89E-17 | 8,83E-15 |
| **103** | DUSP5P1 | -5,45 | -0,98 | 1,09E-04 | 1,09E-03 |
| **104** | DUSP6 | 3,47 | 0,29 | 7,20E-04 | 5,24E-03 |
| **105** | ECEL1 | -1,28 | 2,38 | 2,50E-06 | 4,47E-05 |
| **106** | EDIL3 | -1,37 | 3,16 | 9,85E-17 | 1,40E-14 |
| **107** | EEF1A1P50 | 2,37 | -1,04 | 1,54E-03 | 9,71E-03 |
| **108** | EEF1A2 | 1,70 | 7,12 | 1,12E-38 | 9,22E-36 |
| **109** | EFCAB15P | 2,40 | -0,94 | 5,86E-04 | 4,42E-03 |
| **110** | EFEMP1 | -1,94 | 1,74 | 3,77E-16 | 4,89E-14 |
| **111** | EGLN1P1 | 1,63 | 0,18 | 3,38E-04 | 2,81E-03 |
| **112** | EHD2 | -1,81 | -0,22 | 9,73E-05 | 1,00E-03 |
| **113** | ELL3 | 1,22 | 1,33 | 2,46E-05 | 3,16E-04 |
| **114** | EMILIN2 | -1,22 | 3,84 | 2,52E-26 | 1,06E-23 |
| **115** | EML2-AS1 | 1,75 | -0,59 | 1,56E-04 | 1,48E-03 |
| **116** | EML3 | 1,05 | 4,28 | 3,78E-06 | 6,30E-05 |
| **117** | EMP3 | 1,25 | 4,62 | 2,65E-14 | 2,54E-12 |
| **118** | EMX1 | 1,14 | 2,11 | 2,58E-04 | 2,25E-03 |
| **119** | ENO2 | 1,02 | 4,68 | 1,48E-10 | 7,50E-09 |
| **120** | ENTREP2 | 1,07 | 2,99 | 1,67E-10 | 8,23E-09 |
| **121** | EPHB1 | -2,30 | 1,76 | 6,84E-20 | 1,46E-17 |
| **122** | ERBB3 | 1,28 | 1,11 | 2,30E-08 | 7,45E-07 |
| **123** | ETS1 | -1,10 | 1,63 | 9,86E-07 | 2,01E-05 |
| **124** | ETV4 | 1,38 | 3,51 | 6,06E-15 | 6,39E-13 |
| **125** | FAAH2 | -2,06 | 0,52 | 1,90E-10 | 9,27E-09 |
| **126** | FAM131C | 1,18 | 1,91 | 6,61E-05 | 7,21E-04 |
| **127** | FAM13C | 1,26 | 1,61 | 1,86E-06 | 3,51E-05 |
| **128** | FAM167A | 1,48 | 0,52 | 3,91E-06 | 6,46E-05 |

| | | | | | |
|---|---|---|---|---|---|
| **129** | FAM83G | 1,09 | 4,26 | 1,20E-11 | 7,31E-10 |
| **130** | FBXL7 | -4,26 | 0,82 | 5,68E-22 | 1,38E-19 |
| **131** | FCGBP | 1,31 | 0,97 | 1,20E-04 | 1,19E-03 |
| **132** | FCHO1 | 1,60 | 2,18 | 3,70E-13 | 2,96E-11 |
| **133** | FGF12 | 1,08 | 2,47 | 4,79E-09 | 1,84E-07 |
| **134** | FGF13 | -3,98 | 1,02 | 7,69E-38 | 5,99E-35 |
| **135** | FGFBP3 | 1,11 | 2,40 | 6,73E-12 | 4,33E-10 |
| **136** | FHIP2A | 1,04 | 5,05 | 5,08E-16 | 6,39E-14 |
| **137** | FKBP2 | -2,27 | 1,90 | 6,26E-04 | 4,66E-03 |
| **138** | FLI1 | -2,13 | 1,54 | 9,91E-14 | 8,87E-12 |
| **139** | FLNB-AS1 | 2,16 | -0,51 | 3,62E-04 | 2,97E-03 |
| **140** | FN1 | -1,49 | 4,48 | 2,39E-25 | 8,48E-23 |
| **141** | FNDC1 | -1,20 | 0,56 | 2,20E-04 | 1,99E-03 |
| **142** | FOSB | 1,01 | 3,28 | 1,24E-11 | 7,52E-10 |
| **143** | FOSL2 | -1,70 | 4,59 | 1,01E-60 | 1,57E-57 |
| **144** | FOXF1 | -1,14 | 2,05 | 3,24E-06 | 5,51E-05 |
| **145** | FOXN4 | 2,02 | 1,00 | 4,23E-17 | 6,59E-15 |
| **146** | FOXO6 | 1,02 | 2,34 | 4,13E-04 | 3,32E-03 |
| **147** | FOXP2 | 1,17 | 4,50 | 1,17E-23 | 3,27E-21 |
| **148** | FRAT1 | 1,29 | 1,28 | 2,32E-04 | 2,08E-03 |
| **149** | FRMPD3 | 1,18 | 1,52 | 6,55E-08 | 1,86E-06 |
| **150** | FXYD6 | -1,68 | 2,06 | 7,62E-14 | 6,99E-12 |
| **151** | FZD9 | 2,19 | 2,64 | 1,64E-10 | 8,16E-09 |
| **152** | GABRE | -3,07 | -0,08 | 1,34E-12 | 9,66E-11 |
| **153** | GABRQ | -3,62 | -0,11 | 7,89E-10 | 3,50E-08 |
| **154** | GALNT14 | -3,04 | 0,59 | 1,26E-24 | 4,19E-22 |
| **155** | GALNT3 | 1,17 | 3,23 | 2,33E-12 | 1,61E-10 |
| **156** | GAS2L1 | 1,18 | 0,92 | 1,69E-04 | 1,59E-03 |
| **157** | GASAL1 | 1,54 | 0,29 | 8,47E-05 | 8,89E-04 |
| **158** | GCM1 | 1,04 | 0,19 | 8,94E-04 | 6,22E-03 |
| **159** | GDPD3 | 1,59 | -0,46 | 4,95E-04 | 3,84E-03 |
| **160** | GFI1 | 1,04 | 2,30 | 2,37E-07 | 5,73E-06 |
| **161** | GFRA1 | 1,27 | 4,17 | 1,13E-30 | 6,08E-28 |
| **162** | GHR | -1,04 | 2,62 | 1,12E-08 | 3,92E-07 |
| **163** | GJB2 | 1,21 | 0,92 | 1,09E-03 | 7,34E-03 |
| **164** | GOLGA2P8 | 1,44 | -0,42 | 1,19E-03 | 7,86E-03 |
| **165** | GPER1 | 1,66 | 0,22 | 2,67E-04 | 2,32E-03 |
| **166** | GPR37 | 1,42 | 2,47 | 1,55E-09 | 6,53E-08 |
| **167** | GPR50 | -1,58 | 1,39 | 8,23E-05 | 8,67E-04 |
| **168** | GRIN2D | 1,17 | 2,21 | 2,56E-08 | 8,18E-07 |
| **169** | GRIP2 | 2,17 | -0,22 | 1,24E-05 | 1,75E-04 |
| **170** | GRM8 | -3,00 | 0,01 | 3,43E-08 | 1,05E-06 |
| **171** | GSPT2 | -1,02 | 3,68 | 1,19E-13 | 1,05E-11 |
| **172** | GUCY1B2 | 1,35 | -0,45 | 1,41E-03 | 9,00E-03 |
| **173** | H1-1 | 1,01 | 4,09 | 6,64E-08 | 1,88E-06 |
| **174** | H19 | -1,60 | -0,17 | 4,20E-05 | 4,96E-04 |

| | | | | | |
|---|---|---|---|---|---|
| **175** | H2AC8 | 1,06 | 5,19 | 2,70E-09 | 1,10E-07 |
| **176** | H2BC10 | 2,22 | -0,69 | 1,49E-03 | 9,45E-03 |
| **177** | H3C11 | 1,85 | 3,22 | 3,75E-12 | 2,50E-10 |
| **178** | HAGHL | 1,15 | 2,10 | 3,62E-05 | 4,38E-04 |
| **179** | HAND1 | -1,09 | 3,92 | 6,23E-07 | 1,35E-05 |
| **180** | HBQ1 | 1,26 | 0,69 | 2,03E-04 | 1,86E-03 |
| **181** | HES7 | 2,79 | -1,11 | 5,33E-06 | 8,40E-05 |
| **182** | HHLA3 | -1,06 | 3,36 | 2,48E-04 | 2,19E-03 |
| **183** | HLA-DOA | -1,98 | 0,71 | 2,38E-10 | 1,16E-08 |
| **184** | HMGA2 | -1,31 | 6,32 | 5,12E-21 | 1,16E-18 |
| **185** | HNF4G | 1,11 | 2,35 | 9,86E-06 | 1,42E-04 |
| **186** | HNRNPH1 | -1,61 | 8,51 | 3,69E-11 | 2,07E-09 |
| **187** | HNRNPH2 | -1,13 | 4,16 | 3,81E-04 | 3,09E-03 |
| **188** | HOOK1 | 1,26 | 4,17 | 5,59E-17 | 8,46E-15 |
| **189** | HOXB8 | -1,15 | 3,37 | 3,31E-05 | 4,07E-04 |
| **190** | HSBP1L1 | 1,94 | 1,04 | 3,41E-14 | 3,22E-12 |
| **191** | HSF2BP | -1,81 | -0,47 | 1,10E-04 | 1,11E-03 |
| **192** | HTRA1 | -1,00 | 2,34 | 1,42E-07 | 3,68E-06 |
| **193** | ICAM5 | 1,57 | 2,38 | 3,68E-09 | 1,45E-07 |
| **194** | IER3 | 1,07 | 1,90 | 3,84E-04 | 3,11E-03 |
| **195** | IFI30 | 1,26 | 3,70 | 4,34E-19 | 8,36E-17 |
| **196** | IFIT3 | 3,62 | -0,72 | 1,02E-03 | 6,96E-03 |
| **197** | IFITM2 | -1,01 | 5,02 | 4,06E-05 | 4,82E-04 |
| **198** | IFITM3 | -1,07 | 0,96 | 1,83E-05 | 2,45E-04 |
| **199** | IGF2 | -2,27 | 3,99 | 8,46E-17 | 1,23E-14 |
| **200** | IGFBP3 | -2,35 | -0,99 | 8,63E-06 | 1,26E-04 |
| **201** | IGFBP5 | -1,57 | 3,31 | 2,86E-17 | 4,69E-15 |
| **202** | IGFBP7 | -1,34 | 1,56 | 8,29E-08 | 2,28E-06 |
| **203** | IKBKGP1 | 1,02 | 2,43 | 1,73E-04 | 1,63E-03 |
| **204** | IKZF3 | 1,62 | 3,91 | 3,03E-32 | 1,89E-29 |
| **205** | IL12RB2 | 2,44 | 3,53 | 1,28E-15 | 1,55E-13 |
| **206** | IL15RA | 1,59 | 1,41 | 1,69E-05 | 2,28E-04 |
| **207** | INSIG1-DT | 1,32 | 0,89 | 4,03E-05 | 4,80E-04 |
| **208** | INSM1 | 1,90 | 0,34 | 4,06E-06 | 6,68E-05 |
| **209** | IRF5 | 1,12 | 2,54 | 4,45E-08 | 1,33E-06 |
| **210** | ITGA7 | 1,05 | 2,51 | 2,66E-04 | 2,31E-03 |
| **211** | ITM2C | -1,06 | 6,48 | 1,66E-12 | 1,17E-10 |
| **212** | ITPR3 | 1,29 | 4,68 | 1,66E-25 | 6,12E-23 |
| **213** | JAKMIP2 | -1,56 | 2,68 | 4,36E-08 | 1,30E-06 |
| **214** | KCNK5 | 1,23 | 2,78 | 8,90E-07 | 1,84E-05 |
| **215** | KCNQ4 | 1,18 | 2,31 | 2,24E-06 | 4,09E-05 |
| **216** | KCNS3 | -1,25 | 0,47 | 2,12E-04 | 1,94E-03 |
| **217** | KIAA0513 | 1,41 | 3,09 | 1,32E-11 | 7,95E-10 |
| **218** | KRT222 | -2,22 | -0,68 | 8,67E-08 | 2,37E-06 |
| **219** | L3MBTL4 | 1,66 | 1,99 | 3,81E-17 | 5,99E-15 |
| **220** | LENG9 | 1,29 | 2,92 | 1,51E-10 | 7,61E-09 |

| | | | | | |
|---|---|---|---|---|---|
| **221** | LGALS3 | -2,12 | 0,24 | 2,69E-07 | 6,42E-06 |
| **222** | LIAT1 | -1,89 | -0,16 | 7,40E-04 | 5,36E-03 |
| **223** | LIME1 | 1,16 | 1,08 | 7,07E-05 | 7,65E-04 |
| **224** | LINC00632 | -2,00 | 6,06 | 3,80E-04 | 3,08E-03 |
| **225** | LINC01311 | 1,03 | 1,76 | 1,50E-04 | 1,44E-03 |
| **226** | LINC01551 | 1,02 | 0,54 | 1,22E-03 | 8,06E-03 |
| **227** | LINC02983 | 1,17 | 0,33 | 8,85E-04 | 6,17E-03 |
| **228** | LINC02985 | 1,14 | 0,51 | 3,27E-04 | 2,74E-03 |
| **229** | LINC03011 | 1,44 | 0,13 | 3,91E-06 | 6,46E-05 |
| **230** | LINC-PINT | 1,15 | 1,45 | 5,47E-04 | 4,18E-03 |
| **231** | LINGO2 | 1,82 | 3,17 | 6,66E-24 | 2,00E-21 |
| **232** | LMOD1 | -1,94 | -0,31 | 1,04E-03 | 7,04E-03 |
| **233** | LOC100101148 | -6,69 | -1,26 | 2,98E-06 | 5,16E-05 |
| **234** | LOC100129931 | 2,83 | -0,42 | 9,83E-05 | 1,01E-03 |
| **235** | LOC100133165 | 1,03 | 1,59 | 1,76E-06 | 3,35E-05 |
| **236** | LOC100507412 | -1,64 | 2,03 | 2,23E-09 | 9,12E-08 |
| **237** | LOC101593348 | 1,38 | -0,28 | 1,20E-04 | 1,19E-03 |
| **238** | LOC105377622 | 1,10 | 1,36 | 5,46E-05 | 6,14E-04 |
| **239** | LOC107985388 | 1,43 | 1,02 | 1,20E-05 | 1,70E-04 |
| **240** | LOC122526782 | -1,22 | 2,93 | 2,59E-06 | 4,60E-05 |
| **241** | LOC401261 | 1,24 | 1,27 | 5,69E-05 | 6,34E-04 |
| **242** | LOC730668 | 1,75 | 0,37 | 3,98E-07 | 9,14E-06 |
| **243** | LONRF3 | 1,35 | 3,77 | 7,65E-22 | 1,81E-19 |
| **244** | LRP2 | 1,21 | 4,02 | 3,05E-10 | 1,45E-08 |
| **245** | LRRC8E | 1,05 | 3,37 | 9,04E-08 | 2,45E-06 |
| **246** | LRRK2 | 1,05 | 2,28 | 1,51E-09 | 6,42E-08 |
| **247** | LRRTM4 | -7,81 | -1,30 | 2,78E-12 | 1,88E-10 |
| **248** | LRSAM1 | 1,02 | 3,11 | 1,01E-10 | 5,35E-09 |
| **249** | LSAMP | -2,12 | -0,35 | 1,06E-04 | 1,07E-03 |
| **250** | LTBP4 | 1,12 | 5,53 | 1,19E-09 | 5,12E-08 |
| **251** | MAGEB2 | 2,55 | 4,31 | 9,12E-65 | 1,58E-61 |
| **252** | MAL2 | -5,24 | 1,18 | 2,43E-44 | 2,92E-41 |
| **253** | MAMLD1 | 1,28 | 2,03 | 1,42E-10 | 7,25E-09 |
| **254** | MAP2 | -1,89 | 1,39 | 1,47E-16 | 2,02E-14 |
| **255** | MAP3K15 | 1,46 | 3,28 | 4,21E-26 | 1,73E-23 |
| **256** | MAP6 | -1,61 | 0,42 | 2,93E-05 | 3,67E-04 |
| **257** | MATK | 1,44 | 4,37 | 2,32E-07 | 5,63E-06 |
| **258** | MBP | 1,20 | 1,50 | 7,51E-06 | 1,13E-04 |
| **259** | MEA1 | -1,15 | 5,70 | 1,57E-06 | 3,02E-05 |
| **260** | MET | 1,04 | 6,61 | 9,31E-32 | 5,58E-29 |
| **261** | MFAP2 | -9,84 | 2,08 | 3,63E-99 | 1,42E-95 |
| **262** | MFSD13A | 1,01 | 3,98 | 1,20E-12 | 8,75E-11 |
| **263** | MGC16275 | 1,59 | 0,51 | 4,28E-04 | 3,41E-03 |
| **264** | MICA-AS1 | 2,91 | -0,30 | 6,91E-06 | 1,05E-04 |
| **265** | MIR3648-1 | 1,95 | 2,25 | 3,85E-04 | 3,11E-03 |
| **266** | MIR924HG | 1,88 | -0,69 | 1,04E-04 | 1,06E-03 |

| | | | | | |
|---|---|---|---|---|---|
| **267** | MKX | 1,81 | 3,04 | 1,01E-31 | 5,83E-29 |
| **268** | MLXIPL | 1,20 | 3,28 | 3,16E-13 | 2,58E-11 |
| **269** | MME | -1,70 | 2,89 | 1,74E-22 | 4,38E-20 |
| **270** | MORF4L1P1 | -1,14 | 3,46 | 1,32E-03 | 8,57E-03 |
| **271** | MRFAP1L1 | -1,10 | 4,36 | 9,98E-05 | 1,02E-03 |
| **272** | MRPL51 | -1,13 | 5,33 | 2,19E-04 | 1,98E-03 |
| **273** | MSL3P1 | -1,38 | 2,14 | 8,60E-10 | 3,80E-08 |
| **274** | MSS51 | 1,56 | 0,29 | 3,83E-04 | 3,11E-03 |
| **275** | MSX2 | -1,41 | 4,34 | 2,89E-29 | 1,50E-26 |
| **276** | MT1F | -2,40 | -1,06 | 9,33E-08 | 2,51E-06 |
| **277** | MT1X | -1,50 | 3,17 | 1,97E-12 | 1,37E-10 |
| **278** | MT2A | -1,88 | 3,96 | 9,69E-14 | 8,73E-12 |
| **279** | MYT1 | 1,58 | -0,44 | 1,98E-04 | 1,82E-03 |
| **280** | NCL | -1,56 | 9,48 | 6,13E-08 | 1,76E-06 |
| **281** | NEBL | 1,01 | 5,10 | 9,80E-25 | 3,32E-22 |
| **282** | NEFM | 1,93 | 9,65 | 1,15E-18 | 2,16E-16 |
| **283** | NFKBID | 1,20 | 0,67 | 4,78E-04 | 3,73E-03 |
| **284** | NGEF | -1,04 | 2,27 | 5,45E-05 | 6,14E-04 |
| **285** | NHS | 1,14 | 4,76 | 1,45E-10 | 7,35E-09 |
| **286** | NIBAN1 | 1,46 | 2,37 | 3,67E-13 | 2,95E-11 |
| **287** | NKX3-1 | -1,13 | 2,22 | 9,19E-05 | 9,51E-04 |
| **288** | NLRC5 | 1,38 | 2,53 | 4,70E-15 | 5,16E-13 |
| **289** | NMNAT2 | 1,85 | 0,48 | 2,93E-08 | 9,17E-07 |
| **290** | NOS2 | 1,66 | 1,78 | 4,68E-11 | 2,58E-09 |
| **291** | NOXA1 | 1,19 | 0,07 | 2,67E-04 | 2,32E-03 |
| **292** | NPR1 | -3,22 | -0,32 | 3,07E-09 | 1,24E-07 |
| **293** | NPR2 | 1,06 | 1,41 | 2,41E-05 | 3,12E-04 |
| **294** | NPTXR | -1,14 | 2,46 | 6,77E-08 | 1,92E-06 |
| **295** | NSA2 | -1,02 | 6,79 | 6,15E-04 | 4,60E-03 |
| **296** | NUTM2A | 1,32 | 2,04 | 7,94E-06 | 1,18E-04 |
| **297** | OPTN | 1,03 | 2,55 | 2,60E-04 | 2,27E-03 |
| **298** | OSMR | 1,37 | 2,14 | 7,25E-11 | 3,91E-09 |
| **299** | OSTN | -1,23 | 2,26 | 2,87E-08 | 9,01E-07 |
| **300** | OXCT1 | -2,06 | 2,96 | 3,06E-33 | 2,07E-30 |
| **301** | P2RX5 | 1,31 | 2,06 | 6,77E-10 | 3,06E-08 |
| **302** | P4HTM | -2,10 | 1,63 | 7,64E-10 | 3,41E-08 |
| **303** | PAK3 | -3,01 | 0,37 | 2,78E-12 | 1,88E-10 |
| **304** | PANX2 | 1,15 | 2,12 | 9,14E-08 | 2,47E-06 |
| **305** | PAOX | 1,26 | 0,62 | 3,45E-04 | 2,86E-03 |
| **306** | PAPPA | -4,64 | 0,54 | 6,35E-15 | 6,64E-13 |
| **307** | PARP12 | 1,24 | 2,87 | 2,56E-13 | 2,12E-11 |
| **308** | PAX1 | -6,41 | -1,60 | 9,24E-06 | 1,34E-04 |
| **309** | PAX7 | -1,29 | 0,20 | 8,62E-04 | 6,04E-03 |
| **310** | PCDHB2 | -3,61 | -1,08 | 2,86E-05 | 3,60E-04 |
| **311** | PCK2 | 1,42 | 4,11 | 1,11E-11 | 6,83E-10 |
| **312** | PCYT1B | 1,25 | 4,47 | 8,32E-23 | 2,13E-20 |

| | | | | | |
|---|---|---|---|---|---|
| **313** | PDE3A | -4,69 | -0,99 | 5,13E-08 | 1,52E-06 |
| **314** | PDE4B | -1,77 | -0,19 | 2,80E-06 | 4,91E-05 |
| **315** | PGR | 1,14 | 4,52 | 4,90E-16 | 6,21E-14 |
| **316** | PHEX | 1,04 | 0,95 | 1,48E-03 | 9,36E-03 |
| **317** | PIK3AP1 | 1,31 | 3,65 | 2,90E-24 | 8,86E-22 |
| **318** | PILRB | 1,55 | 1,23 | 3,25E-04 | 2,73E-03 |
| **319** | PIMREGP3 | 1,09 | 0,64 | 4,72E-05 | 5,46E-04 |
| **320** | PITRM1-AS1 | 1,35 | 0,79 | 2,27E-05 | 2,96E-04 |
| **321** | PKD1 | 1,09 | 5,73 | 6,83E-10 | 3,08E-08 |
| **322** | PKD1P3 | 3,52 | 0,77 | 8,80E-08 | 2,40E-06 |
| **323** | PLCL1 | 3,50 | 3,98 | 3,73E-154 | 5,81E-150 |
| **324** | PLEKHA2 | 1,04 | 2,49 | 1,85E-11 | 1,11E-09 |
| **325** | PLIN2 | 1,73 | 1,90 | 1,23E-14 | 1,23E-12 |
| **326** | PLPP4 | -2,85 | 0,60 | 4,08E-14 | 3,83E-12 |
| **327** | PLXDC2 | -6,90 | -0,12 | 4,10E-20 | 8,87E-18 |
| **328** | PNMA2 | 1,14 | 5,30 | 1,59E-21 | 3,69E-19 |
| **329** | POU3F2 | -1,01 | 4,48 | 9,80E-10 | 4,30E-08 |
| **330** | PPP1R1A | 1,14 | 0,34 | 4,46E-06 | 7,22E-05 |
| **331** | PPP1R26-AS1 | -1,64 | 1,07 | 2,23E-07 | 5,44E-06 |
| **332** | PRAME | 1,56 | 3,80 | 1,24E-14 | 1,23E-12 |
| **333** | PRDM6-AS1 | -1,34 | 1,52 | 2,38E-05 | 3,08E-04 |
| **334** | PRECSIT | 1,84 | -0,32 | 1,11E-03 | 7,44E-03 |
| **335** | PRKCB | -2,00 | 1,88 | 1,96E-20 | 4,31E-18 |
| **336** | PRKCQ | 1,02 | 2,18 | 1,29E-05 | 1,81E-04 |
| **337** | PRKCQ-AS1 | 1,12 | 2,10 | 9,23E-09 | 3,32E-07 |
| **338** | PRKG1 | 1,16 | 5,57 | 6,89E-25 | 2,39E-22 |
| **339** | PROSER2 | 1,13 | 4,03 | 8,77E-07 | 1,81E-05 |
| **340** | PRR18 | 1,39 | 1,00 | 5,76E-04 | 4,36E-03 |
| **341** | PSG4 | 1,13 | 1,36 | 8,48E-05 | 8,89E-04 |
| **342** | PTER | 1,00 | 4,26 | 5,40E-10 | 2,51E-08 |
| **343** | PTH1R | -1,24 | -0,10 | 1,22E-03 | 8,06E-03 |
| **344** | PTMA | -1,16 | 10,08 | 1,61E-07 | 4,11E-06 |
| **345** | PTMAP2 | -1,58 | 2,93 | 6,18E-04 | 4,62E-03 |
| **346** | PTMAP9 | -1,76 | 5,04 | 1,04E-04 | 1,06E-03 |
| **347** | PTN | -8,56 | -1,41 | 6,34E-09 | 2,36E-07 |
| **348** | PUDP | -9,02 | 2,84 | 1,98E-93 | 6,19E-90 |
| **349** | PWWP2B | 1,09 | 3,30 | 5,94E-04 | 4,47E-03 |
| **350** | RAB26 | 1,43 | 1,10 | 8,44E-06 | 1,24E-04 |
| **351** | RAB39A | 2,01 | 0,59 | 3,51E-06 | 5,91E-05 |
| **352** | RAMP1 | -1,38 | 3,16 | 1,00E-07 | 2,69E-06 |
| **353** | RASA4 | 1,26 | 3,32 | 1,88E-19 | 3,85E-17 |
| **354** | RASA4B | 1,46 | 2,82 | 2,50E-18 | 4,59E-16 |
| **355** | RASSF2 | 1,30 | 2,78 | 2,08E-09 | 8,57E-08 |
| **356** | RASSF4 | 1,56 | 1,17 | 3,92E-12 | 2,59E-10 |
| **357** | RELN | -5,36 | 0,31 | 7,46E-31 | 4,16E-28 |
| **358** | RELT | 1,05 | 3,96 | 3,94E-09 | 1,55E-07 |

| | | | | | |
|---|---|---|---|---|---|
| **359** | REPIN1-AS1 | 1,14 | 1,15 | 1,19E-04 | 1,18E-03 |
| **360** | RERG | 1,04 | 2,94 | 1,51E-09 | 6,42E-08 |
| **361** | RGL1 | -1,34 | 3,04 | 2,23E-23 | 6,07E-21 |
| **362** | RGS16 | 1,26 | 4,10 | 4,70E-09 | 1,81E-07 |
| **363** | RGS7BP | -3,65 | -0,57 | 7,43E-12 | 4,69E-10 |
| **364** | RHOF | 1,38 | 1,89 | 1,27E-07 | 3,30E-06 |
| **365** | RIMS3 | 1,27 | 4,10 | 1,16E-19 | 2,44E-17 |
| **366** | RIN2 | 1,23 | 1,26 | 2,65E-05 | 3,38E-04 |
| **367** | RNA45SN1 | 1,91 | 7,08 | 1,88E-09 | 7,81E-08 |
| **368** | RNA45SN2 | 2,17 | 10,31 | 1,83E-12 | 1,28E-10 |
| **369** | RNA45SN3 | 1,70 | 2,82 | 1,45E-04 | 1,40E-03 |
| **370** | RND3 | -1,08 | 5,23 | 1,96E-20 | 4,31E-18 |
| **371** | RNF122 | 1,10 | 5,60 | 6,51E-16 | 8,05E-14 |
| **372** | RNF180 | -3,99 | -0,39 | 1,13E-08 | 3,94E-07 |
| **373** | RNU1-3 | -1,44 | 6,73 | 1,43E-05 | 1,97E-04 |
| **374** | RNVU1-18 | -1,44 | 6,73 | 1,43E-05 | 1,97E-04 |
| **375** | ROR2 | -1,53 | 0,64 | 1,27E-04 | 1,24E-03 |
| **376** | RPL11 | -1,47 | 8,58 | 9,20E-04 | 6,38E-03 |
| **377** | RPL21P75 | -1,09 | 2,59 | 4,12E-06 | 6,75E-05 |
| **378** | RPL22P1 | -1,67 | 5,30 | 6,58E-04 | 4,85E-03 |
| **379** | RPL26 | -1,02 | 9,15 | 2,84E-05 | 3,58E-04 |
| **380** | RPL27A | -1,15 | 8,31 | 1,25E-03 | 8,16E-03 |
| **381** | RPL27AP6 | -1,29 | 4,93 | 2,73E-04 | 2,36E-03 |
| **382** | RPL36AP37 | -3,10 | 5,60 | 9,19E-04 | 6,37E-03 |
| **383** | RPL37 | -1,24 | 8,78 | 3,37E-04 | 2,81E-03 |
| **384** | RPL37A | -1,29 | 8,81 | 1,76E-07 | 4,42E-06 |
| **385** | RPL38 | -1,59 | 7,96 | 5,01E-04 | 3,88E-03 |
| **386** | RPRM | -1,41 | 2,46 | 2,82E-11 | 1,65E-09 |
| **387** | RPS26 | -1,08 | 6,31 | 9,97E-04 | 6,82E-03 |
| **388** | RPS26P7 | -2,36 | -0,27 | 3,11E-04 | 2,63E-03 |
| **389** | RPS6KL1 | 1,00 | 3,58 | 2,58E-17 | 4,28E-15 |
| **390** | RTN1 | -1,46 | 3,11 | 5,59E-15 | 5,97E-13 |
| **391** | RYR2 | -1,28 | 3,70 | 6,95E-10 | 3,12E-08 |
| **392** | SBK1 | 1,17 | 2,50 | 2,80E-08 | 8,85E-07 |
| **393** | SCARA5 | 1,24 | 0,72 | 2,80E-06 | 4,91E-05 |
| **394** | SCARNA7 | -3,71 | 3,31 | 7,19E-04 | 5,23E-03 |
| **395** | SCIN | -2,04 | 0,17 | 1,02E-09 | 4,48E-08 |
| **396** | SCLY | -1,16 | 5,21 | 1,20E-25 | 4,57E-23 |
| **397** | SEC11C | -1,29 | 3,61 | 1,58E-03 | 9,88E-03 |
| **398** | SELENOP | -1,36 | 4,79 | 1,49E-24 | 4,84E-22 |
| **399** | SEMA3A | 1,22 | 3,09 | 4,77E-17 | 7,29E-15 |
| **400** | SEMA6D | 1,19 | 2,36 | 1,59E-08 | 5,36E-07 |
| **401** | SERHL2 | 1,04 | 1,51 | 1,38E-04 | 1,34E-03 |
| **402** | SERPINF1 | -2,08 | 1,52 | 9,57E-24 | 2,71E-21 |
| **403** | SFRP1 | -5,75 | 1,93 | 1,03E-69 | 2,28E-66 |
| **404** | SFRP2 | 3,73 | 2,69 | 4,74E-50 | 6,71E-47 |

| | | | | | |
|---|---|---|---|---|---|
| **405** | SFXN3 | 1,21 | 4,18 | 2,43E-27 | 1,11E-24 |
| **406** | SGK2 | 1,52 | 0,16 | 1,09E-04 | 1,09E-03 |
| **407** | SH2B2 | 1,38 | 2,42 | 1,17E-10 | 6,12E-09 |
| **408** | SH2D3C | 1,61 | 0,91 | 1,65E-10 | 8,17E-09 |
| **409** | SHC3 | 2,23 | 0,31 | 1,51E-08 | 5,12E-07 |
| **410** | SHLD2 | 1,11 | 6,00 | 1,34E-26 | 5,95E-24 |
| **411** | SLC10A3 | 1,09 | 3,18 | 2,03E-07 | 5,03E-06 |
| **412** | SLC10A4 | 1,47 | 1,91 | 7,72E-06 | 1,15E-04 |
| **413** | SLC10A5 | 1,05 | 0,53 | 4,80E-04 | 3,75E-03 |
| **414** | SLC13A3 | 1,87 | 1,17 | 1,56E-12 | 1,11E-10 |
| **415** | SLC19A3 | -6,63 | -0,58 | 1,48E-07 | 3,81E-06 |
| **416** | SLC22A17 | -1,05 | 1,61 | 4,88E-04 | 3,80E-03 |
| **417** | SLC25A21-AS1 | 1,07 | 0,20 | 2,52E-04 | 2,21E-03 |
| **418** | SLC30A3 | 1,14 | 1,03 | 2,83E-06 | 4,94E-05 |
| **419** | SLC45A3 | 1,13 | 2,17 | 3,77E-04 | 3,07E-03 |
| **420** | SLC6A11 | 1,06 | 5,26 | 1,69E-25 | 6,12E-23 |
| **421** | SLC6A9 | 1,35 | 3,66 | 4,27E-11 | 2,37E-09 |
| **422** | SLC7A5P2 | 1,13 | 2,78 | 6,53E-09 | 2,41E-07 |
| **423** | SLC9A5 | 1,20 | 1,03 | 7,09E-06 | 1,08E-04 |
| **424** | SLCO3A1 | -2,47 | -0,56 | 4,54E-09 | 1,75E-07 |
| **425** | SLCO5A1 | 1,10 | 3,74 | 1,58E-08 | 5,36E-07 |
| **426** | SLFN12 | -9,59 | 0,33 | 1,18E-40 | 1,32E-37 |
| **427** | SLIRP | -1,45 | 4,54 | 1,41E-04 | 1,36E-03 |
| **428** | SLIT1 | 1,41 | 3,64 | 3,33E-17 | 5,30E-15 |
| **429** | SLTM | -1,05 | 4,96 | 1,13E-03 | 7,54E-03 |
| **430** | SLX9 | -1,09 | 3,99 | 2,22E-08 | 7,27E-07 |
| **431** | SMAD6 | -1,84 | 2,97 | 4,45E-23 | 1,18E-20 |
| **432** | SMARCA1 | -2,47 | 4,32 | 3,02E-66 | 5,88E-63 |
| **433** | SNAI2 | -2,07 | 4,06 | 2,50E-26 | 1,06E-23 |
| **434** | SNRPF | -1,39 | 4,75 | 1,07E-03 | 7,24E-03 |
| **435** | SOX11 | -4,71 | 0,16 | 9,75E-16 | 1,20E-13 |
| **436** | SOX21 | 1,58 | 1,32 | 6,32E-05 | 6,91E-04 |
| **437** | SP6 | 1,76 | 0,02 | 2,35E-04 | 2,10E-03 |
| **438** | SPAG7 | -1,33 | 3,10 | 4,93E-05 | 5,66E-04 |
| **439** | SPARC | -8,62 | 2,59 | 5,26E-111 | 2,73E-107 |
| **440** | SPHK1 | 1,05 | 2,11 | 1,85E-04 | 1,71E-03 |
| **441** | SPIRE2 | 1,03 | 2,78 | 6,34E-07 | 1,37E-05 |
| **442** | SPNS2 | 1,16 | 1,58 | 2,71E-05 | 3,43E-04 |
| **443** | SPOCD1 | 1,42 | -1,13 | 1,18E-03 | 7,80E-03 |
| **444** | SPOCK1 | -4,77 | 0,07 | 5,64E-29 | 2,83E-26 |
| **445** | SPRN | 1,55 | 0,77 | 2,82E-06 | 4,93E-05 |
| **446** | SPTBN4 | 1,14 | 0,83 | 7,22E-04 | 5,25E-03 |
| **447** | SRRM3 | 1,10 | 1,10 | 1,19E-04 | 1,18E-03 |
| **448** | SSUH2 | 1,20 | -0,72 | 1,13E-03 | 7,55E-03 |
| **449** | STAM-DT | 1,52 | -0,51 | 9,51E-05 | 9,80E-04 |
| **450** | STARD13 | 1,07 | 2,73 | 6,16E-10 | 2,80E-08 |

| | | | | | |
|---|---|---|---|---|---|
| **451** | STING1 | -1,49 | 2,29 | 3,13E-12 | 2,09E-10 |
| **452** | STK16P1 | 1,50 | -0,04 | 2,47E-04 | 2,19E-03 |
| **453** | STS | -11,79 | 3,18 | 3,27E-133 | 2,55E-129 |
| **454** | SULT1A1 | -1,07 | 0,57 | 1,44E-03 | 9,15E-03 |
| **455** | SULT1C4 | -5,06 | 0,17 | 3,37E-18 | 6,03E-16 |
| **456** | SYBU | 1,10 | 2,91 | 4,42E-10 | 2,07E-08 |
| **457** | SYN3 | 1,32 | 2,23 | 2,71E-11 | 1,59E-09 |
| **458** | SYNDIG1 | -7,79 | -1,03 | 1,71E-09 | 7,15E-08 |
| **459** | TAC1 | 1,10 | 1,75 | 6,20E-06 | 9,55E-05 |
| **460** | TAL1 | 1,13 | 0,97 | 9,48E-05 | 9,78E-04 |
| **461** | TEDC2-AS1 | 1,40 | -0,49 | 1,28E-03 | 8,32E-03 |
| **462** | TEPSIN | 1,02 | 3,38 | 1,25E-05 | 1,75E-04 |
| **463** | TFEB | 1,33 | 1,49 | 2,05E-07 | 5,06E-06 |
| **464** | TFPI2 | -1,15 | 1,59 | 3,29E-05 | 4,07E-04 |
| **465** | TIGD1 | -1,01 | 3,46 | 6,05E-05 | 6,67E-04 |
| **466** | TINCR | 1,15 | 0,90 | 6,48E-04 | 4,79E-03 |
| **467** | TIPARP-AS1 | 1,46 | -0,06 | 1,77E-05 | 2,39E-04 |
| **468** | TLCD3B | 1,11 | 0,63 | 1,45E-04 | 1,39E-03 |
| **469** | TMC5 | 1,01 | 0,91 | 5,39E-04 | 4,13E-03 |
| **470** | TMCC3 | 1,05 | 2,74 | 7,54E-08 | 2,10E-06 |
| **471** | TMEM102 | 1,50 | 1,65 | 6,62E-06 | 1,01E-04 |
| **472** | TMEM132E | 1,41 | 0,97 | 8,34E-05 | 8,77E-04 |
| **473** | TMEM200B | 2,49 | -0,57 | 3,46E-05 | 4,22E-04 |
| **474** | TMEM266 | 1,06 | 1,51 | 4,57E-05 | 5,33E-04 |
| **475** | TNC | 1,31 | 9,63 | 5,18E-33 | 3,36E-30 |
| **476** | TNFSF9 | 2,36 | 0,87 | 1,20E-07 | 3,14E-06 |
| **477** | TNXB | 1,50 | 1,83 | 1,40E-08 | 4,81E-07 |
| **478** | TOMM40P2 | 2,27 | 1,61 | 4,14E-04 | 3,32E-03 |
| **479** | TRAF3IP1 | -1,22 | 4,18 | 5,67E-08 | 1,65E-06 |
| **480** | TRAPPC14 | 1,26 | 3,62 | 1,12E-14 | 1,13E-12 |
| **481** | TRIM6 | 1,40 | 1,41 | 4,01E-07 | 9,16E-06 |
| **482** | TRIP6 | 1,01 | 7,05 | 1,24E-08 | 4,30E-07 |
| **483** | TRPC6 | 1,93 | -0,09 | 3,22E-06 | 5,49E-05 |
| **484** | TYW1B | 1,15 | 1,67 | 1,11E-03 | 7,44E-03 |
| **485** | U2AF1 | -1,65 | 5,82 | 1,73E-04 | 1,62E-03 |
| **486** | UBE2L6 | 2,37 | -0,55 | 5,60E-05 | 6,25E-04 |
| **487** | UBE2QL1 | -2,44 | 0,63 | 8,90E-13 | 6,63E-11 |
| **488** | UGT3A2 | -1,38 | 1,85 | 5,49E-09 | 2,08E-07 |
| **489** | UNCX | -2,72 | -1,16 | 1,56E-04 | 1,49E-03 |
| **490** | USP32P2 | -2,19 | 0,33 | 1,55E-07 | 3,98E-06 |
| **491** | USP40 | -1,12 | 4,19 | 4,00E-18 | 7,01E-16 |
| **492** | VAX2 | -1,18 | 0,95 | 3,39E-04 | 2,82E-03 |
| **493** | WDFY1 | -1,15 | 5,90 | 8,77E-28 | 4,14E-25 |
| **494** | WNT8B | 1,06 | 0,29 | 1,22E-04 | 1,20E-03 |
| **495** | WSCD1 | 1,58 | 4,67 | 8,42E-24 | 2,43E-21 |
| **496** | WWC3 | -1,76 | 4,13 | 2,37E-39 | 2,17E-36 |

| | | | | | |
|---|---|---|---|---|---|
| **497** | XK | 1,26 | 3,75 | 5,71E-19 | 1,09E-16 |
| **498** | YBX1 | -1,03 | 10,16 | 1,23E-03 | 8,06E-03 |
| **499** | ZC3H12A | 1,03 | 2,40 | 5,67E-07 | 1,24E-05 |
| **500** | ZC3H12D | 1,29 | 2,23 | 1,33E-05 | 1,86E-04 |
| **501** | ZCCHC12 | -2,21 | 2,93 | 1,20E-15 | 1,46E-13 |
| **502** | ZNF155 | -1,25 | 2,56 | 5,09E-09 | 1,94E-07 |
| **503** | ZNF22 | -5,89 | 1,08 | 6,28E-39 | 5,44E-36 |
| **504** | ZNF222 | -1,01 | 1,22 | 3,45E-04 | 2,86E-03 |
| **505** | ZNF223 | -1,29 | 1,49 | 5,50E-08 | 1,61E-06 |
| **506** | ZNF229 | -1,24 | 2,57 | 5,97E-08 | 1,72E-06 |
| **507** | ZNF365 | 1,22 | 2,20 | 1,98E-06 | 3,69E-05 |
| **508** | ZNF417 | -1,13 | 3,49 | 3,97E-12 | 2,61E-10 |
| **509** | ZNF43 | -7,58 | 0,58 | 2,35E-40 | 2,44E-37 |
| **510** | ZNF438 | 1,10 | 1,93 | 5,35E-08 | 1,57E-06 |
| **511** | ZNF525 | -1,20 | 2,88 | 4,52E-13 | 3,57E-11 |
| **512** | ZNF595 | -3,42 | -0,69 | 8,37E-06 | 1,23E-04 |
| **513** | ZNF785 | 1,93 | 0,76 | 3,15E-09 | 1,26E-07 |
| **514** | ZNF883 | -1,56 | 3,05 | 4,96E-13 | 3,84E-11 |
| **515** | ZSWIM8 | 1,06 | 5,62 | 1,09E-12 | 8,05E-11 |
| **Homozygous *vs.* heterozygous E475G cell line** | | | | | |
| | **Gene** | **logFC** | **logCPM** | **PValue** | **FDR** |
| **1** | ADAM12 | -3,76 | -0,42 | 8,90E-08 | 1,33E-05 |
| **2** | ADAMTS16 | -3,27 | 1,48 | 4,38E-26 | 9,75E-23 |
| **3** | AMBN | -1,33 | 1,14 | 1,30E-05 | 9,45E-04 |
| **4** | ARC | -1,08 | 2,26 | 1,10E-06 | 1,16E-04 |
| **5** | ATP2B3 | -1,75 | 2,09 | 1,57E-12 | 5,55E-10 |
| **6** | ATP8A2 | -1,36 | 2,16 | 1,60E-10 | 3,83E-08 |
| **7** | C8orf34 | 1,33 | 1,06 | 1,40E-04 | 6,17E-03 |
| **8** | CA2 | -1,15 | 6,37 | 5,75E-11 | 1,57E-08 |
| **9** | CASP10 | 1,35 | 0,80 | 1,33E-04 | 5,99E-03 |
| **10** | CCN2 | -1,48 | 4,18 | 3,60E-21 | 4,31E-18 |
| **11** | CCN3 | 1,05 | 2,12 | 2,69E-07 | 3,61E-05 |
| **12** | CDH20 | 1,86 | -0,61 | 2,72E-04 | 9,77E-03 |
| **13** | CDH23 | -1,07 | 2,50 | 1,16E-10 | 2,96E-08 |
| **14** | CHRDL1 | -5,51 | -0,68 | 8,77E-05 | 4,41E-03 |
| **15** | CLSTN2 | -1,24 | 1,47 | 1,42E-05 | 1,01E-03 |
| **16** | COL12A1 | -1,84 | 2,36 | 3,27E-11 | 9,27E-09 |
| **17** | CRLF1 | -1,41 | 0,02 | 1,61E-04 | 6,82E-03 |
| **18** | CRTAC1 | -1,20 | 0,53 | 1,44E-04 | 6,27E-03 |
| **19** | CSRNP1 | -1,40 | 4,16 | 4,86E-18 | 3,61E-15 |
| **20** | DTX3 | -1,62 | 0,84 | 2,23E-06 | 2,12E-04 |
| **21** | EGFR | 1,22 | 4,59 | 1,44E-20 | 1,60E-17 |
| **22** | ELL3 | 1,12 | 1,33 | 1,36E-04 | 6,10E-03 |
| **23** | EMILIN2 | -1,00 | 3,84 | 3,22E-17 | 2,18E-14 |
| **24** | ERBB3 | 1,28 | 1,11 | 9,52E-08 | 1,41E-05 |
| **25** | ESRRG | 1,18 | 2,42 | 4,82E-10 | 1,07E-07 |

| | | | | | |
|---|---|---|---|---|---|
| **26** | ETS1 | -1,52 | 1,63 | 2,14E-11 | 6,53E-09 |
| **27** | ETV1 | 1,42 | 2,95 | 4,52E-16 | 2,82E-13 |
| **28** | ETV4 | 1,99 | 3,51 | 4,74E-27 | 1,48E-23 |
| **29** | ETV5 | 1,15 | 4,73 | 2,62E-24 | 3,72E-21 |
| **30** | FAAH2 | -1,45 | 0,52 | 2,68E-05 | 1,71E-03 |
| **31** | FAM83B | 1,09 | 2,28 | 3,26E-05 | 2,05E-03 |
| **32** | FBXL7 | -2,02 | 0,82 | 2,62E-05 | 1,68E-03 |
| **33** | FGF13 | -2,85 | 1,02 | 6,57E-16 | 3,66E-13 |
| **34** | FLI1 | -1,55 | 1,54 | 3,30E-07 | 4,35E-05 |
| **35** | FOSL2 | -1,09 | 4,59 | 6,52E-25 | 1,13E-21 |
| **36** | FOXF1 | -1,03 | 2,05 | 4,71E-05 | 2,78E-03 |
| **37** | FOXN4 | 1,32 | 1,00 | 6,43E-08 | 1,01E-05 |
| **38** | GABRE | -2,41 | -0,08 | 2,06E-07 | 2,79E-05 |
| **39** | GALNT14 | -1,59 | 0,59 | 4,14E-06 | 3,57E-04 |
| **40** | GALNT3 | 1,23 | 3,23 | 4,04E-12 | 1,34E-09 |
| **41** | GPR37 | 1,14 | 2,47 | 2,30E-06 | 2,17E-04 |
| **42** | GPR50 | -2,04 | 1,39 | 4,27E-07 | 5,28E-05 |
| **43** | GRIK5 | -1,83 | -0,36 | 1,33E-05 | 9,63E-04 |
| **44** | HMGN2P3 | -1,00 | 3,20 | 4,49E-05 | 2,70E-03 |
| **45** | HMOX1 | -1,29 | 3,44 | 8,35E-10 | 1,78E-07 |
| **46** | HNF4G | 1,13 | 2,35 | 1,80E-05 | 1,23E-03 |
| **47** | HOXC4 | 1,33 | 2,80 | 7,64E-06 | 6,04E-04 |
| **48** | HSPA6 | -3,75 | -0,87 | 8,48E-07 | 9,37E-05 |
| **49** | IFI44L | 1,71 | -0,10 | 3,89E-05 | 2,37E-03 |
| **50** | IGF2 | -3,83 | 3,99 | 8,17E-41 | 4,25E-37 |
| **51** | IGFBP7 | -1,54 | 1,56 | 1,09E-09 | 2,24E-07 |
| **52** | IKZF3 | 1,00 | 3,91 | 2,09E-13 | 9,57E-11 |
| **53** | IL10RB | -1,05 | 3,05 | 1,93E-04 | 7,61E-03 |
| **54** | INSM1 | 2,50 | 0,34 | 7,57E-07 | 8,49E-05 |
| **55** | ITPR3 | 1,28 | 4,68 | 1,40E-24 | 2,19E-21 |
| **56** | KIT | 1,44 | 3,88 | 5,74E-25 | 1,12E-21 |
| **57** | LAPTM5 | -5,20 | -0,56 | 7,41E-20 | 7,22E-17 |
| **58** | LOC100101148 | -6,51 | -1,26 | 8,53E-06 | 6,65E-04 |
| **59** | MAL2 | -2,73 | 1,18 | 5,08E-09 | 9,77E-07 |
| **60** | MAP6 | -1,97 | 0,42 | 3,36E-07 | 4,41E-05 |
| **61** | MDGA2 | 1,07 | 2,36 | 1,10E-07 | 1,61E-05 |
| **62** | MFAP2 | -5,10 | 2,08 | 2,95E-11 | 8,69E-09 |
| **63** | MOCOS | 1,00 | 1,74 | 1,02E-04 | 4,88E-03 |
| **64** | MYLIP | 1,00 | 3,33 | 2,21E-08 | 3,87E-06 |
| **65** | MYOM2 | 1,35 | 4,84 | 3,13E-38 | 1,22E-34 |
| **66** | NIBAN1 | 1,07 | 2,37 | 1,41E-07 | 1,96E-05 |
| **67** | NUP153-AS1 | 1,06 | 0,17 | 6,43E-05 | 3,59E-03 |
| **68** | OLMALINC | 1,08 | 2,08 | 4,54E-05 | 2,71E-03 |
| **69** | PAX7 | -1,47 | 0,20 | 1,91E-04 | 7,55E-03 |
| **70** | PLEKHA2 | 1,07 | 2,49 | 3,48E-11 | 9,68E-09 |
| **71** | PLPP4 | -1,69 | 0,60 | 2,59E-05 | 1,67E-03 |

| | | | | | |
|---|---|---|---|---|---|
| **72** | PLXDC2 | -4,65 | -0,12 | 4,36E-07 | 5,31E-05 |
| **73** | PPP1R1A | 1,16 | 0,34 | 1,02E-05 | 7,62E-04 |
| **74** | PRKCB | -1,56 | 1,88 | 2,95E-12 | 1,00E-09 |
| **75** | PSG4 | 1,53 | 1,36 | 5,66E-07 | 6,67E-05 |
| **76** | PUDP | -9,05 | 2,84 | 5,51E-96 | 4,29E-92 |
| **77** | RELN | -2,73 | 0,31 | 1,90E-06 | 1,84E-04 |
| **78** | RGS2 | 1,14 | 3,60 | 7,60E-13 | 2,89E-10 |
| **79** | RGS7BP | -2,52 | -0,57 | 1,18E-04 | 5,44E-03 |
| **80** | RIMS3 | 1,06 | 4,10 | 5,77E-14 | 2,81E-11 |
| **81** | RN7SKP71 | -3,09 | 3,10 | 2,17E-04 | 8,27E-03 |
| **82** | RN7SL4P | -2,31 | 5,52 | 5,45E-05 | 3,12E-03 |
| **83** | RN7SL5P | -3,84 | 3,21 | 4,96E-05 | 2,89E-03 |
| **84** | RNA45SN1 | 1,49 | 7,08 | 2,21E-06 | 2,12E-04 |
| **85** | RNA45SN2 | 1,79 | 10,31 | 3,88E-09 | 7,66E-07 |
| **86** | RNA45SN3 | 1,98 | 2,82 | 1,27E-05 | 9,25E-04 |
| **87** | RNF122 | 1,11 | 5,60 | 7,43E-16 | 3,99E-13 |
| **88** | RNU1-3 | -1,24 | 6,73 | 1,70E-04 | 6,99E-03 |
| **89** | RNVU1-18 | -1,24 | 6,73 | 1,70E-04 | 6,99E-03 |
| **90** | RPS4XP6 | -2,59 | 0,01 | 2,13E-04 | 8,12E-03 |
| **91** | SCN3B | -1,29 | 0,65 | 4,61E-05 | 2,73E-03 |
| **92** | SEMA3A | 1,22 | 3,09 | 1,58E-15 | 8,23E-13 |
| **93** | SEMA6D | 1,00 | 2,36 | 3,19E-06 | 2,85E-04 |
| **94** | SFRP1 | -2,92 | 1,93 | 4,28E-14 | 2,15E-11 |
| **95** | SFRP2 | 1,34 | 2,69 | 7,69E-10 | 1,66E-07 |
| **96** | SHLD2P3 | 1,44 | 1,05 | 2,24E-04 | 8,47E-03 |
| **97** | SLC25A21-AS1 | 1,45 | 0,20 | 1,39E-05 | 9,97E-04 |
| **98** | SLC37A1 | -1,29 | 1,77 | 6,69E-06 | 5,37E-04 |
| **99** | SLFN12 | -6,13 | 0,33 | 1,49E-07 | 2,06E-05 |
| **100** | SP8 | 2,27 | 1,27 | 8,94E-05 | 4,48E-03 |
| **101** | SPARC | -3,96 | 2,59 | 5,01E-13 | 2,08E-10 |
| **102** | SPOCK1 | -2,72 | 0,07 | 8,51E-08 | 1,29E-05 |
| **103** | SPRY1 | 1,04 | 4,28 | 2,72E-13 | 1,21E-10 |
| **104** | SPRY4 | 1,51 | 1,09 | 2,81E-04 | 9,95E-03 |
| **105** | STAT6 | 1,01 | 4,18 | 2,75E-17 | 1,95E-14 |
| **106** | STK16P1 | 1,73 | -0,04 | 9,04E-05 | 4,51E-03 |
| **107** | STS | -11,78 | 3,18 | 1,05E-145 | 1,63E-141 |
| **108** | SYBU | 1,12 | 2,91 | 1,75E-09 | 3,55E-07 |
| **109** | SYNDIG1 | -5,74 | -1,03 | 1,45E-04 | 6,27E-03 |
| **110** | TFPI2 | -1,72 | 1,59 | 6,64E-10 | 1,46E-07 |
| **111** | USP41 | 2,80 | -0,54 | 4,87E-05 | 2,85E-03 |
| **112** | VGF | -1,59 | 1,33 | 1,96E-12 | 6,78E-10 |
| **113** | WWC3 | -1,27 | 4,13 | 3,03E-20 | 3,14E-17 |
| **114** | ZCCHC12 | -2,82 | 2,93 | 6,21E-24 | 8,06E-21 |
| **115** | ZNF22 | -2,48 | 1,08 | 2,79E-04 | 9,91E-03 |
| **116** | ZNF43 | -6,08 | 0,58 | 3,45E-19 | 3,16E-16 |
| **117** | ZNF785 | 1,23 | 0,76 | 1,82E-04 | 7,30E-03 |
