## Supplementary Table S3 for "Pathological PNPase variants with altered RNA binding and degradation activity affect the phenotype of bacterial and human cell models"

**Supplementary Table S3. mtRNA relative amount**

| | Replicate | Treatment[a] | Read number | Reads mapped on mtDNA | % reads mapped on mtDNA | % Overall Alignment rate | Reads mapped on nuclear DNA | % mapped on mt on total mapped reads |
|---|---|---|---|---|---|---|---|---|
| WT | 1 | NT | 40029427 | 185474 | 0.46 | 0.50 | 22646816 | 0.81 |
| | 2 | | 56587765 | 301193 | 0.53 | 0.57 | 35692289 | 0.84 |
| | 3 | | 57640935 | 648986 | 1.13 | 1.31 | 34872480 | 1.83 |
| | 1 | STS | 71838957 | 619309 | 0.86 | 0.92 | 53029318 | 1.15 |
| | 2 | | 69365863 | 662112 | 0.95 | 1.07 | 50104425 | 1.30 |
| | 3 | | 72968012 | 5354426 | 7.34 | 8.30 | 39322215 | 11.98 |
| p.E475G heterozygous cells | 1 | NT | 42626416 | 719725 | 1.69 | 1.84 | 25322850 | 2.76 |
| | 2 | | 50987934 | 300919 | 0.59 | 0.65 | 34790712 | 0.86 |
| | 3 | | 69263486 | 714180 | 1.03 | 1.19 | 34008468 | 2.06 |
| | 1 | STS | 35182032 | 370529 | 1.05 | 1.18 | 22896840 | 1.59 |
| | 2 | | 34811331 | 257734 | 0.74 | 0.91 | 24079352 | 1.06 |
| | 3 | | 59566472 | 569659 | 0.96 | 1.05 | 39129125 | 1.43 |
| p.E475G homozygous cells | 1 | NT | 59830192 | 239562 | 0.40 | 0.46 | 36393427 | 0.65 |
| | 2 | | 65691076 | 390101 | 0.59 | 0.65 | 41425942 | 0.93 |
| | 3 | | 51397615 | 285983 | 0.56 | 0.59 | 25763098 | 1.10 |
| | 1 | STS | 45518323 | 599101 | 1.32 | 1.46 | 27962048 | 2.10 |
| | 2 | | 51906820 | 356144 | 0.69 | 0.74 | 32071953 | 1.10 |
| | 3 | | 32917426 | 309617 | 0.94 | 0.99 | 20792371 | 1.47 |

[a]NT, not treated; STS, incubated with staurosporine
