## Supplementary Table S4 for "Pathological PNPase variants with altered RNA binding and degradation activity affect the phenotype of bacterial and human cell models"

**Supplementary Table S4. Apoptosis pathway enrichment**

| | Regulation of Apoptotic Process (GO:0042981)[a] | |
|---|---|---|
| | p-value[b] | q-value[b] |
| 293T[b] | 3.39E-08 | 5.30E-06 |
| p.E475G he[b] | 1.13E-07 | 2.40E-05 |
| p.E475G ho[b] | 5.59E-10 | 1.65E-07 |

[a]Gene Ontology (GO) analysis for overrepresented gene sets among DEGs with FDR > 0.01 in the pairwise comparisons between samples of RNA extracted was performed from the indicated cells untreated and treated with staurosporine.

[b]Values calculated with Enrichr (37) using the Benjamini-Hochberg method for correction for multiple hypotheses testing.
